# Systematic revision of the suborder Astrophorina (Porifera: Demospongiae) in the temperate Northeast Pacific

**DOI:** 10.64898/2026.06.22.733819

**Authors:** Thomas L. Turner

## Abstract

This study presents a systematic revision of the suborder Astrophorina for the temperate Pacific coast of the United States and Canada. Major findings include a reduction in the number of species previously thought to range into the region from Japan; validation of most *Geodia* species erected by Lendenfeld (1910), which were later synonymized by de Laubenfels (1932); the formal description of 10 new species (*Poecillastra alaskensis* sp. nov., *Vulcanella explorata* sp. nov., *Vulcanella rupta* sp. nov., *Stelletta cardenasi* sp. nov., *Stelletta nicolenya* sp. nov., *Stelletta limuwensis* sp. nov., *Dercitus (Stoeba) giveni* sp. nov., *Penares anyapax* sp. nov., *Penares foxi* sp. nov., and *Thenea diastra* sp. nov.); and one new combination, *Penares orientalis* comb. nov. Extensive SCUBA-based collection efforts yielded new samples for 11 of the 26 species identified in the region, which enabled an integrative taxonomic approach that combined field photography, fresh material for DNA sequencing, and improved characterization of species ranges and morphological variability in previously described taxa. Illumina sequencing generated complete nuclear ribosomal haplotypes for five species, while Sanger sequencing of the 28S and cox1 loci placed 20 of the 26 species within molecular phylogenies. The use of very short “mini-barcode” amplicons also enabled sequence recovery from historic type specimens up to 137 years old. This study additionally reports the discovery of sponge grounds of abundant, large *Geodia* at diving depths in Southern California. Together, these results substantially advance our understanding of global astrophorid diversity and systematics, and the biogeography of sponge diversity in the Northeast Pacific.

Note about species names: this pre-print is not intended to be a publication of the associated species names for the purposes of zoological nomenclature.

## Introduction

The suborder Astrophorina is a diverse group of demosponges, with over 1,000 described species found throughout the world’s oceans (Díaz *et al*. 2024; de Voogd, *et al*. 2025). They can be abundant and diverse at a wide variety of depths, but are best known from deep waters, where they form ecologically important "sponge grounds" in the North Atlantic and Mediterranean (Klitgaard & Tendal 2004; Maldonado *et al*. 2015). Similar dense aggregations of large astrophorid sponges have not been described from the North Pacific, but astrophorid sponges have been the focus of less research effort in this region. Recent work in the Aleutian Islands (Alaska) has revealed animal forests and sponge grounds that are among the most diverse and ecologically important deep-water habitats in the world (Beckmann *et al*. 2026; Stone 2014). Astrophorid sponges are an integral part of these habitats, with five new astrophorid species described from deep waters around the Aleutian Islands in the past 20 years (Lehnert *et al*. 2006, 2014; Lehnert & Stone 2014), and four new species described from seamounts in the nearby Northwestern Pacific (Shilov *et al*. 2023). In contrast, the remainder of the US and Canadian temperate Pacific region has seen no newly described astrophorid species in nearly a century (since 1930). Astrophorid sponges are seen regularly in deep-water ROV surveys in the region (Beckmann *et al*. 2026; Graiff & Lipski 2023; Lindholm *et al*. 2013), and I have encountered fields of large *Geodia* as shallow as 25 m off California (see results below), but identifying these sponges has proven difficult based on past literature. In the case of *Geodia*, there are eight species described from the region, all from Lendenfeld (1910). These sponges were described from material collected between 1888 and 1904 by the first purpose-built scientific research vessel, the USS Albatross (Lendenfeld 1910). Though this work was very meticulous and detailed, it was later called into question by de Laubenfels (1932). Lendenfeld split the *Geodia* samples from the Albatross into taxa based on what could be perceived as minor differences: including named varieties, Lendenfeld described 14 taxa of *Geodia* in the region, the majority of which were represented by only a single sample. In his later monograph on the sponges of California, de Laubenfels (1930, 1932) reduced these 14 taxa to only three, lumping all *Geodia* with mesotriaene spicules under the name *G. mesotriaena* Lendenfeld, 1910. This was done despite the fact that de Laubenfels did not reexamine any of Lendenfeld’s samples. Instead, this decision was based on what de Laubenfels perceived as insufficient differences in Lendenfeld’s descriptions. It also seems that de Laubenfels harbored doubts about the accuracy of Lendenfeld’s work in general, stating that inaccuracies had been found by others, and that "we can only surmise the true status of his species pending a re-examination of the material by some competent investigator" (de Laubenfels, 1932 p. 24). Later authors apparently found the decision to lump most of these species to be premature, as all of Lendenfeld’s species (but not varieties) are considered valid in the World Porifera Database at the time of this writing (de Voogd, *et al*. 2025). However, because de Laubenfels’ monograph was used as a foundational reference for most later work in the North Pacific, I know of no further references to the species that he doubted, except by Lee *et al*. (2007), who stated that "the status of California geodias is not satisfactory and needs intensive review". Clearly, further work on this group is long overdue.

The status of *Poecillastra* in the region is likewise confused and uncertain. The type species for the genus, *P. compressa* (Bowerbank 1866), is described from the North Atlantic, but North Pacific sponges are quite similar. Some historical assessments postulated that *P. compressa* is cosmopolitan and found throughout the Atlantic, Pacific, and Indian Oceans (Burton 1930; Maldonado 2002). This assessment was rejected by Maldonado (2002), who noted that some species previously lumped with *P. compressa* possess differently sized asters than described for the type species. One of these species is *P. tenuilaminaris* (Sollas 1886), described from Sagami Bay, Japan. This species was therefore considered valid by Maldonado, but he also surmised that the California species *P. rickettsi* de Laubenfels, 1930 was a junior synonym of the Japanese *P. tenuilaminaris*. As a result, some later authors considered *P. tenuilaminaris* to be the only *Poecillastra* expected in the Northeast Pacific, and many samples have been assigned to that name. Later work disagreed, and pointed to the long, hair-like ectosomal oxeas of *P. rickettsi* as a character differentiating the California species from *P. tenuilaminaris* (Carvalho *et al*. 2011). Following this work, *P. rickettsi* is now considered a valid species in the World Porifera Database (de Voogd, *et al*. 2025). When describing *P. rickettsi*, de Laubenfels (1930, 1932) also reported *P. tenuilaminaris* from California, so both species may be present in the region. In contrast to these assessments, Koltun (1966) described all *Poecillastra* from the North Pacific as belonging to *P. japonica* (Thiele 1898). This species, also described from near Sagami Bay, Japan, is said to have two classes of oxeas like *P. rickettsi*, but both classes are thicker, and the long oxeas are not as long as those described from *P. rickettsi*. Koltun reported that the spicules of North Pacific *Poecillastra* were highly variable, but he could find no clear evidence of multiple species among them. Noting their similarity to *P. compressa*, he made *P. japonica* a subspecies of *P. compressa*, and stated that all North Pacific samples were *P. compressa japonica* and all North Atlantic and Arctic samples were *P. compressa compressa* (*P. japonica* is once again at species rank in the World Porifera Database). It is unclear why Koltun thought the North Pacific samples he examined were *P. japonica*, but he was presumably motivated by finding very thick primary oxeas and a second category of long and thick oxeas, which are the two characteristics that differentiate the original description of *P. japonica* from *P. tenuilaminaris* and *P. rickettsi*. However, Koltun also described the asters in his *P. compressa japonica* as 14–70 μm, which is in stark contrast to the type description of 15–20 μm. Though Koltun said the largest asters were sometimes absent, he did not specify when or where this was the case. As a result, *Poecillastra* from the Northeast Pacific have variously been assigned to *P. compressa, P. japonica, P. tenuilaminaris,* and *P. rickettsi*, and a comprehensive review is sorely needed.

In addition to *Geodia*, *Poecillastra*, and the recently described Aleutian species, there are five astrophorids described in the region, all from de Laubenfels (1930). These species, allocated to the genera *Stelletta*, *Penares*, and *Dercitus*, are known from shallow waters, but very little follow-up work has been done on them. I have repeatedly failed to obtain DNA from museum vouchers originally collected by de Laubenfels (Turner *et al*. 2024, 2025; Turner 2026; Turner & Pankey 2023). Consequently, performing integrative taxonomy on these species requires including more recently collected material. Moreover, these species were all collected before the advent of scientific diving, mostly from the intertidal zone, with the shallow subtidal zone being poorly sampled for taxonomic work in sponges.

To address these challenges, I have undertaken a revision of the suborder Astrophorina for the region. I have attempted to be inclusive of the area from the Aleutian Islands, Alaska, to Southern California (figure 1). Within this region, I have attempted to obtain new morphological and genetic data from all described species, except for those recently described from the Aleutian Islands (Lehnert *et al*. 2006, 2014; Lehnert & Stone 2014). This involved extensive collections in shallow waters, mostly on SCUBA, ranging from Sitka, Alaska to San Diego, California. I also examined many museum vouchers, including reexamining the type samples from most species. I did not, however, attempt to examine all previously collected museum vouchers that have been assigned to the Astrophorina. As a result, there are likely additional species which have already been collected that await description, and there are undoubtedly more species waiting to be collected for the first time. However, by using modern integrative taxonomy on an extensive new and historic collection of samples, I hope to establish a firm foundation on which future research efforts in this group can be built. Together with a recent revision of the family Tetillidae (Turner 2026), this work creates a jumping-off point for future work for the entire order Tetractinellida for the US and Canadian Pacific coast.

**Figure 1.**
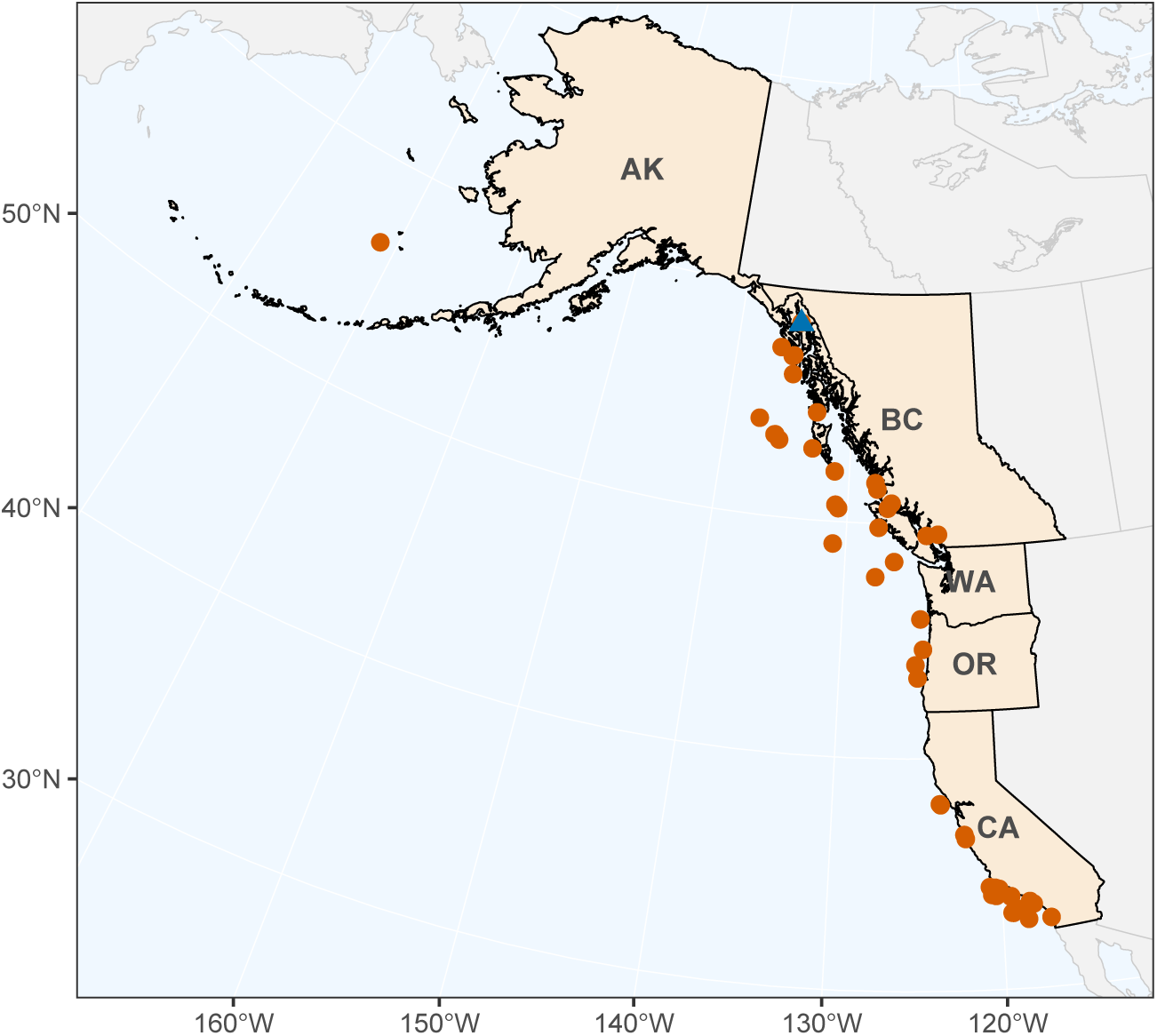
Map of the study region. The Pacific coasts of US states of California (CA), Oregon (OR), Washington (WA), and Alaska (AK), and the Canadian province of British Columbia (BC), were included. Orange points indicate the collection locations for samples previously collected by others and acquired on loan for reanalysis here. The blue triangle indicates where samples were collected by the author near Juneau, AK; for author-collected samples in California, see figure 2.

## Materials & Methods

### Collections

Subtidal collections on SCUBA were conducted by the author at 128 sites from San Diego, California, USA to Juneau, Alaska, USA, between 2019 to 2026. These collections were primarily in Southern and Central California, with additional sampling at five sites in Washington, USA, four in British Columbia, Canada, and five in Alaska, USA. Intertidal collections were made at 15 additional sites across the same range, again concentrated in California. In total, samples from 50 astrophorid sponges were collected: 42 from California (figure 2), eight from Alaska (figure 1), and none from Washington and British Columbia. Megan Lily (City of San Diego) contributed an additional sample, collected during the Southern California Bight 2023 project. Floating docks in 23 harbors and marinas were surveyed, but no astrophorids were found at these sites.

**Figure 2.**
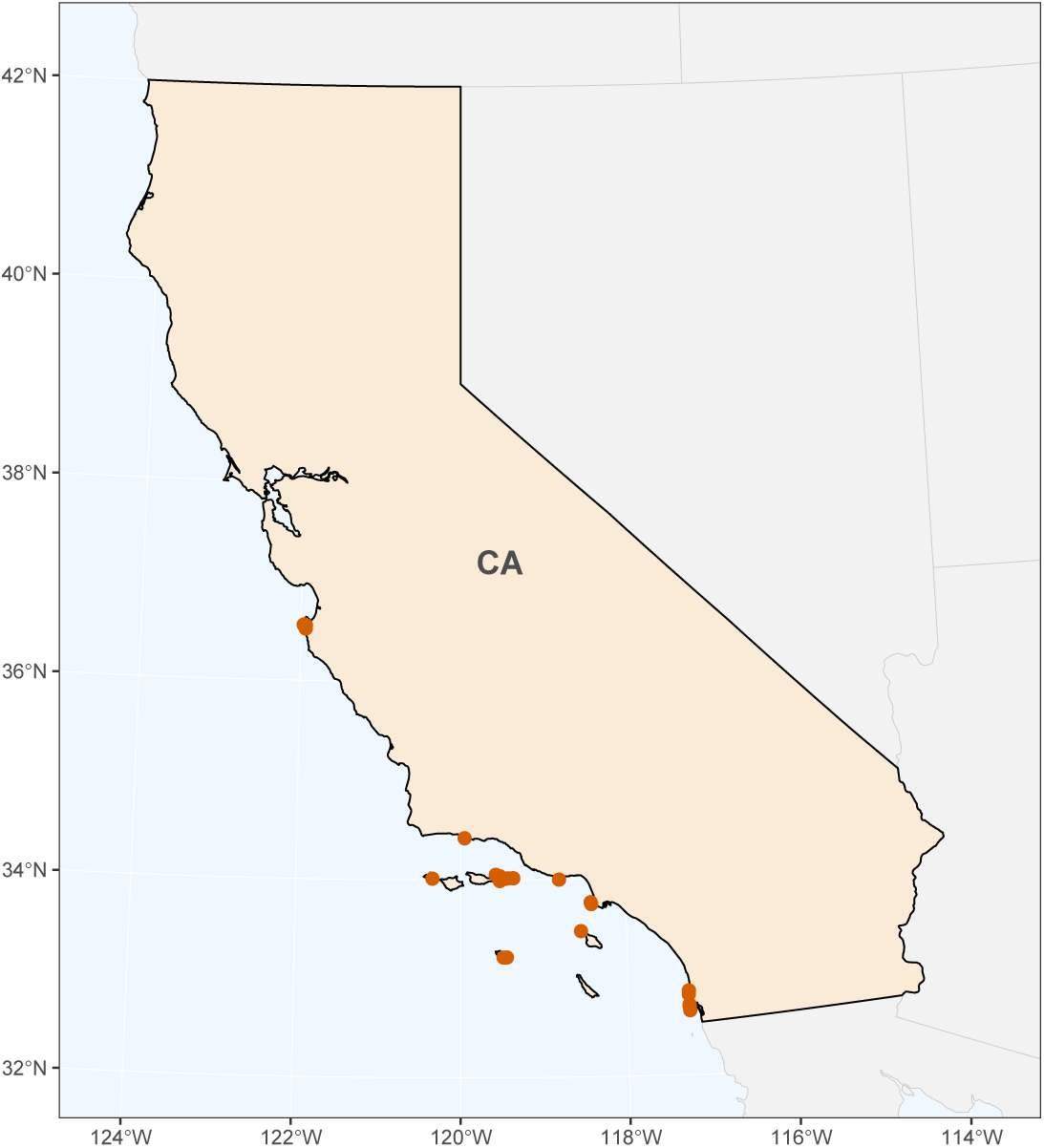
Map of samples collected by the author in California (CA). The most northern points are in Central California, around the Monterey Peninsula, while all other points are within the warm temperate Southern California Bight marine biogeographic province.

An additional 71 samples were acquired on loan from the following museum collections (figure 2): the Smithsonian National Museum of Natural History (USNM), the California Academy of Sciences (CASIZ), the Royal British Columbia Museum (RBC), and the Santa Barbara Museum of Natural History (SBMNH). Newly collected material was vouchered at these institutions, and also at the Cheadle Center at the University of California, Santa Barbara (UCSB), the Florida Museum of Natural History (BULA), and the Natural History Museum of Los Angeles (NHMLA). Voucher numbers are listed in the systematics section, tabulated in Table 4, and included in an expanded Table S1 archived at Data Dryad (doi.org/10.5061/dryad.z08kprrwf). The expanded table includes additional metadata including field numbers, GenBank accession numbers, collection locations and depths, iNaturalist links, and other notes.

### Morphology

Skeletal morphology was investigated using hand-cut tissue sections digested in 97% Nuclei Lysis Solution (Promega, Wizard Genomic DNA Purification Kit) and 3% Proteinase K (20 mg/ml, Promega). Cut sections were examined before digestion, and imaged immediately if sufficiently transparent. If not, they were placed on a coverslip, and the premixed solution was dripped on the section. The solution dissolves the organic material in the sample, but does not dissolve spongin. Skeletal features that are not bound by spongin will disassociate completely, so images of these features are taken during digestion, as the tissue becomes clear enough to image, but before digestion is completed.

Individual spicules were isolated by digesting sponge subsamples in bleach. Light images were taken with a Nikon D3500 SLR and an Amscope NDPL-1 microscope adaptor; scanning electron micrographs (SEMs) were taken with a FEI Quanta400F Mk2 after coating spicules with 20 nm of carbon. Measurements were made in ImageJ (Schneider *et al*. 2012). For simple spicules (e.g., oxeas), length was measured as the longest possible straight line from tip to tip, even when curved. Very long spicules are an exception, as they were measured in several straight intervals across multiple photos using landmarks. Spicule width was measured at the widest point, excluding adornments like tyles. Spicules like anatriaenes and mesoprotriaenes have a complex morphology, and additional measurements were made on each. Anatriaenes: rhabd length (excluding clads), rhabd width (widest point near the clad end), clad length (tip of clad to top of spicule), clad width (measured where the clad meets the rhabd), and clad angle (the angle from clad tip to the top of the spicule to center of rhabd). Mesoprotriaenes: rhabd length and width as above, longest clad length (tip of clad to rhabd junction), epirhabd length (same as clad lengths, but for central spire), clad width (widest point), and clad angle (180° minus the angle from rhabd center to clad junction to clad tip). A visual guide to these measurements is provided in the supporting information of Turner (2026).

For statistical comparisons, t-tests (*t.test*), Pearson correlations (*cor.test*), and principal components analysis (*prcomp*) were performed in R. Measurements are reported as minimum–mean–maximum with *n* equal to the number of spicules measured (which may differ across traits due to spicules with broken portions, or cases where only some measurements were made on a spicule). All 24,420 spicule measurements made are provided in the supporting table S2 archived at Data Dryad.

### Illumina sequencing

DNA was extracted using the Qiagen Blood & Tissue kit, treated with RNase A, and repurified with the Zymo DNA Clean & Concentrator. Library preparation and sequencing were performed at the UC Davis DNA Technologies Core Facility using the Super-High-Throughput (SHT) protocol, with dual indexing of 96 pooled samples. Six species of Atrophorida were included: *G. mesotriaena*, *S. cardenasi* sp. nov., *S. estrella, S. limuwensis* sp. nov., and two individuals each of *G. angulata* and *S. clarella*, alongside other sponges analyzed elsewhere (Turner *et al*. 2025; Turner 2026). Libraries were sequenced on an Illumina NovaSeq, generating paired-end 150 bp reads with a median of 2.1 million reads per sample (range: 208–17 million). This variability likely reflected differences in DNA preservation and secondary chemistry. Reads were trimmed with fastP (Chen *et al*. 2018) and assembled de novo with Megahit (Li *et al*. 2015). Assemblies yielded 0–1,080 contigs per sample (median: 22). Blast searches indicated that many contigs were microbial, or occasionally from fish, mollusks, or other taxa, but the nuclear ribosomal locus was typically the highest-coverage contig. Nearly complete ribosomal loci were assembled for all eight astrophorid sponges. These contigs were then used as references: reads were aligned with bwa-mem (Li & Durbin 2009), and consensus sequences were generated with SAMtools (Danecek *et al*. 2021). This served as the final sequence for that sample.

### Sanger sequencing

I attempted to sequence the Folmer region of cox1 and two adjacent amplicons of the nuclear ribosomal 28S locus (the D1D2 and D3D5 regions). These loci were chosen because they have lots of available comparative data and a reasonable rate of evolution for species discrimination. However, pan-poriferan primers previously published by others (Folmer *et al*. 1994; Morrow *et al*. 2012; Rot *et al*. 2006) performed poorly in astrophorids, except for the D3D5 region of 28S, which amplified consistently. Using Illumina assemblies generated here and data from GenBank, I designed eight new primer sets, which are provided in the supporting information archived at Data Dryad. These primer sets were variable in length, in order to gather longer sequences in well preserved samples while still obtaining short sequences from degraded samples. Amplicons 650–700 bp in length performed well in most recently collected material, but often failed in older samples. Additional primer sets targeting regions of variable size, down to as small as 150 bp "mini-barcode" regions (Cárdenas & Moore 2019) were developed, and sequentially smaller amplicons were attempted in important samples. Sequences newly generated here varied greatly in length, from full ribosomal loci 7113 bp long to <100 bp sequences amplified from old or degraded samples. These sequences have been deposited in GenBank, except for amplicons rejected by the archive due to short length. These sequences are provided in the supplementary materials. Accession numbers are listed in Table 4 and in the supporting table S1 archived at data dryad.

### Phylogenetic methods

Astrophorid sequences were compiled from GenBank via BLAST searches. For the cox1 locus, sequences were included if they minimally encompassed the Folmer region of cox1, but full mitochondrial genomes were used when available (available for five species). For the 28S locus, sequences were included if they minimally encompassed the C2D2 barcoding region, but longer sequences up to the full ribosomal locus were included. The final dataset includes data from numerous previous studies (Cárdenas 2020; Cárdenas *et al*. 2010, 2011; Despujols 2021; Díaz *et al*. 2024; Erpenbeck *et al*. 2015; Kelly *et al*. 2019; Lavrov *et al*. 2008, 2023; Lim *et al*. 2017; Ngwakum *et al*. 2021; Nichols 2005; Nunley *et al*. 2025; Plese *et al*. 2021; Shilov *et al*. 2023; Steffen *et al*. 2023; Thacker *et al*. 2013; Williams *et al*. 2024; Zeng *et al*. 2016). Alignments were generated in MAFFT v.7 (Katoh *et al*. 2017). Phylogenies were inferred with maximum likelihood in IQ-TREE version 3.1.1 (Nguyen *et al*. 2015; Trifinopoulos *et al*. 2016), with nodal support assessed using ultrafast bootstrapping and SH-aLRT tests (Hoang *et al*. 2018). Optimal substitution models were selected with ModelFinder, as implemented in IQ-TREE (Kalyaanamoorthy *et al*. 2017); TVM+F+I+R2 was selected for cox1, and TN+F+I+G4 was selected for 28S.

## Results

### Molecular phylogenies

Figures 3 and 4 show the maximum likelihood phylogeny of the Astrophorina, including all species for which data are available for both mitochondrial and nuclear ribosomal loci. Single-locus trees include many more species and samples, and are provided in the supplementary materials archived at Data Dryad (doi.org/10.5061/dryad.z08kprrwf). Single-locus trees include five more species from the study region: *Stelletta limuwensis* sp. nov., *Thenea diastra* sp. nov., and *Poecillastra alaskensis* sp. nov. are only represented in the 28S tree, and *Geodia carolae* (Lendenfeld, 1910) and *Penares saccharis* (de Laubenfels 1930) are only represented in the cox1 tree. DNA sequencing was unsuccessful for four previously described species (*Geodia ovis* Lendenfeld, 1910*, G. mesotriaenella* (Lendenfeld, 1910)*, G. bicolor* (Lendenfeld 1910), and *Dercitus syrmatitus* de Laubenfels, 1930, and one newly described species, *Dercitus giveni* sp. nov. In addition to Sanger sequencing, Illumina sequencing was performed on some recently collected samples. Metagenomic assembly of contigs from these samples successfully generated nearly-complete (∼7,000 bp) ribosomal loci from four *Stelletta* species and two *Geodia* species. These are included in figures 3 and 4, except for *Stelletta limuwensis* sp. nov., which is included in the single-locus 28S tree only.

**Figure 3.**
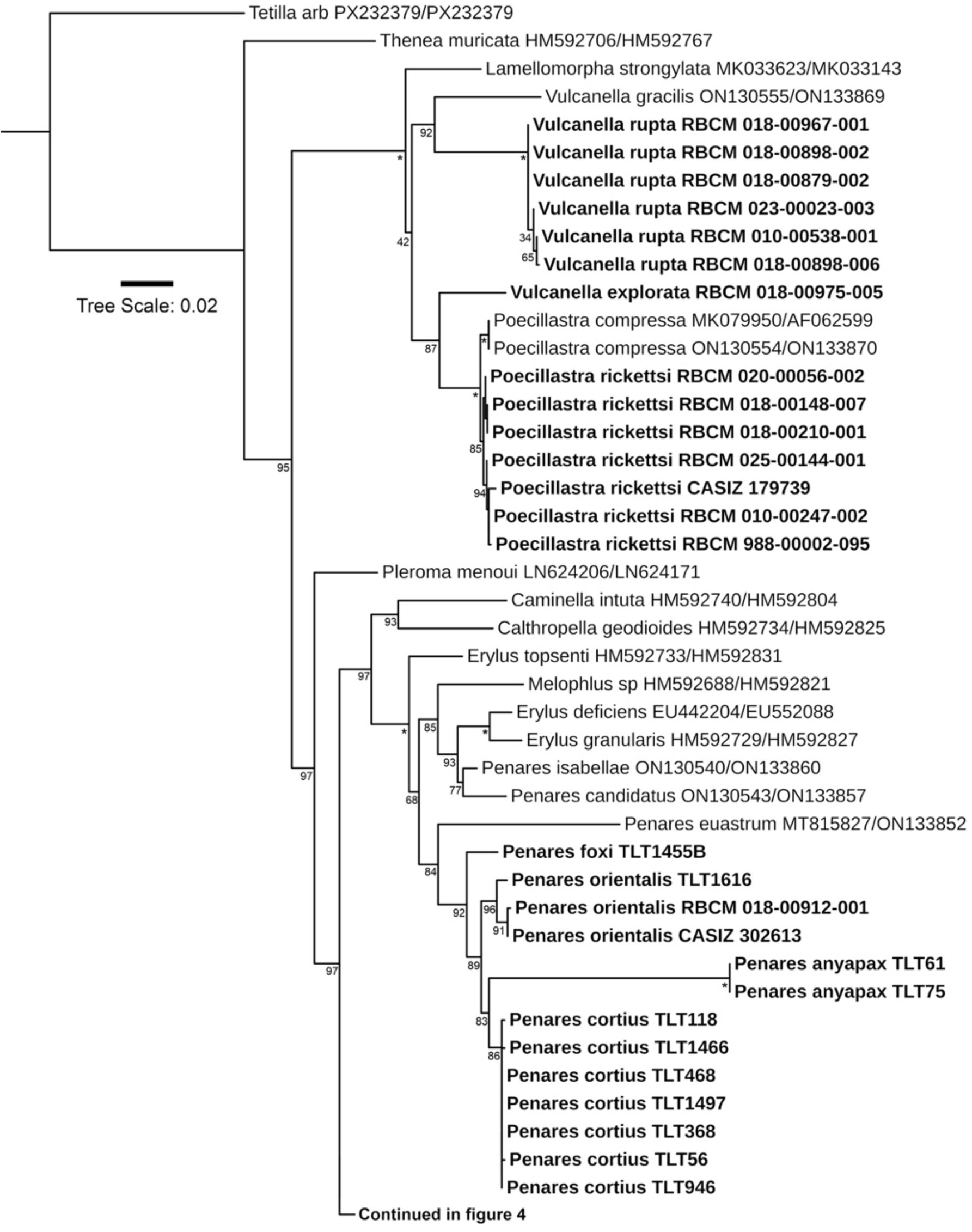
Maximum likelihood phylogeny of the Astrophorina, part 1. Mitochondrial and ribosomal sequences were concatenated and partitioned; only species with data from both partitions are included. New sequences are shown in bold, along with the sample name. Data mined from GenBank includes accession numbers. Node confidence is indicated with bootstrap values; asterisks indicate nodes ≥ 98% confidence. Scale bar indicates substitutions per site.

**Figure 4.**
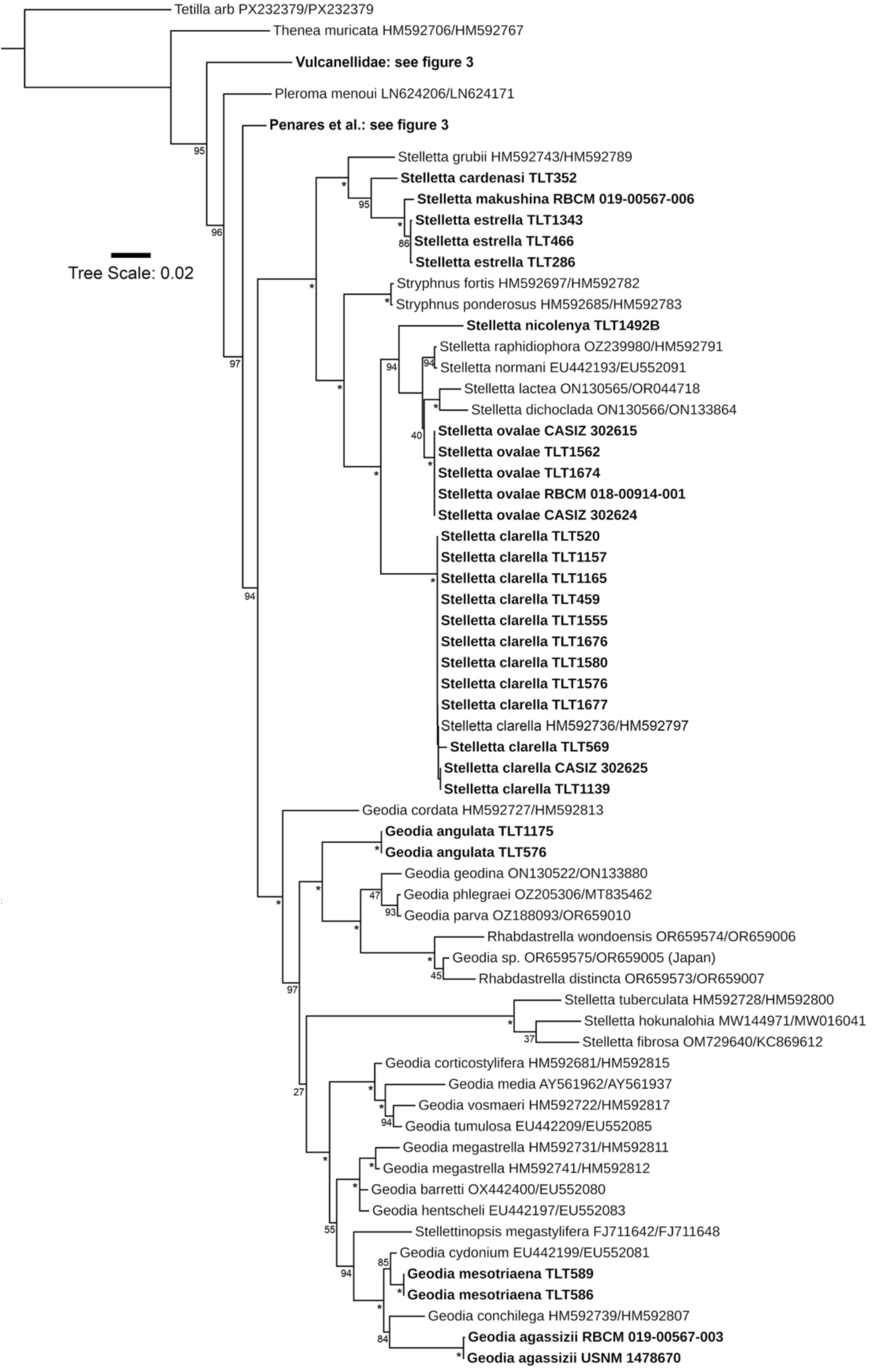
Maximum likelihood phylogeny of the Astrophorina, part 2. Mitochondrial and ribosomal sequences were concatenated and partitioned; only species with data from both partitions is included. New sequences are shown in bold, along with the sample name. Data mined from GenBank includes accession numbers. Node confidence is indicated with bootstrap values; asterisks indicate nodes ≥ 98% confidence. Scale bar indicates substitutions per site.

The phylogeny generated here is broadly consistent with previous phylogenies for the order, including the finding that many genera (*Stelletta*, *Geodia*, *Penares,* etc.) are polyphyletic (Cárdenas 2020; Cárdenas *et al*. 2011; Díaz *et al*. 2024; Schuster *et al*. 2015, 2018). Interestingly, all Northeast Pacific species of *Penares* formed an exclusive clade relative to other sequenced species, with the exception of *P. saccharis* (de Laubenfels 1930). *P. saccharis* is the only species of *Penares* in the region that lacks asters. As a result, it was originally assigned to the genus *Papyrula* Schmidt, 1868, which was later synonymized with *Penares* (Uriz 2002a). The absence of asters alone appears to be a weak character for generic delimitation. However, it is noteworthy that all *Penares* species lacking asters, and for which sequence data are available, form an exclusive clade. *P. candidatus* (Schmidt 1868), the former type species of *Papyrula*, is the only such species with both cox1 and 28S sequence data and is therefore the only representative included in Figure 3. In the *cox1* phylogeny, *P. saccharis* is sister to *P. candidatus*. Together, they form a clade with *P. sphaera* (Lendenfeld 1907) and *P. alatus* (Lendenfeld 1907). All of these species were originally assigned to *Papyrula* except *P. alatus*, which has plagiotriaenes rather than dichotriaenes and was therefore placed in a different family altogether. Additional *Penares* species lacking asters are known, but to my knowledge, none currently have available DNA data. Although it remains possible that asters have been lost independently multiple times within this group, the current evidence suggests that *Papyrula* may represent a natural lineage. Regardless of which characters are ultimately used in future taxonomic revisions, it seems likely that *Papyrula* will be resurrected to accommodate *P. saccharis* and its close relatives.

The single-locus phylogenies also include additional samples of species represented in figures 3 and 4. Some of these were successful only for small "mini-barcode" (100–200 bp) regions of a single locus, but these were useful in increasing confidence when assigning names to specific clades. For example, a ∼135 bp region of cox1 was successfully sequenced from many of Lendenfeld’s *Geodia* type samples, collected between 1888 and 1904. This includes the holotypes of all three named varieties of *G. mesotriaena*, two of three varieties of *G. angulata*, and two *G. agassizii* syntypes. Unfortunately, DNA sequencing was unsuccessful for many historical samples as well, with no DNA amplified from the types of *G. carolae, G. ovis, G. mesotriaenella, G. breviana,* or *G. bicolor.* In contrast to the partial success of DNA sequencing in *Geodia* samples collected by the Albatross, I was unable to amplify DNA from any of de Laubenfels’ type samples, which were collected in California in the 1920s. This is consistent with previous failed attempts to sequence DNA from de Laubenfels’ California material in other sponge taxa (Turner *et al*. 2024, 2025; Turner 2026; Turner & Pankey 2023).

My primary purpose in constructing the phylogeny was to use the inferred patterns of DNA similarity and divergence to aid in taxonomic inference and species delimitation. In this regard, the DNA data was indispensable. In the case of *Geodia*, comparing the short cox1 sequences from the *G. agassizii* and *G. mesotriaena* type samples revealed that they are quite different (divergence (Dxy) = 8.7%; Fst = 1.0) and verify that at least some of Lendenfeld’s *Geodia* species — later doubted by de Laubenfels — are truly distinct species. I was then able to obtain longer sequences of cox1 and 28S from more recent vouchers, and use these to determine the phylogenetic position of these species. Though *G. agassizii* and *G. mesotriaena* are both within the Cydonium^P^ clade (sensu Cárdenas *et al*., 2011), they fall into well-differentiated subclades within it (figure 4). They are each more closely related to species from the Atlantic and Mediterranean than to each other, with a common ancestor estimated to have lived in the early Cretaceous (Rossi *et al*. 2026).

In the family Vulcanellidae, species of *Vulcanella* were not previously reported from the Northeast Pacific, but genetic data supports two new species from deep water seamounts off British Columbia. These samples are described below as *V. rupta* sp. nov. and *V. explorata* sp. nov. *Poecillastra* is also in this family, but in this case, the sequence data are somewhat equivocal regarding where to draw species boundaries. I attempted to obtain genetic data from 21 *Poecillastra* samples, and was successful in 13 cases. These data were consistent with a single species, with some reservations. Two genotypes were seen at the cox1 locus. These differ at only a few base pairs (Dxy=0.010, Fst = 0.75), but this is a similar level of divergence as seen when these genotypes are compared to other species. When they are each compared to samples identified as *P. compressa* from the North Atlantic and Mediterranean, Dxy=0.007–0.012 and Fst=0.89–0.93, and when they are each compared to an unidentified species from South Africa, Dxy=0.005–0.010 and Fst=0.84–0.92. In contrast to data at cox1, however, no differences were seen between these mitochondrial clades within the North Pacific at the 28S locus. This differs when samples are compared across regions: North Pacific samples and North Atlantic *P. compressa* samples are divergent at 28S (Dxy=0.045, Fst=0.90). These data are therefore consistent with either a single species among the North Pacific samples, or two species that are too closely related to confirm without more genetic data. I also note that a complete mitochondrial genome assembly for one of these samples (RBCM 025-00144-001) was generated by Kristen Westfall (Fisheries and Oceans Canada), and is published here for the first time. When aligned to the mitogenome previously published for a *P. compressa* from Norway, divergence (Dxy) is 0.0072 (Plese *et al*. 2021). As discussed in the systematic section, these samples were all morphologically assigned to the species *P. rickettsi*, except for two, which were assigned to the new species *P. alaskensis* sp. nov. (only a small region of 28S was successfully sequenced for this new species, as discussed in the systematics section).

DNA sequences were also critical in delineating new species in the genera *Stelletta* and *Penares* that were very morphologically similar. *Stelletta cardenasi* sp. nov., *S. limuwensis* sp. nov., *S. nicolenya* sp. nov., and *S. estrella* are all morphologically distinguishable, but given the high variability in spicule dimensions seen in some *Stelletta* species, they could easily have been lumped together without genetic support. In the single-locus gene trees, which include many more species than the partitioned tree in figure 4, each of these species is more closely related to *Stelletta* from distant regions than they are to each other. Similarly, there was an unexpectedly high diversity of *Penares* in the region, with five species found where only two were previously known. Unlike *Stelletta*, four of these species form a clade exclusive of all other species with sequence data available. Genetic data was very helpful in establishing species concepts within this clade, because there were four well-differentiated (2%–4% sequence divergence) clades that were found at both the 28S and cox1 loci. Because the mitochondrial and nuclear genomes are unlinked, and because the most closely related genotypes in this clade are sympatric, divergence at both loci is strong evidence of reproductive isolation in nature (Jennings, 1917; Rannala & Yang, 2020). There was weaker evidence for a sixth species, as one sample of *P. orientalis* comb. nov. was distinct at both loci (0.8% divergence at 28S, 0.4% divergence at cox1) and morphologically distinguishable. However, as these genotypes were sister groups, and there is only one sample of this genotype, determining if there are additional reproductively isolated species will await further data.

### Biogeography

The results reported in the systematics section, combined with my recent revision of the family Tetillidae (Turner 2026), revise the entire order Tetractinellida for the US and Canadian temperate Pacific coasts. Figure 5 shows a summary of species level diversity in the region, and how species ranges have changed compared to previous knowledge. Overall, the number of species known in the region has increased by over 50%, from 29 species to 46 species. This is primarily due to the 18 newly described species, but also includes two species not previously known from the region (though in the case of *Tetilla japonica* (Lampe 1886), presence in the Northeast Pacific remains tentative (Turner 2026)).

**Figure 5.**
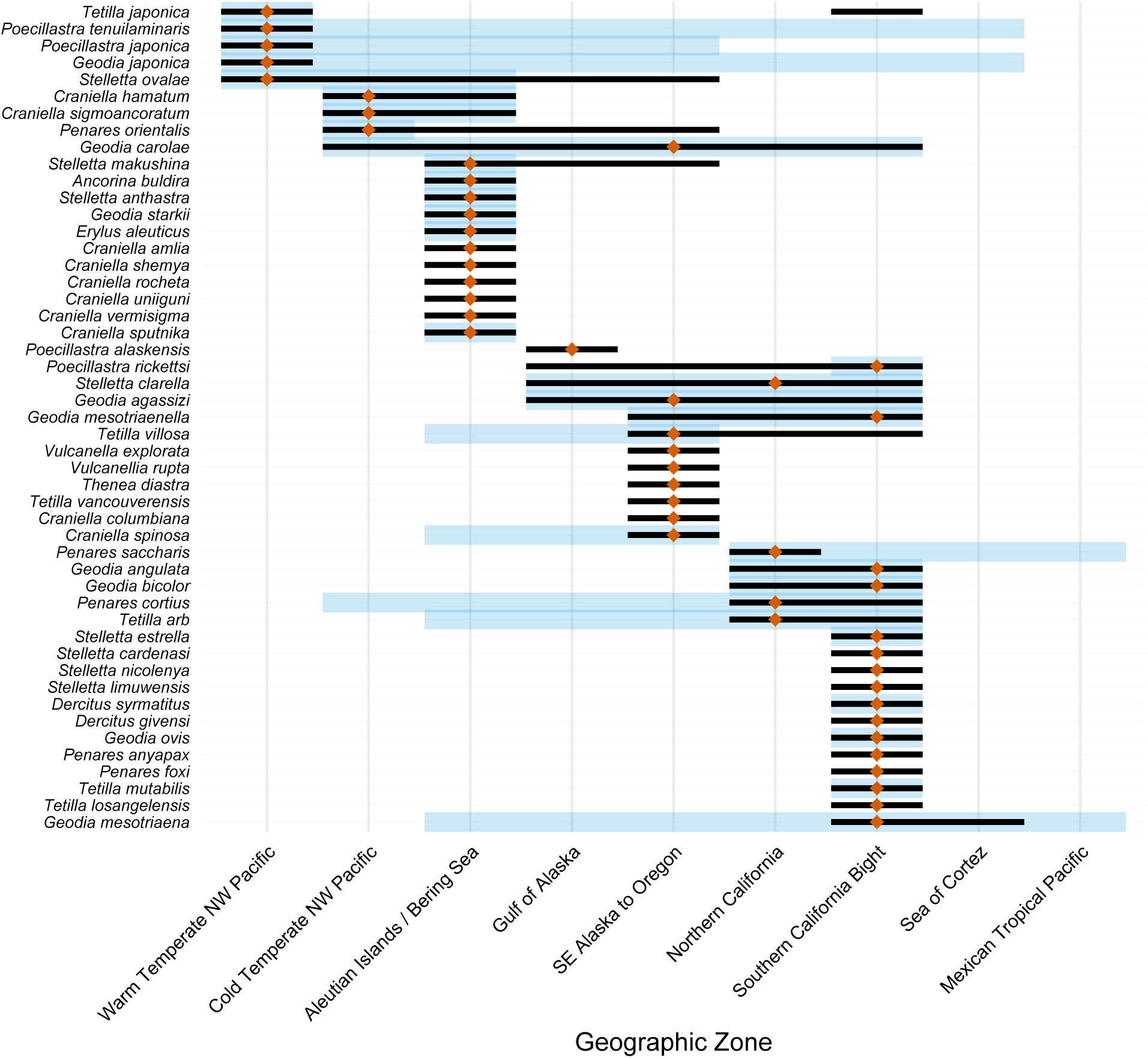
Hypothesized ranges of tetractinellid sponges. Species are included if they are currently or were previously proposed to overlap with the US/Canadian Northeast Pacific. The zone containing the type locality is marked with an orange diamond. Black lines show the newly proposed ranges for a species, and blue lines show the previously suspected ranges (which are absent for species newly described here or in Turner (2026)).

Despite the overall increase in species diversity in the region, these revisions have resulted in more range contractions (nine species) than range expansions (four species), with one species having both (*Tetilla villosa* (Lambe 1895)). This is in contrast to recent taxonomic revisions in this region in other sponge groups, where range expansions were the norm (Turner 2020; Turner *et al*. 2024; Turner & Pankey 2023). This difference could be due to many factors, including the proportion of deep-water species in the Tetractinellida, and a difference in geographic scope, as most other recent revisions were limited to the state of California, rather than the entire US-Canadian temperate zone.

These changes result in a reduction in the number of species with ranges that include both the warm temperate Northwest Pacific and the Northeast Pacific. These species were all described from the vicinity of Sagami Bay, Japan, where substantial taxonomic work on the Astrophorida has been conducted by a variety of authors (Sollas 1888; Tanita 1965; Tanita & Hoshino 1989; Thiele 1898). The results reported here make it seem less likely that sponges are distributed from this region into the Northeast Pacific. Of the four species previously postulated to have such ranges, only *Stelletta ovalae* (Tanita 1965) has survived this revision, and only with caveats. As detailed in the systematics section, I suspect that obtaining more data on *S. ovalae* from Japan — especially genetic data — may result in a split between this Japanese species and the morphologically similar samples found in Alaska and the Commander Islands. One possible exception to this pattern is *Tetilla japonica*, which may have been introduced from the warm temperate Northwest Pacific to the warm temperate Northeast Pacific via human means (Turner 2026). In contrast to sponges from Sagami Bay, a number of species are still thought to be shared between the cold Northwest and cold Northeast Pacific, which is expected due to the continuity of Russian and Alaskan waters. *Geodia carolae* stands out among these species because its range extends as far south as California. However, as discussed in the systematic section, only a single sample is known south of British Columbia, and it is morphologically distinct. This sample was not examined here, but the lack of other Pacific species of tetractinellids with such a broad latitudinal range should motivate future reexamination of the Californian sample as a possible new species.

Despite these range reductions, future work building on the results presented here will likely expand the known distributions of some taxa. This is especially true for newly described species, which will probably be identified from additional localities now that they are recognizable. Previously described species will also be easier to diagnose using the key provided here, allowing more voucher specimens to be confidently assigned to species names and likely broadening their known ranges. For example, the range of *Geodia mesotriaena* is restricted here to Southern California because all examined vouchers from farther north proved to belong to other species. Similarly, all samples of *Penares cortius* (de Laubenfels, 1930) collected north of California were reassigned to *P. orientalis* comb. nov. However, many additional vouchers assigned to these taxa were not examined in the present study, and future examination of this material will likely extend their known distributions to some degree.

Finally, I note that the abundance of large *Geodia* in Southern California has not been previously reported, as far as I am aware. I collected sponges at 12 locations near San Diego, in far Southern California. Six of these were below 25 m, and at these six, large (up to at least 25 cm) *Geodia* were among the most prominent and abundant features of the underwater landscape. Only two samples were collected, and both proved to be *G. mesotriaena*. This is an area with substantial human impacts, very near the city of San Diego, and frequented by divers and sport fishermen; one *Geodia* was seen that appeared to be healing from damage likely caused by an anchor strike (inaturalist.org/observations/339965473). It also remains unknown how widespread such dense aggregations may be: I searched for sponges at 93 other locations throughout Southern California between 2019 and 2026 and did not find this species at any of them, but only three of these locations included depths below 25 m. Collection time was limited at these depths, and no attempt was made to quantify the density or the biomass of these sponge gardens, but I hope that the observations reported here will motivate future work on this habitat.

## Systematic Section

Species known from the region:

Vulcanellidae

*Poecillastra rickettsi* de Laubenfels, 1930
*Poecillastra alaskensis* sp. nov.
*Vulcanella explorata* sp. nov.
*Vulcanella rupta* sp. nov.

Ancorinidae

*Ancorina buldira* Lehnert & Stone, 2014 (not examined here)
*Stelletta ovalae* Tanita, 1965
*Stelletta makushina* Lehnert & Stone, 2014
*Stelletta clarella* de Laubenfels 1930
*Stelletta anthastra* Lehnert & Stone, 2014 (not examined here)
*Stelletta estrella* de Laubenfels 1930
*Stelletta cardenasi* sp. nov.
*Stelletta nicolenya* sp. nov.
*Stelletta limuwensis* sp. nov.
*Dercitus (Stoeba) syrmatitus* de Laubenfels, 1930
*Dercitus (Stoeba) giveni* sp. nov.

Geodiidae Gray, 1867

*Geodia angulata* (Lendenfeld, 1910)
*Geodia bicolor* (Lendenfeld, 1910)
*Geodia carolae* (Lendenfeld, 1910)
*Geodia agassizii* Lendenfeld, 1910
*Geodia mesotriaena* Lendenfeld, 1910
*Geodia ovis* Lendenfeld, 1910
*Geodia mesotriaenella* Lendenfeld, 1910
*Geodia starki* Lehnert, Stone & Drumm, 2014 (not examined here)
*Erylus aleuticus* Lehnert, Stone & Heimler, 2006 (not examined here)
*Penares cortius* de Laubenfels, 1930
*Penares orientalis* comb. nov.
*Penares anyapax* sp. nov.
*Penares foxi* sp. nov.
*Penares saccharis* (de Laubenfels, 1930)

Theneidae

*Thenea diastra* sp. nov.

### Suborder Astrophorina Sollas, 1888

#### Definition

Demospongiae with asterose microscleres, sometimes with microxeas and microrhabds, and with tetractinal megascleres and oxeas radially arranged at least peripherally. From Hooper & van Soest (2002).

### Family Vulcanellidae Cárdenas, Xavier, Reveillaud, Schander & Rapp, 2011

#### Diagnosis

Astrophorina with calthrops, short-shafted triaenes or long-shafted triaenes, in addition to large oxeas and contort or sinuous strongyloxeas. Aster microscleres include several categories of streptasters (spirasters, metasters, amphiasters, and plesiasters). Monaxonic spicules consist of one to three categories of spiny microxeas or microstrongyles. From Kelly *et al*. (2019).

### Genus Poecillastra

#### Diagnosis

Vulcanellidae with spiny microxeas in a single category; triaenes are pseudocalthrops and/or short-shafted triaenes (Cárdenas & Rapp 2012).

### Species accounts

### Poecillastra rickettsi (de Laubenfels 1930)

Figures 6, 21B

**Figure 6.**
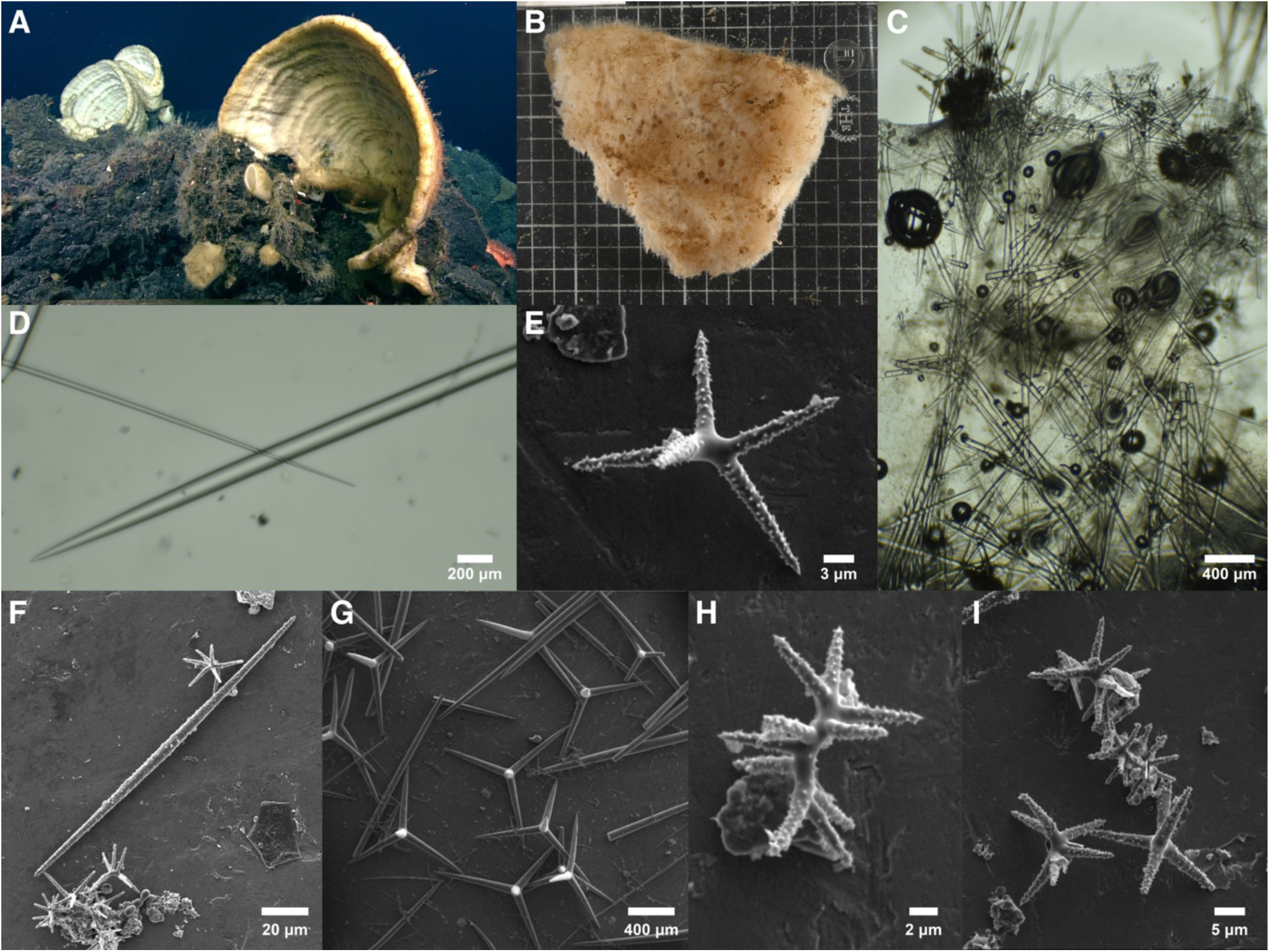
Poecillastra rickettsi. A: In situ photo taken by the ROV ROPOS / CCGS John P. Tully. B: Freshly collected sample before preservation (grid size is likely 1 cm). C: Perpendicular section through sponge surface, showing protruding bundles of oxeas II and confused internal skeleton of oxeas I and calthrops. D: Tips of thick oxea I and thin oxea II. E: Pleasiaster. F: Microxea and pleasiasters. G: Calthrops. H: Spiraster. I: Pleasiasters and spiraster. A-D from sample RBCM 025-00144-001, E-I from holotype. Panels A and B courtesy of Merlin Best, Fisheries and Oceans Canada.

#### Synonyms

*Poecillastra compressa* spp. *japonica* sensu Koltun 1966, in part

*Poecillastra tenuilaminaris* sensu de Laubenfels, 1932

*Poecillastra tenuilaminaris* sensu Green & Bakus, 1994

*Poecillastra* sp. A Green & Bakus, 1994

*Sphinctrella* sp. A Green & Bakus, 1994

#### Material examined

Holotype: USNM 21482, Monterey Bay, California, 80 m, July 1925. Other samples: RBCM 020-00056-002, Baby Bear Seamount, Cascadia Basin, British Columbia, (47.70630, -127.78550), to 2598 m, 8-Aug-1995; RBCM 018-00148-007, Anvil Island, Howe Sound, British Columbia, (49.53333, -123.29000), 21 m, 19-Aug-2011; RBCM 018-00210-001, Washington, depth and collection date unknown; CASIZ 179739, Cordell Bank NMS, California, (38.066283, -123.47035), 86 m, 7-Aug-2008; CASIZ 165832, Monterey Bay canyon wall, California, (36.733333, -122.04250), 500–600 m, 28-Feb-1997; CASIZ 180060, Cordell Bank NMS, California, (38.025, -123.42500), 62 m, 10-Oct-1979; RBCM 010-00247-002, Loudoun Canyon, British Columbia, (48.36832, -126.46247), 1416-1433 m, 29-Aug-2001; RBCM 988-00002-095, Rivers Inlet, Fitz Hugh Sound, British Columbia, (51.45000, -127.81667), 125 m, 19-Jan-1988; RBCM 991-00332-069, Skidegate Channel, British Columbia, (53.06333, - 132.97833), 1230 m, 21-Mar-1999; RBCM 999-00213-004, Rose Inlet, Moresby Island, British Columbia, (52.14517, -131.13150), 50 m, 18-Jun-1999; RBCM 025-00144-001, Dellwood Seamount, British Columbia, (53.50368, -136.05602), 892.76 m, 24-Aug-2024; USNM 21400, W. of Catalina Isl., California, depth unknown, 23-Jun-1916; USNM 1479000, Dixon Entrance near Dall Island, Alaska, (54.63389, -132.85361), 165 m, 9-Jun-2015; USNM 1479051, Offshore of Salisbury Sound, Alaska, (57.32139, -136.32111), 205 m, 9-Jul-1997; USNM 1478681, Shutter Ridge, West of Cape Ommaney, Alaska, (56.19660, -135.10200), depth unknown, 8-Jun-2015; CASIZ 235709, Offshore of Coos Bay, Oregon, (43.42667, -124.80667), 530-603 m, 27-May-1905; RBCM 014-00176-006, Fitz Hugh Sound, British Columbia, (51.72500, -127.98000), 253 m, 21-Oct-1982; SBMNH 467987, W. of Pt Conception, California, (34.459, -120.67270), 168-237 m, 16-Mar-1985.

#### Morphology

Samples are roughly cylindrical or irregularly massive, with one or a few oscula; a previous report stated that these oscula are sometimes surrounded by a palisade of very long spicules (de Laubenfels 1932). One roughly cylindrical sample was examined here, and it measured 5 × 3 cm in size, with a single apparent osculum; additional samples of this sort are described by de Laubenfels. The remaining samples are flat fragments of larger sponges. When the margin of the sponge is included in the sampled fragment, it appears to be rounded in profile, like the edge of a plate-, blade-, or vase-shaped sponge. One sample, which was photographed in situ before collection, formed a large horizontal vase (RBCM 025-00144-001, figure 6A). The sampled fragment of this sponge was roughly triangular, 12 × 9 cm in extent and 3 cm thick; other samples were 0.5–2.0 cm thick. These flat samples have many large pores (presumed oscula) scattered across the surface, somewhat irregular in size and distribution, each 1–3 cm in diameter and generally separated from the nearest neighboring osculum by 1–5 cm. In some cases, the oscula appear to occur on both sides, while in others, they are on one side only. The oscula of these flat fragments are not surrounded by a spicular palisade, and instead, very long protruding oxeas are individually scattered across the entire surface of the sponge, with dense protruding spicules along the edges of the blade. Samples are either white or dark brown post-preservation (figure 21B); at least some dark brown samples appear to be this color because they contain copious sediment. The holotype is tan post-preservation; only a small fragment was loaned to me for destructive sampling, and little about its morphology could be discerned.

#### Skeleton

The choanosomal skeleton is a confused reticulation of oxeas I. A layer of calthrops is present just below the surface as a disorganized jumble, rather than a flat layer supporting the ectosome, and calthrops are also seen in large numbers throughout the choanosome. Oxeas II pierce the surface individually or in flaring bouquets to create a hispid, furry surface. Asters form a dense surface crust and occur in lower numbers throughout the sponge. Microxeas do not appear to occur in the ectosome, and are instead scattered in the choanosome.

#### Spicules

Shown in figure 6 except where noted; see table 1 for a comparison of means among samples. A single, broken anatriaene was found in one sample, but it is presumed to be foreign; several samples contained glass sponge spicules and/or a few *Geodia* sterrasters.

**Table 1.** A subset of spicule dimensions from *Poecillastra* samples. Local. = locality (CA, California, BC, British Columbia, WA, Washington, AK, Alaska). Numbers are lengths (diameters for asters), in μm. Many samples have no unbroken oxeas II, in which case the maximum remaining length measured is shown in parentheses. The first row is the holotype for each species.

| Species | Voucher | Local. | cox1<br>clade | oxeas I<br>(mean) | oxeas II<br>(mean) | calthrops<br>(mean) | microxeas<br>(mean) | asters<br>(mean) | asters<br>(max) |
| --- | --- | --- | --- | --- | --- | --- | --- | --- | --- |
| *P. rickettsi* | USNM 21482 | CA | n/a | 2777 | 4220 | 683 | 208 | 20 | 28 |
| *P. rickettsi* | RBCM 020-00056-002 | BC | A | 3257 | (2481) | 571 | 164 | 24 | 42 |
| *P. rickettsi* | RBCM 018-00148-007 | BC | A | 1479 | (4749) | 382 | 161 | 16 | 27 |
| *P. rickettsi* | RBCM 018-00210-001 | WA | A | 1335 | 2580 | 336 | 155 | 15 | 20 |
| *P. rickettsi* | RBCM 010-00247-002 | BC | B | 3248 | (7306) | 791 | 207 | 20 | 27 |
| *P. rickettsi* | RBCM 988-00002-095 | BC | B | 2682 | (5990) | 503 | 226 | 16 | 23 |
| *P. rickettsi* | RBCM 999-00213-004 | BC | B | 1126 | 1655 | 233 | 157 | 15 | 52 |
| *P. rickettsi* | RBCM 025-00144-001 | BC | B | 2480 | 5490 | 608 | 142 | 19 | 27 |
| *P. rickettsi* | CASIZ 179739 | CA | B | 1497 | (2361) | 334 | 166 | 20 | 33 |
| *P. rickettsi* | USNM 21400 | CA | n/a | 1689 | (7449) | 388 | 141 | 19 | 32 |
| *P. rickettsi* | USNM 1479051 | AK | n/a | 2175 | 6003 | 564 | 183 | 20 | 37 |
| *P. alaskensis* | USNM 1512719 | AK | n/a | 2044 | 12998 | 422 | 162 | 37 | 96 |
| *P. alaskensis* | USNM 1699915 | AK | n/a | 2598 | 8610 | 677 | 163 | 29 | 86 |

Oxeas I: thick choanosomal oxeas (20–110 times as long as wide). Extremely variable in size among samples. All samples pooled: 379–1715–4925 × 7–34–95 μm (n=376). Holotype: 1466–2777–4925 × 43–63–95 μm (n=25). Sample with shortest oxeas I (RBCM 999-00213-004): 591–1126–1614 × 7–19–36 μm (n=113); Sample with longest oxeas I (RBCM 020-00056-002): 1677–3257–4525 × 22–37–50 μm (n=6).

Styles/strongyles: a minority of the thick choanosomal oxeas are modified into styles in some samples; these are usually quite rare, but were a significant minority in a few samples (e.g. RBCM 999-00213-004, RBCM 025-00144-001). Previously reported from the holotype, but not seen in spicules reisolated from that sample here. All samples pooled: 737–1445–2987 × 19–47–130 μm (n=34). Sample RBCM 999-00213-004 also contained a few strongyles, which were 617–733–905 × 32–36–43 μm (n=4). Not shown.

Oxeas II: long thin oxeas (140–400+ times as long as wide). Tapering abruptly to sharp points at the tips. Present in all samples, but in many, all examples are broken at one or both ends. In these cases, the remaining portion is sufficient to differentiate the class from oxeas I. All samples pooled: 1512–4828–7386 (n=22) × 8–19–37 μm (n=57). Holotype only: 3853–4220–4587 (n=2) × 18–22–28 μm (n=4). Previously reported as 17,000 × 15–30 μm from the holotype; I failed to reisolate these very long examples, and the more intermediate-length examples reported here were not mentioned in the type description (de Laubenfels 1932).

Calthrops: with straight or curved clads. Shapes are highly variable within samples, with one or more forked clads seen as a significant minority in some cases; when present, forks are usually near the tips of the clads. Spicules occasionally have more than four clads or some stylote clads. Clads are often unequal in length, but not dramatically so, as seen in the long-shafted triaenes in other species. As pointed out by Carvalho et al. (2011), the distinction between calthrops and orthotriaenes with rhabds nearly the same length as the clads is very subtle, and some spicules in this species could be categorized as the latter. Highly variable in size among samples; samples with longer oxeas generally have longer calthrop clads. The longest intact clad on each spicule was measured; all samples pooled: 100–480–1131 (n=417) × 10–42–98 μm (n=305). Holotype: 187–683–1067 × 18–61–98 μm (n=50). Sample with shortest clads (RBCM 999-00213-004): 111–233–414 × 10–18–35 μm (n=31); Sample with longest clads (RBCM 010-00247-002): 375–791–1131 × 23–58–78 μm (n=38).

Microxeas: completely covered in small spines. Most are evenly spined throughout, but a minority can be seen with spines clustered in a weakly striped pattern near the center of the spicule, and some are very weakly centrotylote. Spicules are gently curved in the holotype, but in many samples, the majority have a sharp bend in the center, and some samples are intermediate or mixed. Occasional styles or x-shaped spicules are seen. Lengths variable among samples, but only one size class in each sample. All samples pooled: 108–180–262 (n=232) × 3–6–10 μm (n=212). Holotype: 127–208–262 × 4–6–7 μm (n=32). Sample with shortest microxeas (RBCM 025-00144-001): 108–142–162 × 4–5–6 μm (n=13); Sample with longest microxeas (RBCM 988-00002-095): 176–226–256 × 6–7–10 μm (n=38).

Asters: the smallest asters were spirasters with many rays and a long centrum, but these graded into larger metasters with fewer rays and a short centrum, which graded into larger plesiasters with few rays and no centrum. No distinct size or shape classes could be identified, as the distribution of aster diameters from any one sample were continuous and unimodal, with shapes variable on a continuum. All were microspined. Diameters of all samples pooled: 7–18–63 μm (n=833). Holotype: 13–20–28 μm (n=52). All 19 examined samples were searched for large asters, to compare with *P. alaskensis* sp. nov. Only 2% of asters were over 30 μm in diameter. Asters 40–63 μm were rare, with single cases in three samples (USNM 1478681, RBCM 999-00213-004, RBCM 020-00056-002) and two cases in one sample (RBCM 014-00176-006).

Toxas: the type description reported toxas, "about 80 μm", which were likely foreign. Only one was seen in the spicules reisolated from the holotype (not measured), and none were seen in any other sample. Not shown in figure 6.

Spherules: occasional small spherical spicules were seen, including in the holotype, but not measured. Not shown in figure 6.

#### Distribution and habitat

The type locality is Monterey Bay, Central California, at 800 m depth. Samples assigned to this species and examined here range from the Alexander Archipelago, Southeast Alaska, to Catalina Island, Southern California. They were found across a wide depth range, from 21 m to at least 1,416 m. *Poecillastra* reported from the Gulf of California may also be this species, but it is not possible to tell from the limited information published (Dickinson 1945).

#### Remarks

To resolve the identity of *Poecillastra* species in the Northeast Pacific, I first aimed to determine how many species of *Poecillastra* were present among my samples. Two samples were found to have an additional class of very large plesiasters, and these are allocated to the new species *P. alaskensis* sp. nov., described below. Among the remaining samples, there was considerable variation in the size of the megascleres (table 1). Indeed, the sponges with the shortest and longest oxeas I were so different that the length distributions were non-overlapping, as detailed in the spicules section. Samples with longer oxeas I also had longer calthrop clads (Pearson correlation coefficient = 0.91, p = 0.0001). However, these variables were continuously distributed, rather than falling into categories, suggesting intraspecific variability. I did not attempt to correlate spicule size with sponge size, as all samples were collected as fragments, but there was a significant correlation of mean oxea I length with collection depth (Pearson correlation coefficient = 0.70, p = 0.04). A role for latitude was less apparent, as the longest and shortest spicules were both seen in samples from British Columbia.

Genetic variation was present at the cox1 locus among these samples, raising the possibility that two species may be present. Two samples from British Columbia and one from Washington fell in one mitochondrial clade (clade A in table 1), while all other samples (including others from British Columbia) fell into a second clade (clade B). These clades differed by only 4 base pairs (1.0% divergence), but this was similar to the divergence seen relative to species from other geographical regions (see results and figure 3). However, unlike divergence to the Atlantic *P. compressa*, these mitochondrial clades were not mirrored by differentiation at the 28S locus. There was also no correspondence between morphological variation and the mitochondrial genetic variation. As shown in table 1, the three representatives of mitochondrial clade A (the rarer clade) included the sample with the longest oxeas but also two samples that were among the shortest.

As I saw no discontinuous morphological variation among these samples, and no correlation of morphological variation with genetic variation, I concluded that there was no evidence for more than one species being present among them. The next task was to determine what name to assign to these samples. The hypothesis of a cosmopolitan *P. compressa* is rejected by both genetic and morphological data. Consistent with past reports (Carvalho *et al*. 2011), the large plesiasters common in North Atlantic *P. compressa* (Cárdenas & Rapp 2012) are lacking in this Pacific species, and the species are differentiated at both the cox1 and 28S loci (figure 3 and supplementary figures). The remaining species to consider are *P. rickettsi*, described from California, and *P. tenuilaminaris* and *P. japonica,* both described from Sagami Bay, Japan. Of these three species, *P. tenuilaminaris* is distinguished by lacking the second category of long, thin, ectosomal oxeas. These were found to be lacking both in the original species description and in samples examined from near the type locality by Lebwohl (Lebwohl 1914; Sollas 1886). In contrast to these Japanese samples, I found long thin oxeas in all samples investigated from the Northeast Pacific. Because they are long, thin, and pierce the surface of the sponge, unbroken examples were found in only some samples, but broken hair-like oxeas were easily found in all samples. This includes nine samples originally identified as *P. tenuilaminaris*, including the one identified as such by de Laubenfels, which contained a broken oxea whose remaining portion was 7449 × 36 μm. There is therefore no strong evidence supporting the presence of *P. tenuilaminaris* in the Northeastern Pacific.

If *P. tenuilaminaris* is rejected as a candidate, then these samples could be assigned to either *P. japonica* or *P. rickettsi*. The main characteristic differentiating *P. rickettsi* from all other *Poecillastra* is said to be the extreme length of the oxeas II, which were described as 17,000 × 15–30 μm by de Laubenfels. These were said to be "coronal oxeas" that surrounded the oscula in the type sample. Though I was loaned a small portion of the type sample for destructive sampling, this subsample did not include the oscula, so I was unable to verify that they are present. However, I did find intermediately sized oxeas II in the holotype that were not reported by de Laubenfels (e.g. 4587 × 18 μm, 3853 × 20 μm). A long oscular fringe of spicules has been described as only occasionally present in *P. compressa* (Cárdenas & Rapp 2012), so this trait may be variable in *P. rickettsi* as well. The intermediately-sized oxeas II that I found in the *P. rickettsi* holotype were a good match to the oxeas II found in the other samples examined, which, in the well-preserved sample RBCM 025-00144-001 ranged up to 7386 × 32 μm. Though long oxeas II are also described from *P. japonica*, they are described as being much thicker: >5000 × 65 μm, whereas the spicules found in my Northeast Pacific samples averaged 8–37 μm thick. The oxeas I in *P. japonica* are also described as thick, 110 μm vs. 63 μm in my reanalysis of the *P. rickettsi* holotype. The other Northeast Pacific samples measured had means 19–50 μm thick, and are therefore a better fit to *P. rickettsi* than *P. japonica*.

Assigning these samples to a species name is therefore not straightforward. The thickness of the oxeas I and oxeas II are both more consistent with *P. rickettsi* than *P. japonica*, while the lack of extremely long oxeas is more consistent with *P. japonica*. The presence of oxeas II in all samples argues against *P. tenuilaminaris*, but it is worth noting that the presence/absence of oxeas II was said to be the main feature distinguishing *P. compressa* and *P. schulzei* (Sollas 1886) in the North Atlantic, and a reanalysis found that they are actually present in both species (Cárdenas & Rapp 2012). Indeed, a *Poecillastra* sponge has been reported from Korea that has both oxeas I and oxeas II, with thicknesses more consistent with the samples reported here than the description of *P. japonica*. This sample was assigned to *P. tenuilaminaris* by the authors, but without discussion of other potential options (Rho & Sim 1981).

Despite these uncertainties, given the potentially variable nature of an oscular fringe in this species, and the difficulty in preserving such long spicules in deep water sponges, the morphological data is most consistent with these samples belonging to *P. rickettsi*. This assignment is strongly supported by biogeography, as *P. rickettsi* is described from Monterey Bay, Central California, and *P. japonica* and *P. tenuilaminaris* are described from Sagami Bay, Japan. Confirming this assignment will require finding a sample with very long coronal oxeas that has well-preserved DNA. I have been unable to amplify even short DNA sequences from any of de Laubenfels’ material, and the holotype of *P. rickettsi* was no exception. I was also unable to obtain any material of *P. japonica* or *P. tenuilaminaris* from the Northwest Pacific. Obtaining additional data from near the type localities for these species is also a high priority for future work. Finally, I note that there is a recently described species from the Bering Sea, *P. malyutini* Shilov *et al*., 2023, that is very similar to the updated concept of *P. rickettsi* reported here. Genetic data from the 28S locus was included in this species’ description, and is quite divergent from the sequences obtained here. However, this previous sequence is identical to those generated for *Vulcanella koltuni* Shilov *et al*., 2023 in the same work, raising the possibility that sample contamination occurred (see figure S2, supplementary materials).

### Poecillastra alaskensis sp. nov

Figure 7

**Figure 7.**
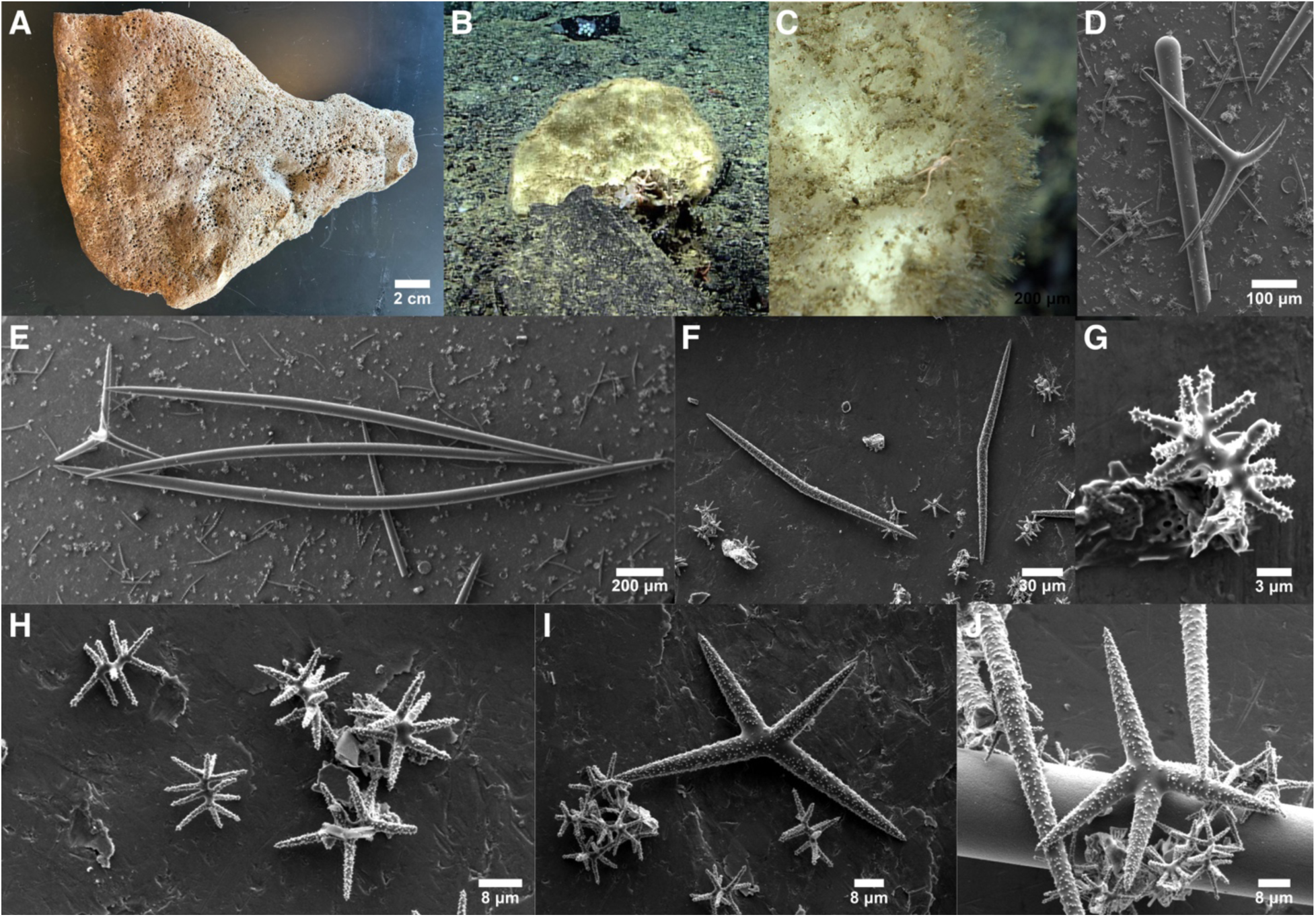
Poecillastra alaskensis. A: Dried holotype. B & C: In situ photos of USNM 1699915 taken by the ROV Deep Discoverer / NOAAS Okeanos Explorer. D: Head of style with calthrop. E: Oxeas I with calthrop. F: Microxeas. G: Spiraster. H: Metasters/plesiasters. I & J: Large plesiasters. All photos from holotype except B & C.

#### Synonyms

*Poecillastra compressa* spp. *japonica* sensu Koltun 1966, in part

#### Material examined

Holotype: USNM 1512719, N. of Pribilof Canyon, Bering Sea, Alaska, (56.41917, -171.39194), 235 m, 7-Dec-2016. Paratype: USNM 1699915, Denson Seamount, Gulf of Alaska, British Columbia, (54.13832, -137.37952), 1456 m, 31-Aug-2023.

#### Etymology

Named for the state of Alaska.

#### Morphology

Both samples are flat fragments of larger blade-, plate-, or vase-shaped sponges. The holotype is light brown and dried, and the paratype is white and preserved in ethanol. The holotype measures 21 × 17 × 2 cm, and the paratype measures 16 × 5 × 1 cm. Both have scattered pores on both sides, but they are more numerous on one side (figure 7A). In situ photographs of the paratype show a fan-shaped, off-white sponge that appears to be profusely hispid. The preserved portion of the paratype is more hispid on one side than on the other, with the greatest density of protruding oxeas along the sponge margin. The dried holotype is less obviously hispid, but very long protruding oxeas are scattered about the surface.

#### Skeleton

A thick layer of calthrops occurs near the sponge surface as a disorganized jumble, and no oxeas I are present in this region. Oxeas I form a confused reticulation below this zone, with some calthrops also present. Oxeas II pierce the surface individually or in small bouquets to create a hispid, furry surface. Small asters form a dense surface crust and are also present throughout the choanosome; large plesiasters are present only within the choanosome. Microxeas are not found in the ectosome or in the calthrop zone near the surface, and are instead scattered in the choanosome among the reticulation of oxeas I.

#### Spicules

Shown in figure 7 except where noted.

Oxeas I: thick choanosomal oxeas (40–150 times as long as wide). Slightly fusiform, but tapering more abruptly at tips. Holotype: 1204–2044–2707 × 23–38–51 μm (n=32); Both samples combined: 1104–2303–3447 × 16–40–64 μm (n=60).

Styles I: Less common than oxeas I, but easily found in both samples. Both samples combined: 978–2068–5917 × 29–55–63 μm (n=8).

Oxeas II: long thin oxeas (250–520 times as long as wide). Tapering abruptly toward the tips. All examples were broken in the holotype, where the remaining portion measured up to 13845 μm long; widths 38–75–104 μm (n=5). Paratype: 5705–8610–11157 × 13–24–38 μm (n=18). Not shown in figure 7; see figure 6D for similar example.

Calthrops: straight and curved clads are common, sharply bent or stylote clads are occasional. Clad lengths are often unequal. The longest intact clad on each spicule was measured; 283–422–640 × 23–36–67 μm (n=28) in the holotype, 283–560–1014 (n=61) × 23–40–67 (n=48) μm for Both samples combined.

Microxeas: curved or sharply bent near the center and completely covered with small, evenly distributed spines. Microxeas are weakly centrotylote in the paratype but not the holotype. Holotype: 128–162–184 × 5–7–9 μm (n=20); Both samples combined: 123–163–190 (n=43) × 3–6–9 μm (n=25).

Large plesiasters: the distribution of total diameter of all asters was bimodal; asters over 40 μm in diameter were considered the large size class, but these graded into the smaller plesiasters. Large plesiasters had 3 to 5 rays and were entirely microspined; they were less common than small plesiasters, but easily found in moderate numbers. Holotype diameters: 42–66–96 μm (n=32); Both samples combined: 40–63–96 (n=46).

Small spirasters-plesiasters: the smallest asters were spirasters with many rays and a long centrum, but these graded into larger metasters with fewer rays and a short centrum, which graded into larger plesiasters with few rays and no centrum. Distinct size or shape classes could not be found, as the distribution of aster diameters from any one sample were continuous and unimodal, with shapes variable on a continuum. Diameters of both samples pooled: 11–24–39 μm (n=140). Holotype: 11–22–34 μm (n=61).

#### Distribution and habitat

The holotype was collected from 235 m depth near the Pribilof Islands, Alaska, in the Bering Sea. The other sample was collected at 1456 m on the Denson Seamount in the Gulf of Alaska. Koltun (1966) described *Poecillastra* from the Northwest Pacific with very large asters, which may be this species as well (exact locations unclear).

#### Remarks

This species is differentiated from all previously named *Poecillastra* by the presence of very large plesiasters. I found that, for 19 of the 21 *Poecillastra* I examined from the region, the asters were mainly between 10–30 μm, with rare examples expanding the range to 7–63 μm. In contrast, two samples had a second, common class of plesiasters 40–96 μm. These are larger than any asters reported from any previous *Poecillastra*, with the only similar asters from *P. antarctica* Koltun, 1964 and *P. maremontana* Carvalho *et al*., 2011. These two species are unlikely congeners based on biogeography, and also have fewer (in the case of *P. antarctica*) or more (for *P. maremontana*) classes than seen here, so these two samples are here named *P. alaskensis sp. nov*. The only genetic data successfully generated for this species was a very short (213 bp) portion of the 28S D1D2 barcoding locus. There were no differences between this sequence and those from *P. rickettsi*, so it does not support the new species name. However, given its short length, this is hardly conclusive.

In his description of *P. compressa japonica* from the Northwest Pacific, Koltun (1966) noted that the asters ranged in size from 14–70 μm, but that the largest class of asters were sometimes missing. It seems possible that the examples with the largest asters were members of the new species described here, but unfortunately, Koltun did not state where these sponges were found.

### Genus *Vulcanella* Sollas, 1886

#### Diagnosis

Vulcanellidae with spiny microxeas in one to three categories, with a more-or-less conspicuous ringed ornamentation; triaenes are calthrops and/or short-shafted triaenes or long-shafted triaenes. From Cárdenas *et al*. (2011).

#### Species accounts

Vulcanella explorata sp. nov.

Figure 8

**Figure 8.**
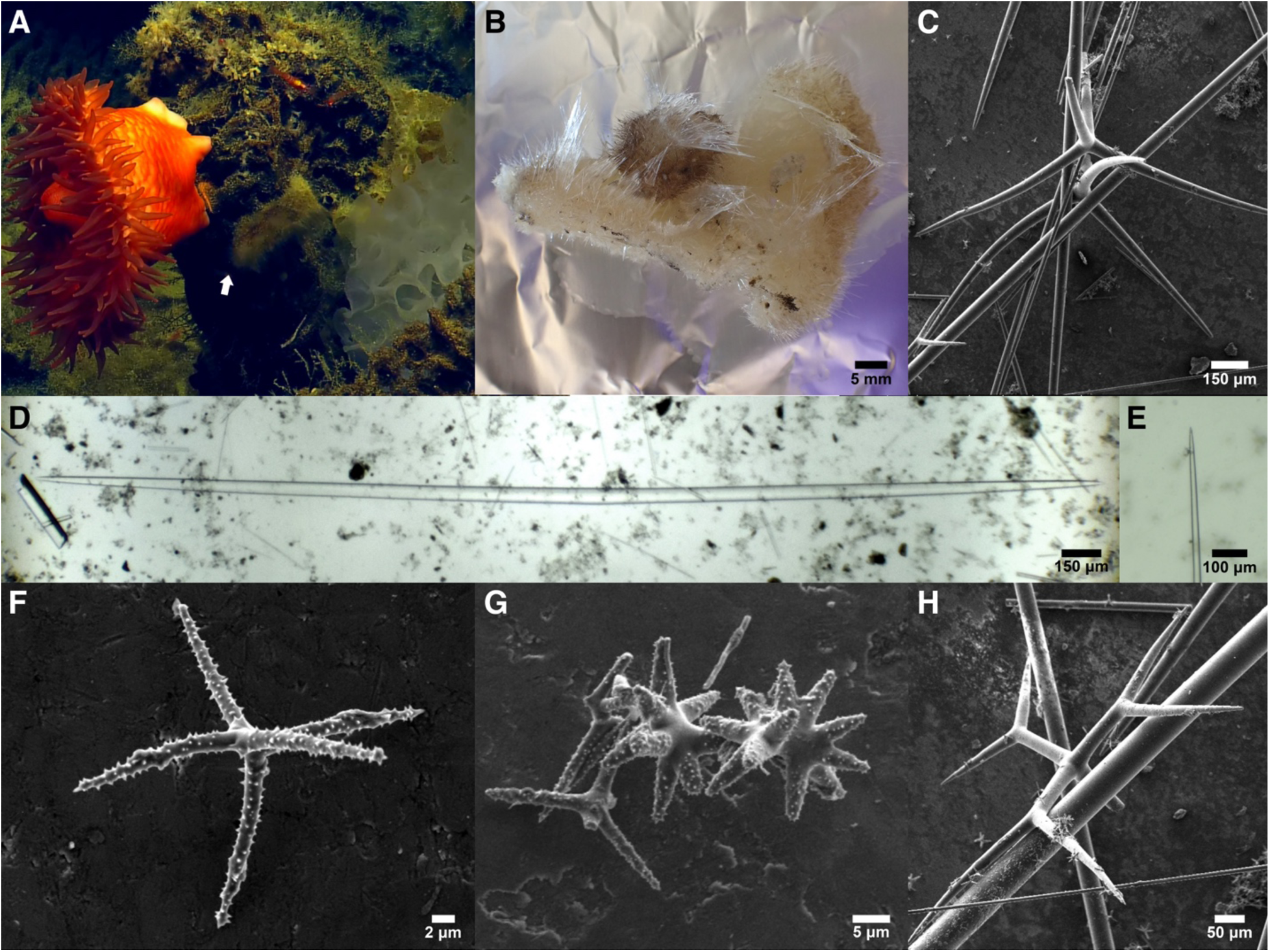
Vulcanella explorata. A: In situ picture taken by the ROV Hercules / EV Nautilus; the entire black structure was sampled, and the holotype and paratype were both removed from it; the arrow indicates a sponge that is likely the paratype. B: Preserved holotype. C: Calthrops. D: Oxea I. E: Tip of oxea II. F: Plesiaster. G: Spirasters. H: Dichotriaene with broken rhabd. All spicules from holotype.

#### Material examined

Holotype (RBCM 018-00975-006) and paratype (RBCM 018-00975-005), both collected at the Explorer Seamount, (49.05900, -130.94180), 795 m, 19-Jul-2018.

#### Etymology

Named for the Explorer Seamount and the United States Coast and Geodetic Survey ship *Explorer*, for which the seamount is named.

#### Morphology

The holotype is a roughly triangular sponge, 56 × 56 × 42 mm across, and 10 mm thick. The paratype is similar, but rectangular, 58 × 19 mm across and 18 mm thick. Samples were white *in situ* and remain white in ethanol. Oscula are clustered in one or two roughly circular depressions, which are surrounded by a fringe of very long (∼12 mm) protruding oxeas II. The rest of the sponge is covered in a shorter, but still very long (∼5 mm) fur of protruding oxeas.

#### Skeleton

Choanosomal skeleton is a confused reticulation of oxeas I. Calthrops are found only near the surface or protruding slightly. Oxeas II form flaring bundles that pierce the surface to create a hispid, furry surface. Both aster types are found in the ectosome and densely scattered throughout the choanosome.

#### Spicules

Shown in figure 8 except where noted.

Oxeas I: thick choanosomal oxeas (50–150 times as long as wide). Slightly fusiform, but tapering more abruptly at tips; occasional styles also seen. Holotype: 2559–4301–5605 × 25–59–76 μm (n=15); Both samples combined: 2559–4899–7560 × 25–58–79 μm (n=30). A few are modified to styles.

Oxeas II: long thin oxeas (250–400 times as long as wide). Tapering abruptly towards the tips. Holotype: 11665–12386–13895 × 32–37–43 μm (n=4); Both samples combined: 9907–11619–13895 × 27–34–43 μm (n=6). Most are broken, and longer examples likely occur.

Calthrops: most with long, thin, curving clads, but some are straight. Highly variable, and some could be categorized as plagiotriaenes. Occasional forked clads or strongylote clads are seen. The longest intact clad on each spicule was measured; holotype: 518–900–1249 × 31–53–76 μm (n=32); Both samples combined: 518–930–1439 × 31–51–76 μm (n=54).

Dichotriaenes: With rhabds and clads of similar length. Less common than calthrops, but not rare in either sample. The deuteroclad is two to three times as long as the protoclad. Occasional strongylote clads or rhabds are seen. Rhabds 503–629–752 × 43–50–67 μm (n=4), all from the paratype; clads 414–517–658 × 25–32–38 μm (n=4) in the holotype, 414–690–1063 × 25–44–63 μm (n=10) in Both samples combined.

Plesiasters: the most abundant type of aster, with long, thin, acanthose rays and little to no unspined central shaft. Diameter 20–29–41 μm (n=33) in holotype, 16–26–41 μm (n=61) in Both samples combined; ray widths at base 1.2–2.0–3.8 μm (n=50).

Spirasters/metasters: also common in both samples, these asters have thicker, but slightly shorter, acanthose rays than plesiasters, and a large unspined central area. Diameter 19–25–30 μm (n=20) in holotype, 18–24–30 μm (n=27) in both samples combined; ray widths at base 1.8–3.0–4.3 μm (n=23).

#### Distribution and habitat

Both samples were collected at 795 m depth on the Explorer Seamount, approximately 230 km west of Vancouver Island, British Columbia. They appeared to be growing on the skeleton of a dead glass sponge.

#### Remarks

This species is confidently placed within the Vulcanellidae clade in both the cox1 and 28S phylogenies, but there are no closely related species with DNA data, and its position within the clade is poorly resolved. At cox1, it is sister to all other Vulcanellidae with data available (figure S1), whereas it is nested well within the family at the nuclear locus (figure S2).

Morphologically, it is somewhat ambiguous whether this species should be placed within *Vulcanella* or *Poecillastra*, because current definitions emphasize the number of categories of microxea as the most important character distinguishing them, and microxeas are absent in this species (Cárdenas *et al*. 2011). I have placed it within *Vulcanella* because this genus tends to have tetractinal spicules found only near the ectosome, rather than scattered throughout the choanosome, as seen here; *Vulcanella* also tend to have oscula grouped into oscular areas, as seen here; and most importantly, one species is already known to have no microxeas. The spicule complement of this species, *V. theneides* (Burton 1959), is similar to the new species described here, and these species could be related. The species are readily distinguished by the previously named species having smaller calthrops, lacking oxeas II, lacking spirasters, and having only rare dichotriaenes. These sponges are also highly unlikely to be the same species for biogeographic reasons, as the previous species is known only from the Maldives.

### Vulcanella rupta sp. nov

Figure 9

**Figure 9.**
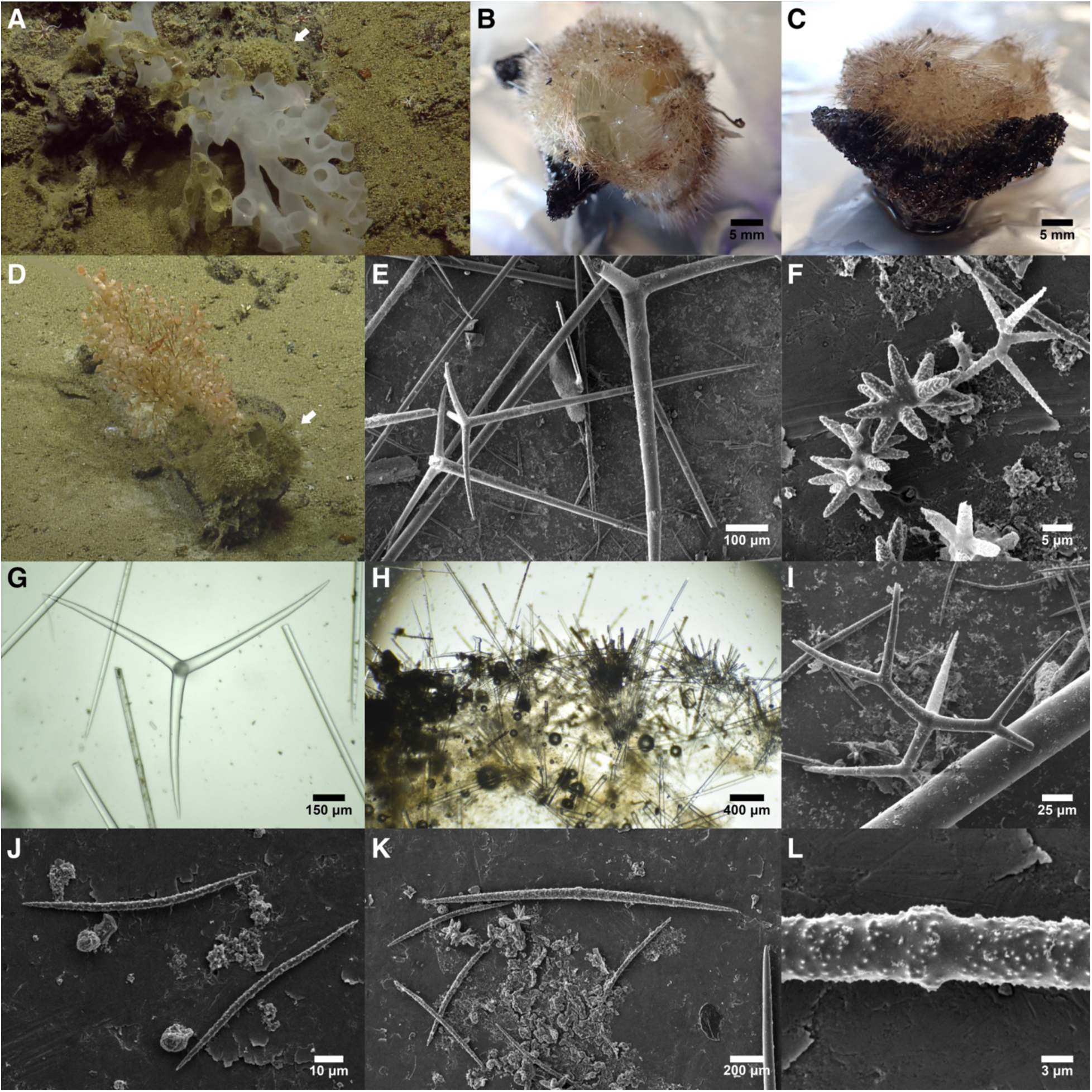
Vulcanella rupta. A: In situ picture taken by the ROV Hercules / EV Nautilus; arrow indicates holotype. B & C: Preserved holotype. D: In situ picture taken by the ROV Hercules / EV Nautilus; arrow indicates sample RBCM 018-00879-002. E: Long-shafted orthotriaenes. F: Plesiaster (upper right) with two spirasters. G: Calthrop from RBCM 018-00967-001. H: Perpendicular section of the sponge surface showing bouquets of protruding oxeas II, triaenes just below the sponge surface, and a confused choanosomal skeleton; sample RBCM 018-00967-001. I: Dichotriaene. J: Small microxeas. K: One large microxea with several small microxeas. L: Centrotylote region of large microxea shown in K. All images from holotype except where indicated.

#### Material examined

Holotype (RBCM 018-00898-006) and paratype (RBCM 018-00898-002), both collected at Hodgkins Seamount, (53.51030, -136.00310), 600-1425 m, 11-Jul-2018. Other samples: RBCM 018-00879-002, Dellwood Seamount, (50.72140, -130.91910), ∼835 m, 7-Jul-2018; RBCM 010-00538-001, off Brooks Peninsula, (50.00575 – 50.00603, -127.966267 – -127.99633), 1245-1296 m, 1-Sep-2004; RBCM 023-00023-003, SG̲áan K̲ínghlas Seamount, GPS location and depth not indicated, 20-Jun-2022; RBCM 018-00967-001, Dellwood Seamount, (50.57920, -130.70550), 1040 m, 18-Jul-2018.

#### Etymology

Named for its resemblance to a sphere ruptured by an internal explosion.

#### Morphology

Off-white, subspherical sponges covered in long protruding spicules. The largest sample is roughly subspherical but compressed vertically, at 7 × 6 × 3 cm, while the smallest is closer to spherical at 1.5 × 1.2 × 1 cm. Each sponge has two to four large, prominent craters that are surrounded by a very long (1–3 cm) palisade of protruding spicules. The internal area of these craters is the only part of the sponge not covered in protruding spicules; instead, they are smooth, membranous, and contain holes of greatly varying size. Some areas within the craters are covered by a membranous sieve, but others are not.

#### Skeleton

Perpendicular sections near the surface revealed the skeleton to be a confused, halichondroid network of oxeas I and occasional styles. Some choanosomal spicules could be described as bundled, but this was mainly due to the many open spaces concentrating and aligning the oxeas. At the surface, there were many protruding oxeas I and II. The oxeas II were primarily in bouquets. Microxeas were concentrated at the surface in a tangential layer, with a few also scattered internally. Asters were seen throughout. Triaenes were also concentrated at the surface, with some slightly protruding. Long-shafted triaenes and dichotriaenes had the rhabd pointing inward, with the clads providing a near-surface layer of additional spiculation.

#### Spicules

Shown in figure 9 except where noted.

Oxeas I: thick choanosomal oxeas (30–155 times as long as wide). Slightly fusiform, tapering gradually to sharp tips. Holotype: 2800–4224–4965 × 48–62–80 μm (n=9); all samples pooled: 1172–3049–5636 × 10–41–90 μm (n=70). A few are modified to styles. Not shown in figure 9; see figure 8D for similar example.

Oxeas II: long thin oxeas (160–330 times as long as wide). Gradually tapering to sharp tips, but occasionally modified to styles. Holotype: 5949–9466–12931 × 25–40–50 μm (n=11), all samples pooled: 3666–8199–12931 × 13–36–57 μm (n=19). Most are broken, and longer examples should occur, as some spicules protrude nearly 3 cm. Not shown in figure 9; see figure 8E for similar example.

Long-shafted orthotriaenes: with long, straight, gradually tapering rhabds and three equal-length clads that are nearly orthogonal to rhabd, but slightly curved. Holotype: Rhabds 814–1143–1478 × 22–46–73 μm (n=12), clads 160–490–813 × 24–41–62 μm (n=12). All samples pooled: rhabds 814–1340–2306 × 22–56–86 μm (n=42), clads 160–539–813 × 24–49–67 μm (n=41).

Calthrops: with straight or curved clads. Occasional forked clads are seen. The longest intact clad on each spicule was measured. Holotype: 483–1042–1300 × 36–62–77 μm (n=11); all samples pooled: 133–934–1571 × 9–65–105 μm (n=63).

Dichotriaenes: With rhabds and clads of similar length. Uncommon but not rare. Rhabds 78–691–1049 × 12–60–87 μm (n=8); clads 132–571–1115 × 9–50–89 μm (n=12). The portion of the clad distal to the fork is two to four times as long as the proximal portion.

Microxeas: Often sharply bent in the middle, but many were evenly curved or straight. The small size class is evenly acanthose throughout, with no ringed pattern. The large size class are often centrotylote, with the tylote bulge acanthose and spines absent to either side, creating a weak central ringed pattern; the remainder of the spicule is evenly acanthose with no ringing. A minority of the larger class are entirely smooth. Length distributions of four of five samples were strongly bimodal, so two size classes are generally seen in each sample. However, the breaks between size classes were inconsistent, varying from 150 μm to 225 μm. All spicules combined were 63–171–416 × 2–6–11 μm (n=255). The holotype had the smallest size classes, with a larger size class 161–201–324 × 6–7–11 μm (n=28) and a smaller size class 63–81–144 × 2–3–6 μm (n=52). All other samples had noticeably larger microxeas, especially for the smaller size class. For example, RBCM 010-00538-001 had a larger size class 202–244–294 × 6–8–11 μm (n=22) and a smaller size class 122–166–198 × 4–6–10 μm (n=45).

Spirasters: With long central axis and shorter rays; rays are variable in thickness but often thick and blunt; acanthose throughout. Diameter, holotype: 12–21–27 μm (n=35); all samples pooled: 12–20–27 μm (n=63); ray lengths 3–7–9 μm (n=63). The holotype was approximately 90% spirasters, with most having thick, blunt spines, and only 10% plesiasters. Other samples varied from roughly even abundances of both types to only 5% spirasters and 95% plesiasters.

Plesiasters: With long, thin, pointed rays and short central shaft; acanthose throughout. Diameter, holotype: 22–24–29 μm (n=4); all samples pooled: 15–24–35 μm (n=56); ray lengths 5–11–18 μm (n=52).

#### Distribution and habitat

Known from seamounts off British Columbia between 600 m and 1425 m depth. Of the four samples with in situ photos, three appeared to be growing on the skeletons of dead glass sponges and one may have been attached to rocky substrate.

#### Remarks

I was able to assign 6 samples to this species concept based on genotypes at the cox1 and 28S loci, and examined the spicules from five of these (one sample was a very small fragment). There is a surprising amount of variation in the size of the microoxeas and in the proportion of spirasters relative to plesiasters. There is also some genetic variation at the 28S locus, but this was not mirrored at the cox1 locus, where all samples were identical. It is possible that a larger sample will discover associations between genetic variation across compartments, or between genetic and morphological variation, but with the evidence available, these individuals are best accommodated within a single species concept.

This species is placed in *Vulcanella* based on the two size classes of microxeas. Morphological data on all previous species was helpfully compiled by Kamenev *et al*. (2023), and no new species have been described since then (de Voogd, *et al*. 2025). The combination of long-shafted triaenes, calthrops, and dichotriaenes differentiates this species from all others. The most similar species is *V. porosa* (Lebwohl 1914). However, the description and photomicrographs of this species clearly indicate that the species has ringed microxeas, with spines on the rings, which was never seen in the new species. Additional differences include a lack of dichotriaenes, much smaller clads on triaenes, and no plesiasters (Koltun 1966; Lebwohl 1914).

### Family Ancorinidae Schmidt, 1870

#### Diagnosis

Encrusting, irregularly massive or spherical sponges, in some cases with long inhalant and/or exhalant tubes. The megascleres are oxeas and triaenes with a long rhabdome. Microscleres are euasters (oxyasters, spherasters, chiasters, tylasters), streptasters (sanidasters, amphiasters), and spiny or smooth microrhabds. From Uriz (2002).

### Genus Stelletta

#### Diagnosis

Massive sponges with a more-or-less collagenous-rich cortex; triaenes often abundant, more rarely absent, oxeas, and from one to three types of euasters, one of them confined to the choanosome, the other(s) sparse throughout the sponge. Occasional accessory ortho- or trichodragmata. From Uriz (2002).

### Species accounts

### Stelletta ovalae Tanita, 1965

Figure 10

**Figure 10.**
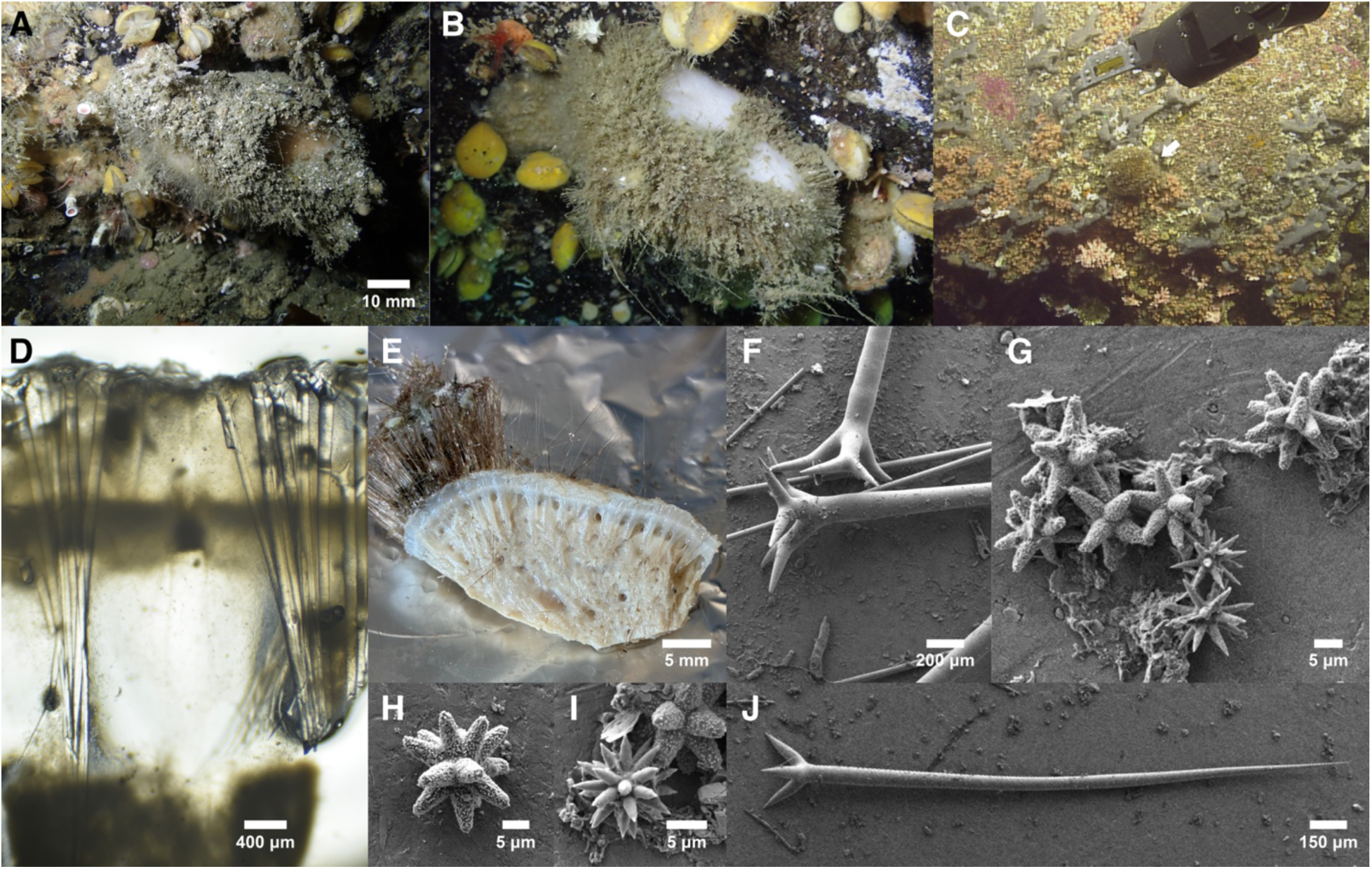
Stelletta ovalae. A: In situ picture of tube-shaped sample TLT1674. B: In situ picture of massive sample TLT1562. C: In situ picture of oval sample RBCM 018-00914-001 taken by the ROV Hercules / EV Nautilus; arrow indicates sponge. D: Perpendicular section of the sponge surface showing bouquets of triaenes supporting ectosome. E: Sample post-preservation, showing cortex, tracts of megascleres supporting cortex and delineating subcortical space, and long protruding oxeas II. F: Clads of dichotriaenes. G: Acanthose strongylasters and smooth oxyasters. H: Acanthose strongylaster. I: Smooth oxyaster. J: Plagiotriaene. All images from TLT1562 except where indicated.

#### Material examined

TLT1674, Bird Island, Juneau, Alaska, (58.49035, -134.85168), 31 m, 27-Jun-2025; TLT1562, Aaron Island, Juneau, Alaska, (58.44080, -134.82440), 32 m, 26-Jun-2025; CASIZ 302615, Deep Inlet, Sitka, Alaska, (56.98150, -135.29117), 33 m, 5-Sep-2010; CASIZ 302624, Bird Island, Juneau, Alaska, (58.49117, -134.84983), 28 m, 11-Sep-2010; RBCM 018-00914-001, SG̲áan K̲ínghlas-Bowie Seamount, British Columbia, (53.30100, -135.65020), to 62 m, 12-Jul-2018.

#### Morphology

Exterior is white in life and post preservation; interior is yellowish-white to beige. Sponges are densely spiculated, firm and unyielding. A cartilaginous cortical layer approximately 2 mm thick is prominent at the ectosome, with large subectosomal spaces below it. Oxeas project through the ectosome to form a thick spicule fur over all or nearly all of the sponge surface, over a centimeter thick in some samples. This extremely hispid surface traps sediment and debris in some habitats, leading to a brown appearance in situ. Samples examined here were subspherical, amorphously massive, or tube-shaped, while published records are globular or ellipsoidal. The largest known sample is 14 cm across and 5 cm thick, but it is a fragment and was larger in life. Oscula not apparent.

#### Skeleton

Sections tangential to the sponge surface reveal columns of megascleres rising to the surface, with oxeas continuing past the ectosome to protrude by a centimeter or more. Triaenes are found at the ectosome, with nearly all cladomes directly below the sponge surface. Asters form a dense layer at the surface, but are also scattered throughout.

#### Spicules

Shown in figure 10 except where noted.

Oxeas I: Thick oxeas, tapering to sharp points at both ends, though occasional styles are also seen; 5474–7651–9801 × 70–85–97 μm (n=17). Japanese material previously described as "up to 10000" × 70–90 μm; thin oxeas not mentioned (Tanita & Hoshino 1989). Aleutian sample previously described with only thin oxeas (Lehnert & Stone 2014). Not shown; see figure 8D for similar example.

Oxeas II: Thin oxeas, tapering to sharp points at both ends; 8817–13701–19117 × 25–50–62 μm (n=6). Japanese material was previously described as having only thick oxeas (Tanita & Hoshino 1989). Aleutian sample previously described as 8000–10,500 × 18–38 µm (Lehnert & Stone 2014). Not shown; see figure 8E for similar example.

Dichotriaenes: The primary triaene. With long rhabds and clads that tend to angle slightly downwards from the rhabd, then bend and flatten out orthogonal to the rhabd. Occasional strongylote clads or rhabds are seen. Rhabds 4987–6597–7633 (n=30) × 110–153–185 (n=42) μm; clads 324–540–729 (n=41) × 89–113–130 (n=20) μm. Japanese material previously described with rhabds "up to 10000" × 120 μm, clads 300 × 80 μm (Tanita & Hoshino 1989). Aleutian sample previously described with rhabds 4500–5600 × 180–230 µm, clads 380–525 × 180–225 (Lehnert & Stone 2014).

Plagiotriaenes: Similar to dichotriaenes but smaller and much less common. The clads are usually straight, in contrast to the dichotriaenes which have bent clads and a flat, forked "foot". A few seen with stylote clads. Rhabds 2232–2471–2711 × 72–76–79 μm (n=2); clads 205–213–222 μm (n=2). Not mentioned in previous descriptions.

Anatriaenes: Not seen in the samples examined, but described in previous publications; they were said to protrude from the base of the sponge (Lehnert & Stone 2014), and the samples examined here were fragments that may not have had that portion included. Japanese material previously described with rhabds "up to 10000" or more × 30 μm, clads 100 μm (Tanita & Hoshino 1989). Aleutian sample previously described with rhabds 2300– 6400 × 40–70 µm, clads 35–78 × 34–56 µm (Lehnert & Stone 2014).

Acanthose strongylasters: The most common type of aster. With many (>8) rays, slightly conical, but with blunt, rounded ends. Completely covered in small spines. The sample examined with SEM had diameters 12–17–26 μm (n=65). Japanese material previously described as oxyasters with 8 or more rays, diameters 20–25 µm (Tanita & Hoshino 1989). Aleutian sample previously described with "oxyasters with conical, blunt rays" 18–24 µm (Lehnert & Stone 2014).

Non-acanthose oxyasters: With many (>20) conical rays with sharp tips; unspined or lightly spined. The sample examined with SEM had diameters 11–14–16 μm (n=18). Japanese material previously described as oxyasters with numerous rays up to 10 µm (Tanita & Hoshino 1989). Aleutian sample previously described with oxyasters 9–12 µm (Lehnert & Stone 2014).

Two additional samples were examined with light microscopy, but no attempt was made to separate the 2 types of asters. These samples had asters 10–21–31 μm (n=33).

#### Distribution and habitat

Known from Honshu, Japan, the Aleutian Island of Unalaska, and now Southeastern Alaska and British Columbia. It is postulated in the remarks below that *S. validissima* form *validissima* sensu Koltun (1966) may be the same species, in which case it is also known from the Southern Kuril Islands, Russia, and the Commander Islands in the Bering Sea. Known from 31–177 m, but samples shallower than 60 m were all collected in fjords in Alaska where deep-water species are known to occur at shallower depths.

#### Remarks

No previous DNA data is available for samples identified as *S. ovalae*. This species was originally described from Japan, but morphological evidence points to a possible range extending from Japan, through the Russian Pacific and Aleutian Islands, then south to British Columbia.

The samples examined here, from Southeast Alaska and British Columbia, are likely to be the same species as a sample previously described from Unalaska Island, in the Aleutians (Lehnert & Stone 2014). These samples are also a good morphological match to samples from the Commander Islands and Southern Kuril Islands in the Russian Pacific (Koltun 1966). These Russian samples were described as *S. validissima* form *validissima*, and have larger maximum aster sizes than the samples I examined, but otherwise seem like a good match. Notably, the Russian samples match *S. ovalae* better than *S. validissima* Thiele, 1898 based on the sizes of the asters, which are slightly larger than those described for *S. ovalae* but much larger than those described for *S. validissima*.

I therefore agree with Lehnert & Stone that their Aleutian sample, and the samples examined here, cannot be differentiated morphologically from the Japanese *S. ovalae*. However, given the morphological similarities between other good species of *Stelletta* described below, obtaining more data — especially DNA data — from Japanese *S. ovalae* is a high priority for future work.

A previously published checklist of sponges from British Columbia included *S. validissima*, describing its range as Japan to the Bering Strait to British Columbia (Austin 1985). I know of no published information on the samples leading to this claim, but speculate that they were based on a comparison with Koltun’s *S. validissima* form *validissima*, and therefore referring to this species.

Genetic data obtained here shows that this species is not closely related to any others in the region, and instead forms a clade with Atlantic and Mediterranean *Stelletta* (figures 4, S1, and S2).

### *Stelletta makushina* Lehnert & Stone, 2014

Figure 11

**Figure 11.**
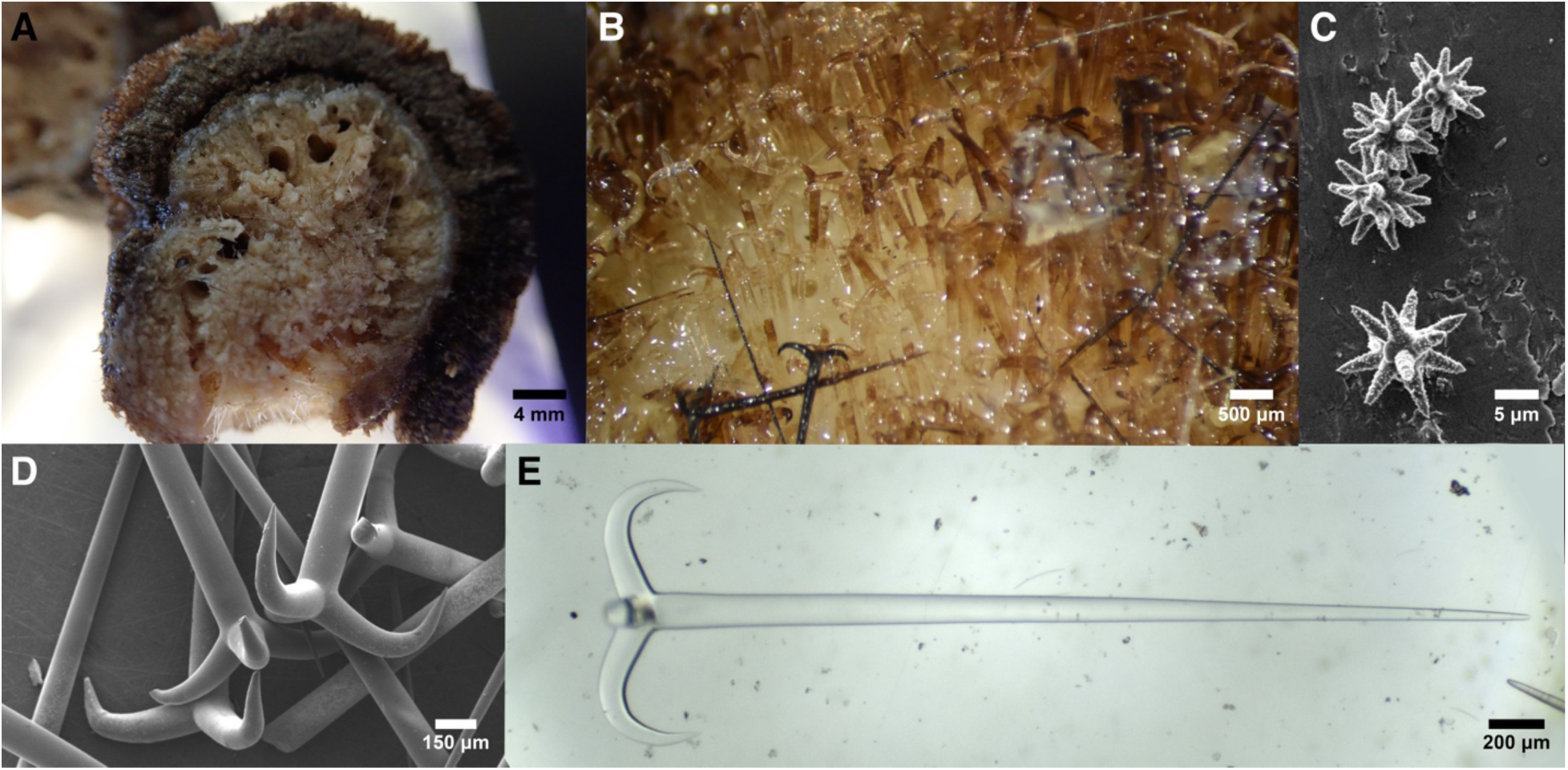
Stelletta makushina. A: Preserved and sectioned sample confined on most sides within a tubular hexactinellid skeleton (darker brown layer). B: Image of sponge surface showing protruding triaene cladomes. C: Oxyasters. D: SEM of orthotriaene cladomes. E: Orthotriaene. All images from sample RBCM 019-00567-006.

#### Material examined

RBCM 019-00567-006, Malcolm Island, British Columbia, (50.6442, - 126.9981), depth not recorded, 28-Sept-2007.

#### Morphology

Exterior is white alive and post preservation, but sponge may appear brownish due to algal growth or detritus on protruding spicules; interior is yellowish-white. A cartilaginous cortical layer approximately 1 mm thick is present at the surface, with the cladomes of projecting triaenes forming a dense layer of spicules above it. The sample examined here was 2 cm in diameter, growing within a tubular skeleton of a dead hexatinellid sponge. The other known sample is ovoid, 45 mm × 38 mm × 32 mm (Lehnert & Stone 2014). Oscules not apparent.

#### Skeleton

Columns of megascleres radiate from the interior, fanning out just below the surface. Triaenes are oriented with the cladomes facing out, and these pierce the sponge surface to create a relatively even layer of cladomes outside of the sponge surface. Interior skeleton composed primarily of oxeas.

#### Spicules

Shown in figure 11 except where noted.

Oxeas: Thick oxeas, tapering to sharp points at both ends; 819–3990–6412 × 21–67–99 μm (n=30). Holotype previously described 4460–7600 × 30–160 µm (Lehnert & Stone 2014). Not shown; see figure 8D for similar example.

Orthotriaenes: With long straight rhabds but strongly recurved clads. Smaller spicules, which are potentially immature, lack the recurved clads. Rhabds 1238–3795–5342 (n=21) × 69–127–283 μm (n=34); clads 161–403–599 (n=40) × 62–112–155 μm (n=12). Holotype also has strongly recurved clads, but many had one or more clads reduced to rounded nubs (not seen in the new sample); previously described with rhabds 2380–7320 × 180–220 µm, clads up to 480 × 200 µm (Lehnert & Stone 2014).

Dichotriaenes: Relatively small and much less common than orthotriaenes. Only one seen, rhabd 362 × 29 μm; clads 190 × 26 μm. Holotype previously described as having rhabds 780–930 × 85–98 µm, clad sizes not given (Lehnert & Stone 2014).

Anatriaenes: One example seen, with a broken rhabd; the remaining portion was 3841 × 44 µm; clad 120 × 41 µm. Holotype previously described as having rhabds 9800 × 47–58 µm; clads 82–90 × 42–55 µm (Lehnert & Stone 2014).

Acanthose oxyasters: With many (>8) conical, pointed rays; completely covered in small spines. Diameters 11–15–22 μm (n=40). Holotype previously described with diameters 10–14 µm (Lehnert & Stone 2014).

#### Distribution and habitat

Only two samples are known: one from Unalaska Island, in the Aleutians, and one from Malcom Island, British Columbia. The Aleutian sample was collected from 177 m; the depth is not recorded for the British Columbia sample, but it is labeled as coming from the "Cook Glass Sponge Reef Community".

### Remarks

The sponge examined here, from British Columbia, is a good morphological match to the only previously known sample of this species, with very long megascleres, recurved triaenes, small dichotriaenes, and only one type of aster. Some differences are also apparent: the holotype had a high frequency of reduced clads, which was not seen in the new sample. Genetic data to support conspecificity is desirable, but was not attempted here, as the holotype is in the ZSM collection in Germany.

The molecular phylogenies indicate that *S. makushina* falls into a clade comprised of *S. estrella*, *S. cardenasi* sp. nov., and species from Hawaii, the Atlantic, and the Mediterranean (figures 4, S1, S2). This clade may include the type species *S. grubii* Schmidt, 1862, based on a sequence from a Portuguese sample (figure S1). *S. makushina* is the sister species of *S. estrella*, with 0.8% divergence at the cox1 locus. Outside of the *Penares* species described below, all other species in this report are more closely related to species from the Atlantic or Mediterranean than to other species in the Northeast Pacific.

### *Stelletta clarella* de Laubenfels, 1930

Figure 12

**Figure 12.**
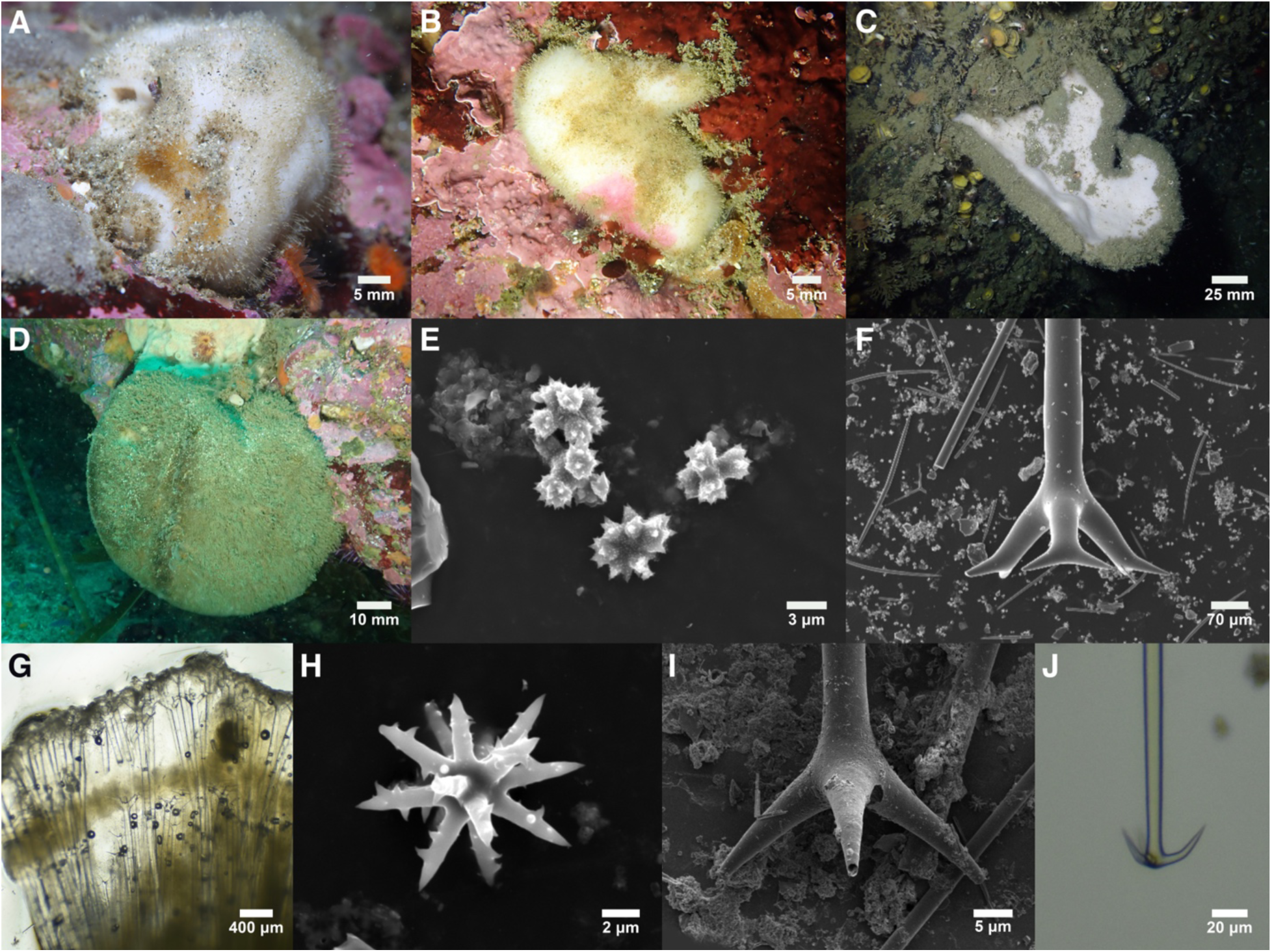
Stelletta clarella. A: TLT569 in situ (photo by Steve Lonhart, NOAA). B: TLT459 in situ. C: TLT1676 in situ. D: TLT1165 in situ. E: Anthasters. F: Dichotriaene. G: Section perpendicular to surface showing two layers of dichotriaenes; note that plagiotriaenes are slightly deeper in the sponge, perhaps because they are immature. H: oxyaster. I: plagiotriaene. J: anatriaene. E-H from TLT1139, I from TLT1555, J from TLT459.

#### Material examined

TLT459, Middle Reef, Point Lobos, California (36.52172,-121.93894), 6-15 m, 23-Nov-2019; TLT569, Middle Reef, Point Lobos, California (36.52172,-121.93894), 6-15 m, 23-Nov-2019; TLT1157, CRABS, California (36.55377,-121.93840), 10-17 m, 21-Sep-2021; TLT1165, Tower house arches, California (36.56187,-121.95950), 9-21 m, 21-Sep-2021; TLT1139, Inner Pinnacle, California (36.55852,-121.96820), 10-24 m, 22-Sep-2021; TLT520, Goalpost, California (32.69438,-117.26860), 12-15 m, 8-Feb-2020; CASIZ 302625, Favorite Channel, Alaska, (58.49117,-134.84983), 24 m, 11-Sep-2010; TLT1555, Aaron Island, Alaska, (58.44080,-134.82440), 24-43 m, 26-Jun-2025; TLT1576, Bird Island, Alaska, (58.49035,-134.85168), 24-41 m, 26-Jun-2025; TLT1580, Bird Island, Alaska, (58.49035,-134.85168), 24-41 m, 26-Jun-2025; TLT1676, Bird Island, Alaska, (58.49035,-134.85168), 24-41 m, 26-Jun-2025; TLT1677, Bird Island, Alaska, (58.49035,-134.85168), 24-41 m, 26-Jun-2025.

#### Morphology

White alive and post-preservation; some samples bear pink spots or yellowish areas that are likely due to algal colonization. Sponges are densely spiculated, firm and unyielding, with a cartilaginous cortical layer 1–2 mm thick at the ectosome. Oxeas project through the ectosome to form a thick spicule fur over the entire sponge surface, up to a centimeter thick in some samples. This extremely hispid surface traps sediment and debris in some habitats, leading to a brown appearance in situ. Samples collected in California were up to 5 cm thick, with unmeasured samples likely thicker, and were found as both amorphous, spreading encrustations and more compact, globular or subspherical shapes (figure 12D). One Alaskan sample was also roughly spherical, but others from Alaska formed concave blades approximately 1 cm thick and up to at least 10 cm long that grew outward from rock walls (figure 12C). An additional sample was blade-like but hanging vertically, with a stretched-looking aspect and subspherical areas at the tips. Oscula were not clearly evident on any sample, but field photos of one appear to show a depressed area with a cluster of small pores that may be oscula.

#### Skeleton

Choanosome densely spiculated with upright oxeas. Triaenes are found at the ectosome, in two layers, with the cladomes facing out. One layer is immediately under the sponge surface, while the other is just below the cortex. Cladomes of surface dichotriaenes sometimes protrude slightly, but spicule fur is mainly comprised of oxeas. Choanosomal oxeas average thicker than oxeas forming the spicule fur. Asters are dense at the surface, but also present throughout. Many foreign spicules are often present in the external spicule fur, which traps considerable debris.

#### Spicules

Shown in figure 12 except where noted.

Oxeas: tapering to sharp points at both ends, though rare styles are also seen. Thick oxeas are slightly fusiform, while thin oxeas are of consistent thickness until tapering at tips. Oxeas are abundant in all samples, but vary greatly in length and width within and between samples.

Originally described as having larger and smaller size classes (de Laubenfels 1932). However, bimodality in thickness was apparent only in some samples, with size classes that were inconsistent across samples, and lengths were not bimodal in any samples. When all measured spicules are pooled, they are 543–3056–11628 × 5–35–96 μm (n=204). Mean lengths per sponge varied from 1356–7169 μm, and this was significantly correlated with latitude (Pearson p=0.0085). Length may be affected by sponge size or other factors as well, as two samples from within Carmel Bay, Central California had non-overlapping distributions of length (TLT459 = 543–1701–2971; TLT1139 = 3881–4913–6118 μm). Type material previously described as having two size classes, 3500 × 50 μm and 1400 × 15 μm (de Laubenfels 1932). Not shown in figure 12; see figure 8D for similar example.

Dichotriaenes: The primary triaene spicules. With long rhabds and clads that tend to angle slightly downwards from the rhabd, but then bend and flatten out orthogonal to rhabd.

Occasional strongylote clads or rhabds are seen. Rhabds 1172–3202–5671 (n=51) × 26–78–137 (n=71) μm; clads 65–235–523 (n=112) × 10–51–124 (n=42) μm. Variable in size across samples, with mean rhabd lengths of 2170–4678 μm, mean clad lengths of 132–414 μm per sponge.

Triaenes were originally described as "orthotriaenes to plagiotriaenes to dichotriaenes", and it is true that they exist on a spectrum (de Laubenfels 1932). Here, I divide them into those with forked clads (dichotriaenes), which are the most abundant type, and those without (plagiotriaenes), which are less common and tend to be smaller. Type material previously described with rhabds 2000–3000 × 20–100 μm; clads 120–180 μm (de Laubenfels 1932).

Plagiotriaenes: Similar to dichotriaenes but usually smaller. The clads are usually straight, in contrast to the bent dichotriaene clads. Transitional examples seen, with a small forked "toe" on clad tips, or with forks on only some clads. Rhabds 235–1286–3944 (n=26) × 7–36–132 (n=33) μm; clads 22–110–531 (n=51) × 5–23–79 (n=29) μm. Grouped with dichotriaenes in type description (de Laubenfels 1932).

Anatriaenes: Only present in some samples. Rhabds 1047–2792–8471 (n=9) × 7–13–32 (n=12) μm; clads 15–51–77 (n=20) × 6–13–32 (n=8) μm. Type material previously described as 1100–2000 × 9–15 μm, clads 45–90 μm (de Laubenfels 1932).

Oxyasters: With many (>10) conical rays, usually ending in sharp points; those examined with SEM were sparsely acanthose or unspined. Diameters 8–11–16 μm (n=69). One sample (CASIZ 302625) is excluded from this distribution because it was an outlier in terms of aster size, possibly because it was contaminated with asters from another sponge (the spicule fur of this species collects considerable debris). This sample had asters with diameters 10–19–36 μm (n=27) (other spicules, and DNA sequences, were congruent with other samples). Type material previously described as 9–15 μm (de Laubenfels 1932).

Anthasters: Small asters that look like irregular sand grains in a light microscope. Possessing few rays which flare into spiny ends. Diameters 3–5–8 μm (n=126). Mentioned in the type description as "small siliceous structures" of uncertain affinity, but documented with SEM in later work (Lee *et al*. 2007).

#### Distribution and habitat

Known from Southern Alaska to Southern California, from the low intertidal to at least 41 m depth. Common at diving depths in Central California, where it is the only known species of *Stelletta*. Not common at diving depths in Southern California, where it grows alongside multiple other species of *Stelletta*; only 1 of 17 Southern California *Stelletta* collected by the author proved to be this species. It also appeared common in fjords around Juneau, Alaska, growing alongside *S. ovalae*. All samples examined for this report were subtidal, but the original species description was based on intertidal samples from Central California, and photos posted to iNaturalist.org appear to show this species at scattered intertidal locations across California and British Columbia. Previously reported from below 91 m (Green & Bakus 1994), but anthasters were not recorded in that sample, so it was likely a different species of *Stelletta*.

#### Remarks

This species was described from intertidal samples from Carmel Bay, Central California (de Laubenfels 1932). Freshly collected samples examined here include one from Carmel Bay, very near the type locality, and several from nearby locations around the Monterey Peninsula. These agree very well with the type description, and no other *Stelletta* have been found from California that contain large dichotriaenes or anthasters.

Lengths and widths of some spicule types varied greatly in this species (see spicule section above). This was correlated with latitude (Pearson p=0.0085), so environmental conditions may play a role. In the family Tetillidae, some species were found to have spicule sizes that were correlated with sponge size (Turner 2026). The sizes of most samples investigated here were not recorded, and only fragments were collected, so this correlation was not tested here. However, casual observation of approximate sponge size appears likely to be correlated with oxea length, so this could be tested in future work.

*Stelletta anthastra* Lehnert & Stone, 2014 was recently described from the Aleutian Islands, Alaska, and was said to be the first sponge known from the Northern Hemisphere to possess anthasters (Lehnert & Stone 2014). The authors were likely unaware that *S. clarella* has anthasters because these spicules were described as "small siliceous structures" that were likely foreign by de Laubenfels (1932). Later work (Lee *et al*. 2007), and the images shown here, clearly document anthasters in *S. clarella*. Additionally, a partial sequence of cox1 from the holotype of *S. anthastra* is available (Erpenbeck *et al*. 2015), and it is identical to sequences from *S. clarella*. It is therefore possible that *S. anthastra* will prove to be a junior synonym of *S. clarella*, but this is not proposed here for two reasons. First, cox1 evolves slowly in sponges, and doesn’t always differentiate apparently good species (Huang *et al*. 2008; López-Legentil *et al*. 2010; Pöppe *et al*. 2010; Turner & Pankey 2023). Second, there are several morphological differences between these species, most notably the triaenes, which are primarily dichotriaenes with flat clads in *S. clarella* but ortho- to plagiotriaenes with strongly recurved clads in *S. anthastra.* Other than *S. anthastra*, *S. clarella* was not closely related to any other species with DNA data available at cox1 or 28S (figures 4, S1, S2). It appears to be more closely related to *Tethyopsis mortenseni* and *Disyringa dissimilis* than it is to the type species of *Stelletta*.

### *Stelletta estrella* de Laubenfels, 1930

Figure 13

**Figure 13.**
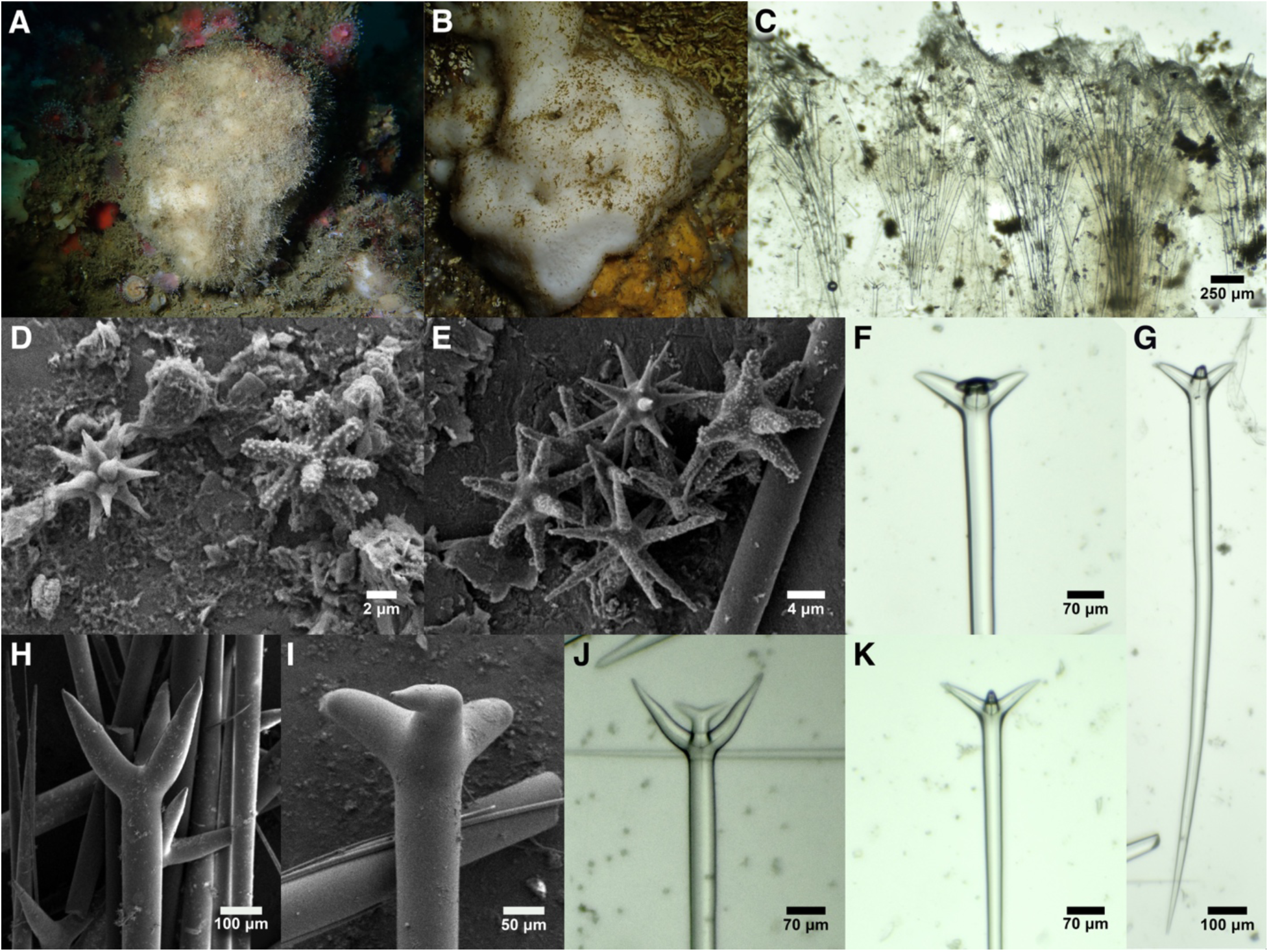
Stelletta estrella. A: TLT286 in situ. B: TLT1343 in situ. C: Perpendicular section at the sponge surface showing disorganized bouquets of plagiotriaenes from TLT1343. D: Unspined oxyaster (left) and acanthose strongylaster (right) from holotype. E: Aster variation in TLT286. F-K: Variability in plagiotriaenes; while G, H, K are the most typical forms, shapes are highly variable within and between samples. F, G, I, K from holotype, H, J from TLT286.

#### Material examined

Holotype: USNM 21399, Southern California, depth unknown, 10-Jul-1926. Other samples: TLT286, Hawthorne Reef, Los Angeles, California, (33.74714, - 118.42090), 18-24 m, 23-Aug-2019; TLT1343, Garbage Cove, Anacapa Isl., California, (34.01706, -119.36391), 2-12 m, 28-Aug-2024; TLT466, Parson’s Landing, Catalina Isl., California, (33.47502, -118.55000), 8-17 m, 10-Nov-2019.

#### Morphology

White cortex and yellowish-white choanosome in life, and these colors do not change with preservation. The cartilaginous cortical layer is about 1 mm thick, unyielding, and hard to cut through, but the choanosome is soft, somewhat yielding, and moderately densely spiculated. Hispid with protruding oxeas to various degrees, from a dense spicule fur to only microscopically hispid. Furred samples trap sediment and debris, leading to a brown appearance in situ. Up to 5 cm thick, irregularly massive to globular/subspherical. Oscula not evident on most samples, but field photos of one appear to show a slightly depressed area with a cluster of small pores that are likely oscula (figure 13B).

#### Skeleton

Choanosome moderately spiculated with oxeas, mostly perpendicular to surface but somewhat disorganized. Triaenes are found only near the surface, as bouquets, with their cladomes facing out. Some clads are immediately below the surface, while others are up to 2 mm inside the sponge. Some cladomes protrude slightly beyond the surface, but most protruding spicules are oxeas. Asters are dense at the surface, but also present throughout.

#### Spicules

Shown in figure 13 except where noted.

Oxeas: tapering to sharp points at both ends, though rare styles and strongyloxeas are seen. Some samples show a bimodal distribution in oxea width, while others do not; all are considered together here due to the high variability within and between samples. When all measured spicules are pooled, they are 465–2123–4781 × 5–38–105 μm (n=111). Mean lengths per sponge varied from 1700–2457 μm, and this was likely related to sponge size, though this was not explicitly tested. Holotype 1392–2463–3332 × 12–68–103 μm (n=20). Not shown in figure 13; see figure 8D for similar example.

Plagiotriaenes: Triaenes are primarily plagiotriaenes with long rhabds and clads that are straight or slightly curved. Smaller spicules have sharply pointed triangular clads, while larger spicules have clads with a conical, somewhat inflated aspect (widest slightly after diverging rather than where they meet the rhabd; see figure 13H). Some have clads that begin as plagiotriaenes but then bend and flatten out into a "foot", orthogonal to the rhabd; some of these are slightly forked at one or more clads and therefore transitional to dichotriaenes. Occasional strongylote clads or rhabds are seen, and a few have two or four clads. Rhabds 408–1290–2624 (n=142) × 8–45–105 (n=148) μm; clads 20–139–335 (n=227) × 8–43–79 (n=60) μm. Variable in size across samples, with mean rhabd lengths of 901–1707 μm and mean clad lengths of 106–176 μm per sponge.

Holotype rhabds 920–1707–2413 (n=25) × 21–68–105 (n=31) μm; clads 49–124–188 μm (n=28). Type material originally described as having rhabds 4000 × 35–200 μm (de Laubenfels 1932). I searched extensively for triaenes this large and did not find them on the slides prepared by de Laubenfels or in new spicule preparations from the holotype.

Highly acanthose oxyaster/strongylasters: With many (>10) rays completely covered in small spines. Rays vary from conical and pointed to strongylote and blunt; the majority are at least somewhat conical and tapering but with rounded tips. Only separated from unspined asters under SEM. Diameters 8–14–26 μm (n=49). Holotype alone: 8–12–26 μm (n=26).

Unspined oxyasters: Non-acanthose asters, less common than acanthose asters. Distinctly oxyote, with many (>10) conical and sharply pointed rays. Diameters 8–13–20 μm (n=40). Holotype alone: 8–11–20 μm (n=22).

#### Distribution and habitat

Only known from Southern California. There is a strong temperature gradient within Southern California (Watson *et al*. 2011), and this species was only found at the warm end of this gradient. Recently collected samples were all from the shallow subtidal, 2–24 m deep. Uncommon, it was found at only 3 of the 92 Southern California locations recently investigated by the author. Several samples were collected near where *S. cardenasi* sp. nov. was also found; in one case, they were found in the same submarine cave. The original description states that it is also found intertidally at Laguna Beach, but these samples have not been reexamined and could be either *S. estrella* or *S. cardenasi* sp. nov.

#### Remarks

In his monographic work on the sponges of California, de Laubenfels described two *Stelletta*: *S. clarella* in Central California, and *S. estrella* in Southern California (de Laubenfels 1932). Later, it was suggested that these were Northern and Southern varieties of the same species (Bakus & Green 1987). Genetic data presented here indicates that these species are only distantly related. *S. estrella* occurs in a well-supported clade together with *S. makushina, S. limuwensis* sp. nov., several recently described Hawaiian species (Nunley *et al*. 2025), and *S. dorsigera* Schmidt, 1864 from the Mediterranean (figure S2). This clade also contains a sample collected in Portugal by Joana Xavier, and identified as the type species *S. grubii* by Paco Cardenas (Cárdenas *et al*. 2011)(figure S1). Note that other samples identified as *S. grubii* from Northern Ireland, and sequenced at 28S, appear to be in a different clade (figure S2).

Attempts to sequence DNA from the holotype of this species were unsuccessful. With the discovery of several additional *Stelletta* in Southern California, care was taken to associate this existing name to the correct clade. *S. clarella* is easily distinguished by the presence of anthasters and large dichotriaenes. *S. nicolenya* sp. nov. and *S. limuwensis* sp. nov. are differentiated by having 3 types of asters, abundant dichotriaenes, and smaller triaenes. *S. cardenasi* sp. nov. and *S. estrella* are quite similar, but several features distinguish them. The large size of the triaenes was considered a defining feature of *S. estrella* by de Laubenfels; though I did not find any as large as claimed in the species description, the *S. estrella* holotype has rhabds averaging 1700 μm, with maximum lengths over 2400 μm. These spicules were variable in size among the other *S. estrella*, but one sample averaged 1440 μm and two had maximum values exceeding 2000 μm. In contrast, the longest spicules among eight *S. cardenasi* samples averaged 1014 μm and all 182 rhabds measured were under 2000 μm. Additionally, all *S. cardenasi* were externally gray in situ and the color was retained after preservation, in contrast to the *S. estrella* species description, holotype, and all other samples of *S. estrella*, which were entirely white before and after preservation. Finally, the acanthose asters in *S. cardenasi* sp. nov. appear to be slightly more oxyote and less strongylote than the *S. estrella* holotype, while the other *S. estrella* sample examined with SEM had a mix of oxyote and strongylote asters, like the holotype. Combined, the sponge color, triaene shapes and sizes, and aster shapes and sizes provide multiple lines of evidence which all point to the same genotype as being associated with the name *S. estrella*, leading to confidence that it has been assigned correctly. Nonetheless, these sympatric species are best distinguished using a combination of morphological and genetic data.

The exact holotype locality was not recorded for *S. estrella*, but the other samples analyzed by de Laubenfels were from Laguna Beach and near San Pedro and Long Beach, off Los Angeles. Sample TLT286 was collected off San Pedro, and was the most morphologically similar to the holotype of all samples examined; genetic data from this sample is therefore the best representation of this species’ genotype.

### Stelletta cardenasi sp. nov

Figure 14

**Figure 14.**
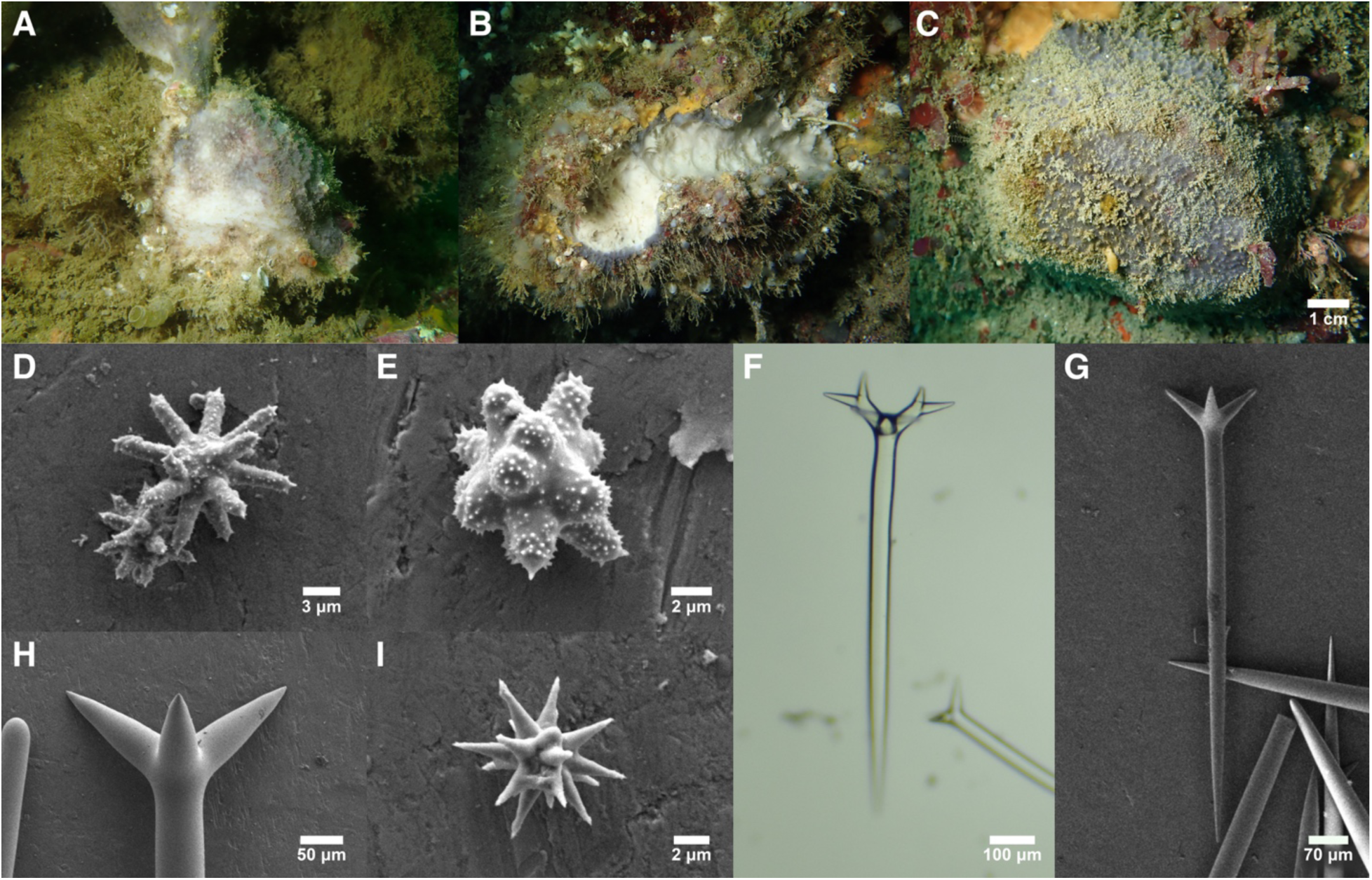
Stelletta cardenasi. A: Holotype in situ. B: TLT391 in situ; note that this is one of several examples of this species that were heavily fouled with other organisms, excepting, in this case, an injury that may have been caused by predation. C: TLT521 in situ. D: Typical acanthose asters. E: Atypical acanthose aster. F: Dichotriaene from TLT270. G-H: Typical plagiotriaenes. I: Unspined oxyaster. All SEM images are from holotype.

#### Material examined

Holotype: TLT352 (NHMLA 16524), Resort Wall, Los Angeles, California, (33.76499, -118.42815), 5-14 m, 23-Aug-2019. Paratype: TLT521, Goalpost, San Diego, California, (32.69438, -117.26860), 12-15 m, 8-Feb-2020. Other samples: TLT270 and TLT271B, Miracle Mile, San Miguel Isl., California, (34.01367, -120.34180), 5-20 m, 25-Aug-2019; TLT776A and TLT786B, Bird Rock, San Diego, California, (32.81447, -117.27431), Intertidal, 10-Jan-2021; TLT391, Saddleback Ridge, Santa Cruz Isl., California, (34.03817, - 119.52470), 5-12 m, 18-Oct-2019; TLT1459B, Garbage Cove, Anacapa Isl., California, (34.01706, -119.36391), 2-12 m, 28-Aug-2024; CASIZ 302848, Three Mile Bank, San Nicolas Isl., California, (33.30817, -119.45783), 27 m, 28-Nov-2011.

#### Etymology

Named for Paco Cardenas, Curator, Museum of Evolution, Uppsala University, in honor of his substantial contribution to sponge taxonomy and systematics, especially in the Tetractinellida.

#### Morphology

Most samples are completely gray externally, but some are mottled gray and white; all are white internally, and colors do not change with preservation. Surface is conulose due to projecting bundles of spicules across the surface. A mesh of small pores is visible between conules in macro photos of living specimens. Sponges are hard, unyielding, and densely spiculated both externally and internally, but especially on the surface, which consists of a cartilaginous cortical layer ∼1 mm thick that is difficult to cut through. Samples are up to 6 cm across and 3 cm thick, irregularly massive to subspherical. Oscula not apparent. Samples were lightly to heavily fouled by other demosponges, tunicates, hydroids, calcareous sponges, or other animals and algae. Several samples appeared to be entirely overgrown by other demosponges, such that the *Stelletta* was only discovered because a sample of the epizooic sponge was collected, and the lower sponge was revealed to be its substrate.

#### Skeleton

The choanosome is densely spiculated with oxeas perpendicular to the surface. Triaenes are mostly found in bouquets that fan out just below the surface, with their cladomes facing out. Some clads are immediately below the surface, while others are up to 2 mm inside the sponge. Some cladomes protrude slightly beyond the surface, but most protruding spicules are bundles of oxeas. Asters present throughout.

#### Spicules

Shown in figure 14 except where noted.

Oxeas: tapering to sharp points at both ends, though rare styles and strongyles are seen. A large range in length and width is seen within samples, but the distribution is not bimodal, nor are spicules significantly different in size when isolated from the ectosome vs. choanosome. When all measured spicules are pooled, they are 427–1980–4619 × 5–42–110 μm (n=143). The range in mean lengths per sponge was 1607–2406 μm. Holotype was 1002–2007–2656 × 14–53–74 μm (n=39). Not shown in figure 14; see figure 8D for similar example.

Plagiotriaenes: Triaenes are primarily plagiotriaenes with long rhabds and clads that are straight or slightly curved. Smaller spicules have sharply pointed triangular clads, while larger spicules have clads with a conical, somewhat inflated aspect (widest slightly after diverging rather than where they meet the rhabd). Some have one or more clads that begin as plagiotriaenes but they then bend and flatten out into a "foot" orthogonal to the rhabd; these are transitional to dichotriaenes. Occasional strongylote clads or rhabds are seen, and a few have two or four clads. Rhabds 406–920–1921 (n=182) × 9–39–76 (n=182) μm; clads 22–119–297 (n=253) × 13–38–58 (n=49) μm. Mean rhabd length per sample varied from 797–1014 μm, mean clad lengths varied from 97–143 μm per sponge. Holotype rhabds 406–928–1237 × 17–45–66 μm (n=63); clads 49–123–173 μm (n=72).

Dichotriaenes: Seen in three of eight samples, where they were less common than plagiotriaenes. With long rhabds and clads that tend to angle slightly downwards from the rhabd, then bend and flatten out orthogonal to the rhabd. Rhabds 680–811–968 × 24–42–66 μm (n=17); clads 85–142–215 μm (n=19). Not seen in the holotype.

Highly acanthose oxyaster: With many (>8) rays completely covered in small spines. Rays are generally pointed but vary from conical and tapering to nearly straight; a few have rays reduced to rounded nubs. Only separated from unspined asters under SEM. Diameters 9–13–19 μm (n=33); measured from SEM images in holotype. When asters were measured in other samples under light microscopy, they could not be separated into acanthose and non-acanthose, but they measured 4–10–23 μm (n=101).

Unspined oxyasters: Non-acanthose asters. Rare. Distinctly oxyote, with many (>10) conical and sharply pointed rays. Diameters 9–13–21 μm (n=3); measured from SEM images in holotype.

#### Distribution and habitat

Only known from Southern California, but in contrast to *S. estrella*, it was found throughout this region. Collected in both the intertidal and the subtidal up to 27 m depth. It may be more common at the offshore Channel Islands, where it was found at 8% of sites recently investigated by the author, including four of the six islands surveyed, versus on the mainland, where it was found at only two locations (5% of sites).

#### Remarks

This species is in the same clade as *S. estrella* in both phylogenies, but they do not appear to be closely related. In the 28S tree, for example, *S. cardenasi* sp. nov. is sister to *S. dorsigera* from the Mediterranean (figure S2). Without genetic data, *S. cardenasi* sp. nov. would likely be lumped with *S. estrella* due to the similarity of their spicules, and it is possible that some samples de Laubenfels assigned to *S. estrella* were actually this species (only the holotype was reexamined). These species can be differentiated morphologically using their external color, the sizes of the triaenes, and the morphology of their asters (also see the dichotomous key and the *S. estrella* remarks). However, it is clear that there is high spicule variability in many astrophorids, so genetic confirmation is strongly suggested when differentiating future samples of these species.

*S. cardenasi* sp. nov. can be differentiated from the more northern species in the Eastern Pacific by having much smaller megascleres, and in the case of the one sympatric species *S. clarella*, the lack of anthasters. It is differentiated from *S. nicolenya* sp. nov. and *S. limuwensis* sp. nov. by having mainly plagiotriaenes rather than dichotriaenes and lacking a third class of aster. No other species are known from the Eastern Pacific. The spicular characteristics of all Northwestern Pacific species were compiled by Lehnert and Stone (2014), and are all differentiated qualitatively or quantitatively by their spicules from the new species; these are also unlikely congeners for biogeographic reasons. The only species described from the Pacific after 2014 were from tropical waters in Hawai’i, and these all have spicular differences compared to the species described here (Nunley *et al*. 2025).

One additional feature was useful in identifying this species: Sanger sequencing of all samples failed at 28S, with multiple sets of primers, despite success on all other *Stelletta* samples tested. The ribosomal locus of the holotype was successfully sequenced with the Illumina technique, and this sequence had no detected mismatches with the primers used. It is therefore likely that there were either primer mismatches specific to this species that were not detected in the Illumina sequence due to sequence errors, or there are problems caused by secondary structures at this locus that are found in only this species. All samples were successfully sequenced at the cox1 locus using the custom primers designed.

### Stelletta nicolenya sp. nov

Figure 15

**Figure 15.**
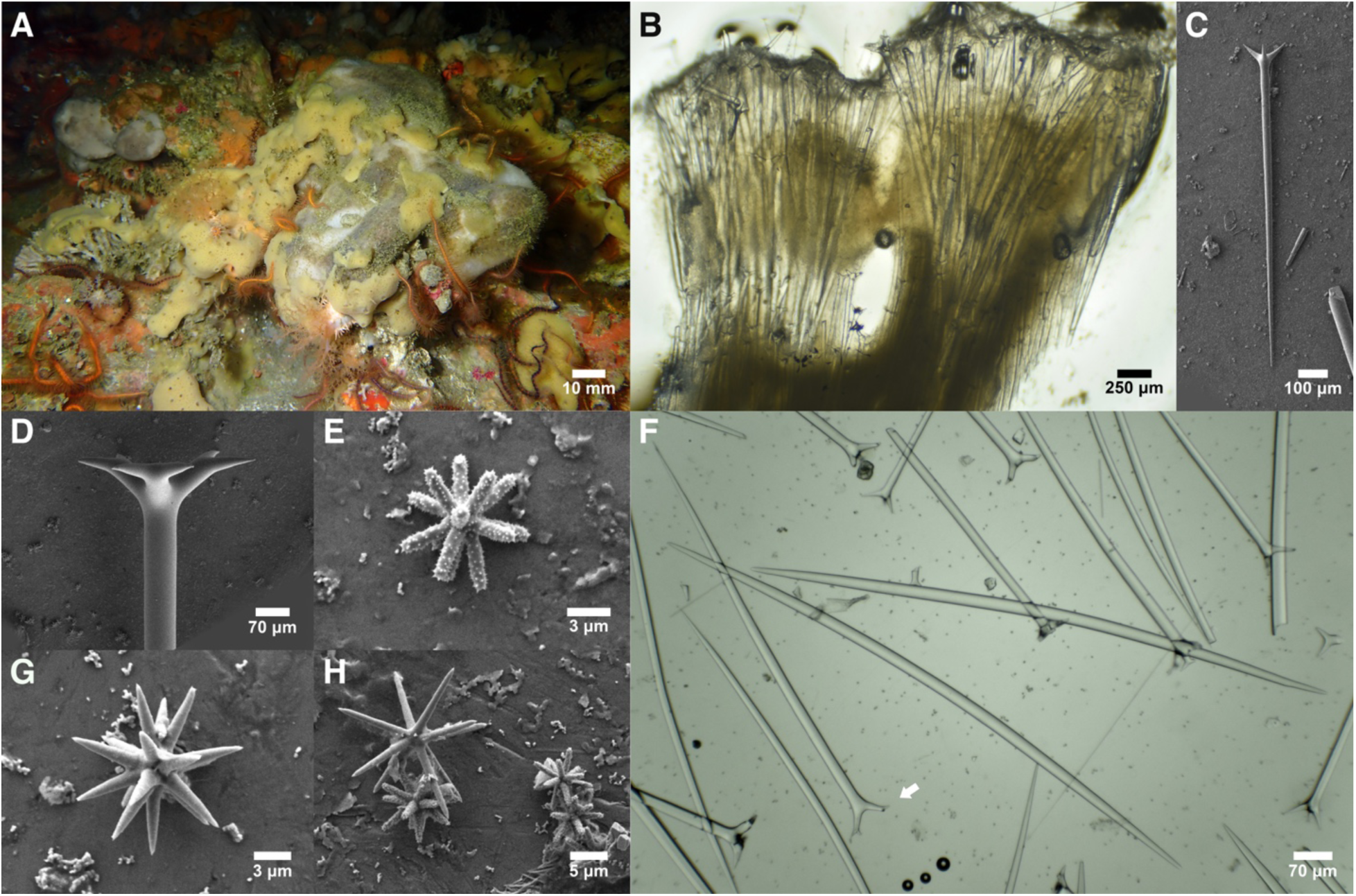
Stelletta nicolenya. A: In situ image; the large white and gray mottled sponge is the type sample, partially overgrown by a yellow Haliclona sp; a Penares sp. is also visible in the upper left. B: Perpendicular section at the sponge surface showing triaenes in bouquets at the surface with another layer below the cortex. C: Plagiotriaene. D: Dichotriaene. E: Acanthose stylaster. F: Oxeas and various triaenes, including one with two clads indicated by the white arrow. G: Small unspined oxyaster. H: Large unspined oxyaster with several stylasters.

#### Material examined

Holotype: TLT1492B, Dutch Harbor East, San Nicolas Isl., California, (33.21600, -119.48413), 10-14 m, 10-Oct-2024.

#### Etymology

Named in honor of the Nicoleño people who once inhabited San Nicolas Island.

#### Morphology

The only known sample was thickly encrusting, approximately 10 cm across at greatest extent; the sampled portion is 1 cm thick but the sponge was likely thicker alive. Mottled gray and white externally, white internally. Densely spiculose, hard and unyielding, with an especially hard ∼1 mm thick cartilaginous cortical layer. A dense fur of protruding spicules occurred in patches, with other areas only microscopically hispid. Oscules not seen. Partially overgrown by an unidentified *Haliclona*.

#### Skeleton

Choanosome densely spiculated with oxeas perpendicular to surface. Triaenes are found only within a few millimeters of the surface, with rhabds pointing internally. One layer of cladomes is found just below the surface, in flared bouquets, while another layer of cladomes is seen below the cortex 1.5 mm deeper into the sponge; only a few cladomes were seen between these layers. Some cladomes protrude slightly beyond the surface, but most protruding spicules are oxeas. Asters present throughout.

#### Spicules

Shown in figure 15.

Oxeas: tapering to sharp points at both ends. 1227–2042–2594 × 7–42–60 μm (n=14).

Dichotriaenes: Triaenes are primarily dichotriaenes with clads that angle away from the rhabd and then flatten orthogonally into a flat, forked foot. A large minority (nearly 10%) have only two clads. Rhabds 1071–1481–1846 × 35–47–59 μm (n=17); clads 81–117–188 μm (n=17).

Plagiotriaenes: Less common and slightly smaller than dichotriaenes, but difficult to separate due to transitional forms. Clads generally straight and triangular. Rhabds 848–1159–1373 × 26–33–39 μm (n=9); clads 88–106–174 μm (n=10).

Highly acanthose stylaster: The most common aster, with many (>10) rays completely covered in small spines. Rays are generally untapering with rounded ends, but some are conical and tapering and therefore transitional to oxyasters. Only separated from unspined asters under SEM. Diameters 7–10–13 μm (n=53).

Oxyasters: Non-acanthose or very lightly-spined asters with many (>10) conical, pointed rays. Size distribution is bimodal, so they are divided into size classes here. Small size class with diameters 7–13–16 μm (n=19); larger size class with diameters 18–23–30 μm (n=15).

#### Distribution and habitat

Known from a single sample collected from San Nicolas Island, Southern California, between 10 and 14 m depth.

#### Remarks

This species is in the same clade as *S. limuwensis* sp. nov in the 28S phylogeny, but it is more closely related to *S. mediterranea* (Topsent 1893) than to this sympatric species (figure S2). It is only distantly related to the other *Stelletta* found in California.

This species is morphologically similar to *S. limuwensis* sp. nov, but has larger megascleres and more strongylote acanthose asters. With only one sample from each species, the range of morphological variation is unknown, and genetic confirmation is strongly suggested when differentiating future samples of these species. *S. estrella* and *S. cardenasi* sp. nov. are also similar, but primarily have plagiotriaenes rather than dichotriaenes and lack the large size class of unspined oxyaster. *S. nicolenya* sp. nov. can be easily differentiated from the more northern species in the Eastern Pacific by having much smaller megascleres, and in the case of the one sympatric species *S. clarella*, lacking anthasters. No other species are known from the Eastern Pacific. Morphological data on Northwestern Pacific species were compiled by Lehnert and Stone (2014), and are all differentiated qualitatively or quantitatively by their spicules; these are also unlikely congeners for biogeographic reasons. The only species described from the Pacific after 2014 were from tropical waters in Hawai’i, and these all have spicular differences compared to the species described here (Nunley *et al*. 2025).

### Stelletta limuwensis sp. nov

Figure 16

**Figure 16.**
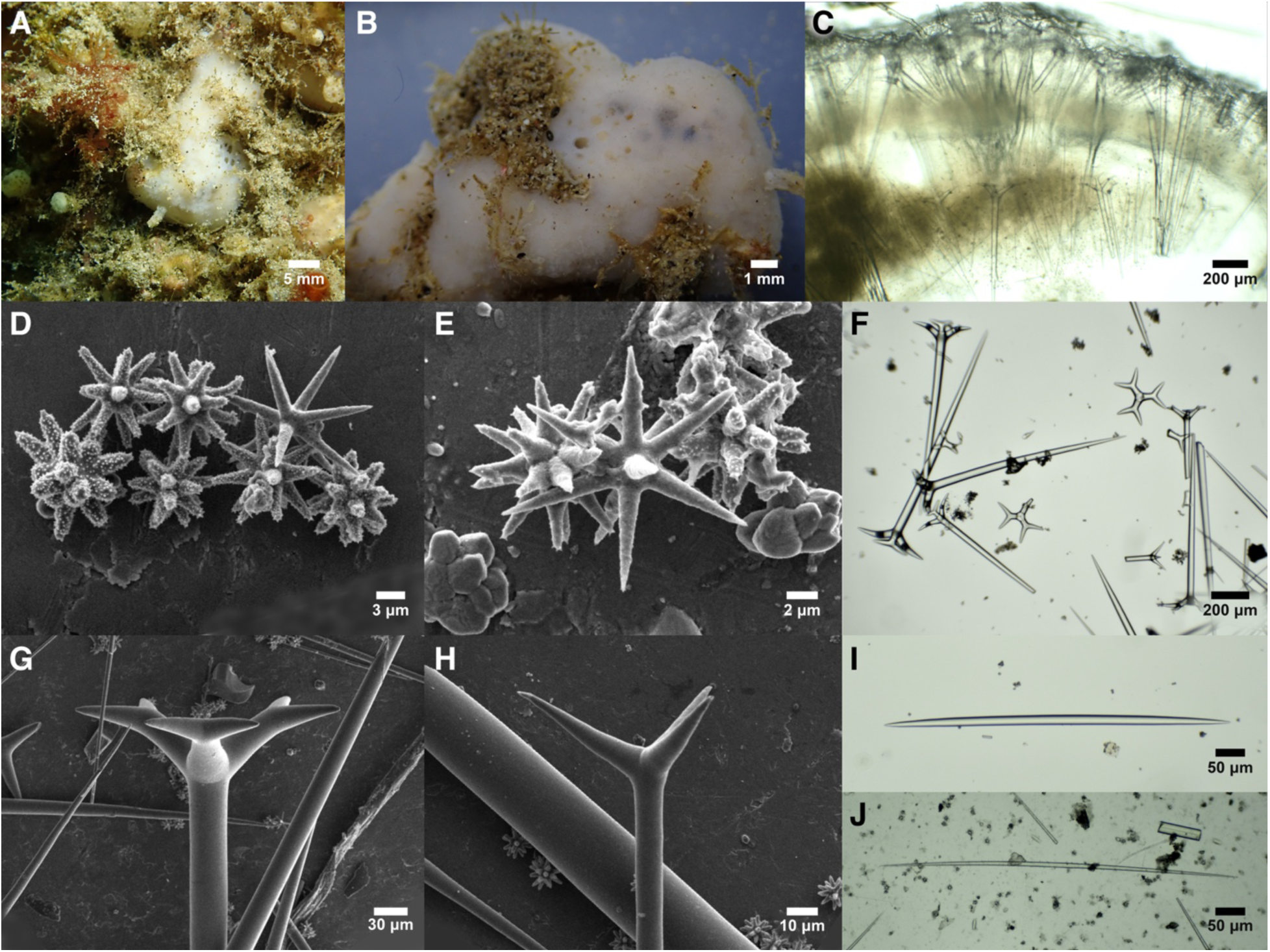
Stelletta limuwensis. A: In situ photo. B: Sample post collection, pre-preservation. C: Perpendicular section at the sponge surface showing two layers of cladomes. D: One large unspined aster (top right) with a variety of acanthose asters. E: Asters, from left to right: small unspined aster, large unspined aster, acanthose strongylaster. F: Typical triaenes. G: Dichotriaene cladome. H: Plagiotriaene cladome. I: Thick oxea. J: Thin oxea.

#### Material examined

Holotype: TLT329, Pink Ribbon, Santa Cruz Isl., California, (33.98967, - 119.52470), 3-11 m, 18-Oct-2019.

#### Etymology

Named inspired by ’Limuw’, the Chumash name for Santa Cruz Island, where the only known sample was found.

#### Morphology

A thickly encrusting sponge, 3 × 2 cm across and 2 cm thick. Rough to the touch but only microscopically hispid. White exterior and yellow-white interior before and after preservation. Cartilaginous cortical layer present but thin, approximately 0.5 mm thick. The choanosome is relatively soft and yielding, without the dense spiculation seen in some *Stelletta*. A cluster of oscula, each 0.5–1 mm across, were visible in the living sample and did not close after collection.

#### Skeleton

Choanosome with oxeas, mostly perpendicular to surface, but somewhat confused. Triaenes are found only within a few millimeters of the surface, with rhabds pointing internally. One layer of cladomes is found just below the surface, in flared bouquets, while another, less dense layer of cladomes is seen below the cortex approximately 1 mm deeper into the sponge. A few scattered oxeas protrude beyond the sponge surface. Asters are present as a dense surface layer, but also scattered throughout the interior.

#### Spicules

Shown in figure 16.

Oxeas: tapering to sharp points at both ends. Highly variable in size, with a bimodal length distribution and a trimodal width distribution. Considered together, 404–1020–1835 × 3–19–40 μm (n=144). If separated into a large and small size class, the large are 747–1259–1835 × 16–27–40 μm (n=77) and the small are 404–746–1195 × 3–10–14 μm (n=67).

Dichotriaenes: The most common triaenes are dichotriaenes, with clads that angle away from the rhabd and then flatten orthogonally into a flat, forked foot. Rhabds 414–982–1265 × 18–37–53 μm (n=32); clads 91–145–194 (n=33) × 23–31–39 (n=13) μm.

Plagiotriaenes: Less common and slightly smaller than dichotriaenes, but difficult to separate due to transitional forms. Clads generally straight and triangular. Rhabds 252–625–1100 × 6–20–50 μm (n=35); clads 20–71–193 (n=36) × 5–17–41 (n=27) μm.

Highly acanthose oxyaster/stylaster: The most common aster, with many (>10) rays completely covered in small spines. Rays are vary from conical to untapered and straight; ray tips are most commonly rounded but sometimes pointed. Only separated from unspined asters under SEM. Diameters 7–11–16 μm (n=64).

Oxyasters: Non-acanthose or very lightly-spined asters with many (>10) conical, pointed rays. The size distribution is bimodal, so they are divided into size classes here. The small size class has diameters 6–10–14 μm (n=16); large size class has diameters 16–20–27 μm (n=22).

#### Distribution and habitat

Known from a single sample collected from Santa Cruz Island, Southern California, between 3 and 11 m depth.

#### Remarks

This species is in the same clade as *S. nicolenya* sp. nov in the 28S phylogeny (figure S2), but it is more closely related to species from the Atlantic than to this sympatric species. It is only distantly related to the other *Stelletta* found in California. Despite attempting both Illumina and Sanger sequencing, data was not obtained from cox1.

This species is morphologically similar to *S. nicolenya* sp. nov, but has small megascleres and acanthose asters that are slightly more oxyote. With only one sample from each species, the range of morphological variation is unknown, and genetic confirmation is recommended when differentiating future samples of these species. *S. estrella* and *S. cardenasi* sp. nov. are also similar, but have primarily plagiotriaenes rather than dichotriaenes and lack the large size class of unspined oxyaster. *S. limuwensis* sp. nov. can be easily differentiated from the more northern species in the Eastern Pacific by having much smaller megascleres, and in the case of the sympatric *S. clarella*, lacking anthasters. No other species are known from the Eastern Pacific. Morphological data on Northwestern Pacific species were compiled by Lehnert and Stone (2014), and are all differentiated qualitatively or quantitatively by their spicules; these are also unlikely congeners for biogeographic reasons. The only species described from the Pacific after 2014 were from tropical waters in Hawai’i, and these all have spicular differences compared to the species described here (Nunley *et al*. 2025).

### Genus Dercitus

#### Definition

Ancorinidae with calthrops or dichocalthrops as megascleres and possessing irregular acanthomicrorhabd-like sanidasters with a thick central axis relative to the actines; further microscleres may include smooth toxa-like forms and aster-like compressed forms; no structural oxea megascleres. Emended from Van Soest *et al*. (2010).

### Subgenus *Stoeba*

#### Diagnosis

*Dercitus* with a single microsclere category in the form of irregular sanidasters (Van Soest *et al*. 2010).

### Dercitus (Stoeba) syrmatitus de Laubenfels, 1930

Figure 17

**Figure 17.**
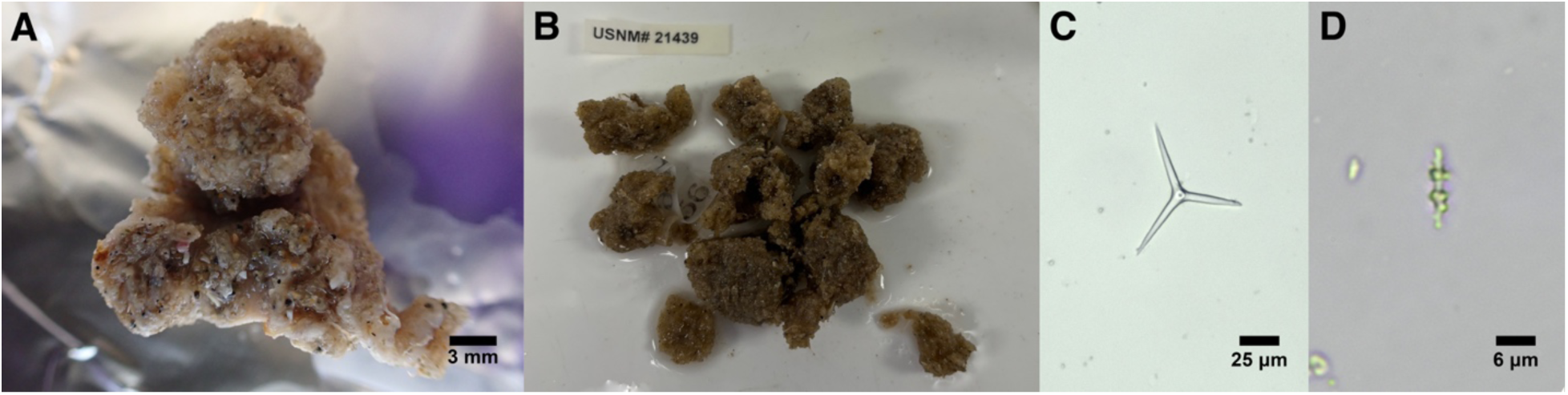
Dercitus syrmatitus. A: Portion of the holotype that was loaned. B: Entire paratype, photo provided by Abigail Reft (Smithsonian). C: Calthrop from holotype. D: Sandiaster from the paratype.

#### Material examined

Holotype (US 21438) and paratype (US 21439), Laguna Beach, California, intertidal, 14-Mar-1926.

#### Morphology

Described by de Laubenfels (1932) as drab and amorphous, with spicules occupying pockets 0.2–2 mm in diameter among sand, foreign spicules, and possibly a keratose sponge.

#### Skeleton

Undetermined.

#### Spicules

Shown in figure 17.

Calthrops: With three to four rays. I was unable to find any calthrops when new spicule preparations were made from small portions of the type samples, but measured those on de Laubenfels’ slides as 35–40–48 μm (n=9); reported as 25–65–80 × 3–8–10 μm by Lee et al. (2007); reported as 25–80 × 3–10 μm in original description.

Sandiasters: I was able to find only two in new spicule preparations, both from the paratype, 10 and 13 μm long. Reported by both de Laubenfels (1932) and Lee et al. (2007) as 8–12 μm.

#### Distribution and habitat

Known from only two samples, both collected in the intertidal zone at Laguna Beach, Southern California.

#### Remarks

The two type samples are the only ones known for this species. I attempted to sequence DNA from these and was unsuccessful. Isolation of new spicules was also mostly unsuccessful, probably because I did not wish to consume a large quantity of the type material, but the sponge is found only as small pockets among sand and possibly a keratose sponge (de Laubenfels 1932). Indeed, I isolated far more foreign spicules (chelae, sigmas, oxeas, etc.) than apparent *Dercitus* spicules. However, these samples were reexamined by Lee et al. (2007), who verified the dimensions reported in the original description, provided a mean value for clad lengths, and was able to acquire SEM images of both spicule types.

### Dercitus (Stoeba) giveni sp. nov

Figure 18

**Figure 18.**
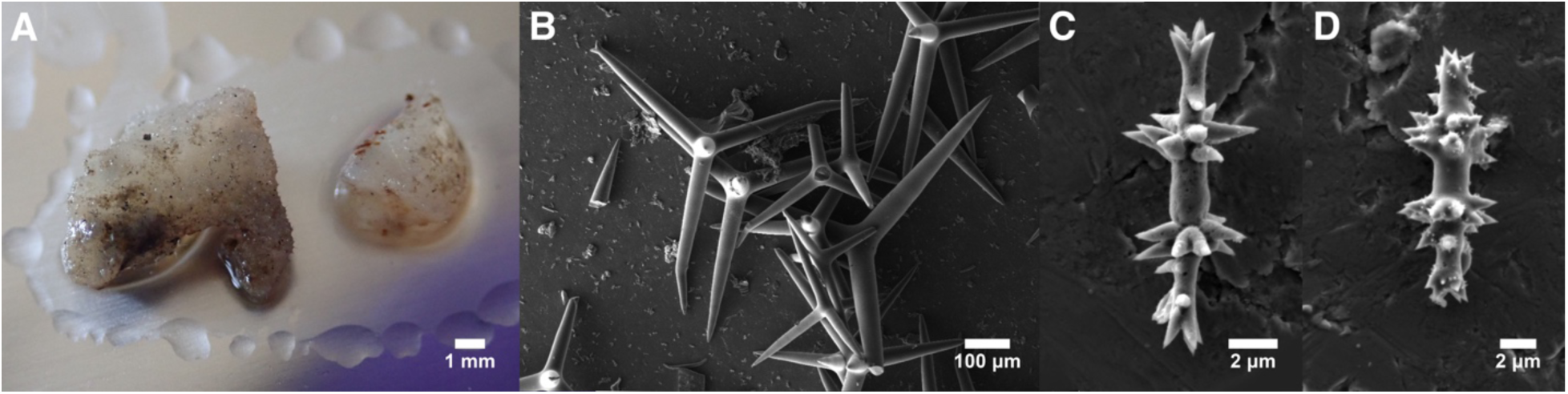
Dercitus giveni. A: Entire preserved sample. B: Calthrops. C-D: Sandiasters. All photos from holotype.

#### Material examined

Holotype (SBMNH 659287), San Elijo Lagoon, San Diego, California, 18 m, 9-Mar-1964.

#### Etymology

Named for Robert Given, the collector of the sample, who was a pioneer of scientific diving and the former president of the American Association of Underwater Scientists (AAUS).

#### Morphology

The only known sample consists of two small, partially transparent whitish fragments, the larger of which is 9 × 8 mm. Appearance in life unknown.

#### Skeleton

Undetermined.

#### Spicules

Shown in figure 18.

Calthrops: Variable in shape, but all appeared to have four rays. The majority were calthrop-shaped, but some had rays at more acute angles. Rays were most commonly straight, but one or more bent or forked rays is present in a large minority of spicules; more rarely, a strongylote ray is seen. 172–328–460 × 19–39–53 μm (n=42).

Sandiasters: 12–14–16 × 1–2–3 μm (n=19).

#### Distribution and habitat

Known from only one sample, collected by Robert Given near San Diego, Southern California. The museum voucher lists the location only as "San Elijo Lagoon". However, the collection date and previous sample identification as *D. syrmatitus* allow this voucher to be associated with a specific sample collected as part of a marine survey offshore of the lagoon (Turner *et al*. 1965). The sample was collected by divers in 18 m of water, in an area with low-lying rock shelves. The precise location was "150 yards north of transect line 1". By comparing the hand-drawn maps provided with satellite imagery, I estimate this location as (33.011246, -117.280434).

#### Remarks

This sample was acquired on loan because it was the only other sample I know of that was previously identified as *D. syrmatitus*. As I have been unable to amplify DNA from any of de Laubenfels’ samples, I hoped that I could place *D. syrmatitus* in the sponge phylogeny using this sample. I was thwarted for two reasons: first, I was unable to amplify DNA from this sample, either. Second, the calthrops of this sample are five times larger than those of *D. syrmatitus*.

Indeed, this sample has larger calthrops than any described sample of *Dercitus* (Calcinai *et al*. 2017; Santos & Pinheiro 2016; Van Soest *et al*. 2010). The only species with similarly sized calthrops is *D. senegalensis* Van Soest *et al*., 2010 from Sengal and Mauritania, which has calthrops 36–480. In addition to being an extremely unlikely conspecific for biogeographic reasons, the new species has a larger mean size and a smaller range of calthrop sizes than *D. senegalensis*, with a minimum clad length of 169 μm, rather than 36 μm.

### Family Geodiidae Gray, 1867

#### Definition

Astrophorina with microrhabds, spherules and a diversity of euasters. Sterrasters are a synapomorphy of this group, although they have been secondarily lost in some taxa. From Cárdenas *et al*. (2011).

### Genus *Geodia* Lamark, 1815

#### Definition

Geodiidae with euasters or large microrhabds in the ectocortex and sterrasters in the endocortex (sometimes secondarily lost). From Cárdenas *et al*. (2011).

### *Geodia angulata* (Lendenfeld 1910)

Figure 19

**Figure 19.**
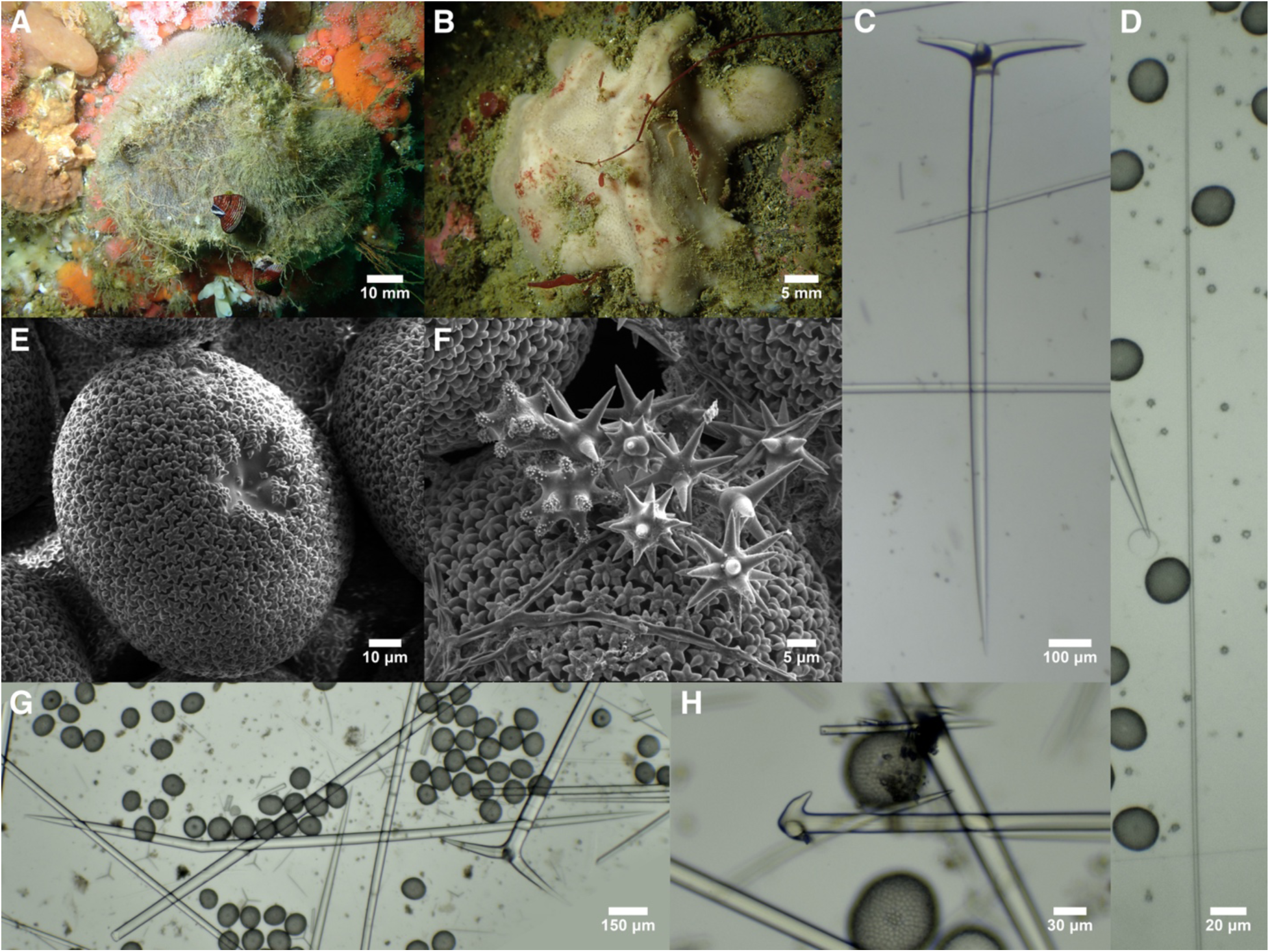
Geodia angulata. A: TLT1175 in situ. B: TLT576 in situ; note the lack of spicule fur. C: Plagiotriaene. D: Thin oxea II from TLT1175. E: Sterraster from USNM 8403. F: Acanthose strongylospherasters (left) and smooth oxyspherasters from USNM 8403. G: Bent oxea I that is the species namesake, from TLT1176. H: Anatriaene from USNM 8380.

#### Material examined

*G. angulata* var. *megana* syntype, USNM 8403, West of Anacapa Isl., California, (34.025, -119.483), 66 m, 12-Feb-1889; *G. angulata* var. *microana* holotype, USNM 8380, NE of Santa Barbara Isl., California, 53 m, 12-Apr-1904; *G. angulata* var. *orthotriaena* holotype, USNM 8402, Anacapa Isl., California, (34.000, -119.492), 55 m, 6-Feb-1889; TLT1175 & TLT1176, Fire Rock, Carmel Bay, California, (36.55898, -121.95110), 10-22 m, 10-Aug-2021; TLT576, Six Fathoms, San Diego, California, (32.71000, -117.26860), 9-18 m, 8-Feb-2020.

#### Morphology

Thickly encrusting, massive, or lobate, up to 112 mm across and 77 mm thick. The exterior is beige to gray alive, with a white interior. Large portions of the surface are decorated with fields of small pores, very uniform in size and arrangement, 0.2–0.5 mm across. Some individuals are covered with a spicule fur up to 8 mm thick, while others are not macroscopically hispid; intermediate to these, some have protruding spicules but only in places where the sponge grows to form a sheltered crevice. When copious spicule fur is present, areas with pore fields are less furry, and less fouled with other organisms, than other portions of the sponge surface. Pores remain open post-preservation. See Lendenfeld (1910) for additional details.

#### Skeleton

Not investigated here, but described in detail by Lendenfeld (1910). Briefly, the choanosome contains bundles of oxeas I extending to the surface. Near the surface, plagiotriaenes, oxeas II, and anatriaenes are added to bundles. The cladomes of the plagiotriaenes occur below the cortical layer at the surface, while the oxeas II and anatriaenes protrude beyond the sponge surface to form the spicule fur. The cortical layer contains the sterrasters. Oxyasters are reported to be limited to the choanosome, with oxyspherasters primarily in the inner cortex and strongylospherasters in the outer cortex.

#### Spicules

Shown in figure 19 except where noted.

Oxeas I: stout, somewhat fusiform, and tapering to sharp points at both ends; a small minority are styles. Some are sharply bent, which led to the species name, but bends are uncommon (and sometimes seen in other sympatric *Geodia*). Highly variable in size, with mean lengths per sponge 1486–2841 μm; the three syntypes examined average 1815, 2028, and 2841 μm. When spicules from all samples are pooled, they are 948–1997–3389 × 13–35–69 μm (n=182).

Reported as 1600–3700 × 20–72 μm in the type description (Lendenfeld 1910).

Oxeas II: differentiated from oxeas I by being thinner relative to their length and less fusiform. Because they pierce the surface, they are abundant in samples with spicule fur and rare in those without; no unbroken examples could be found in sample TLT576, with was unfurred in situ.

Variable in size across samples, with mean lengths per sponge of 3512–6971 μm. When spicules from all samples are pooled, they are 1800–4434–8848 × 6–21–32 μm (n=43). Reported as 2900–9500 × 5–34 μm in the type description (Lendenfeld 1910).

Short oxeas: short oxeas (300–400 μm) are abundant in syntype USNM 8380, but not in the other samples, and are presumed to be foreign. Not mentioned by Lendenfeld (1910).

Plagiotriaenes: with long rhabds, usually tapering to sharp points, but a few are short and strongylote. The clads are frequently unequal in length, and only the largest clad was measured. Rhabds 758–1992–2872 μm (n=72) × 18–57–95 μm (n=100). Mean lengths per sample 1569–2319 μm. Clads 97–352–659 (n=95) × 17–44–74 (n=107) μm. Reported with rhabds 1500–2800 × 47–82 μm and clads 330–700 μm in the type description (Lendenfeld 1910). The rhabd:clad angles were 98°–106°–115° (n=91), with means per sample of 104°–109°. This differs somewhat from angles reported by Lendenfeld, who stated they varied from 93.7° to 105.6° depending on the sample. Measuring these angles is not straight-forward because the clads are curved.

Anatriaenes: none were seen in the three freshly collected samples, despite doing multiple spicule preps from different regions of the sponges, and only 6 were seen in the new spicule preps from the three syntypes. The two unbroken rhabds are 3479–5103 μm long; all six rhabds are 5–15–18 μm wide. Clads are 24–42–62 × 5–14–19 μm, with clad:rhabd angles 33°–39°–43° (n=6). Lendenfeld (1910) reported rhabds up to 9000 μm long, 7–39 μm wide. Clad lengths were reported as variable, with those from var. *megana* 45–210 μm long, var. *orthotriaena* 33–80 μm long, and those of var. *microana* 30–50 μm long. Clad:rhabd angles were reported as 27°–47°–66°.

Sterrasters: Nearly round when seen from above, with length:width ratios of 1.1. Rosettes are not warty. When all samples are pooled, mature sterrasters (those with rays ending in rosettes) are 52–88–125 (n=466) × 51–83–110 (n=209) × 37–61–76 (n=53) μm. Considerable size variation is seen across samples, with mean lengths (maximum diameters) per sample of 74–112 μm. Reported as 85–122 × 75–113 × 57–86 μm in the type description (Lendenfeld 1910).

Oxyasters/oxyspherasters: All are completely smooth with no spines. The type description describes a continuous series from large oxyasters with no centrum to small oxyspherasters with a large centrum. SEM imaging of one syntype was consistent with this description. Oxyspherasters with a prominent centrum are the most common type, but some have no centrum. When attempting to differentiate oxyasers by type, oxyspherasters are 10–17–27 μm (n=63) in total diameter, while oxyasters are 14–24–38 μm (n=26), but categorization by type was fraught. When all oxyasters are pooled, they are 10–18–38 μm (n=101). Reported as 11–64 μm in the type description, which analyzed more samples and includes additional detail about shape and size (Lendenfeld 1910).

Strongylospherasters: with a large, smooth centrum and short, heavily spined, strongylote rays. In the syntype examined with SEM, total diameters were 11–17–20 μm (n=24). Reported as 7–26 μm in the type description, which analyzed more samples and includes additional detail about shape and size (Lendenfeld 1910).

#### Distribution and habitat

Known from Central and Southern California. It is found in shallow waters compared to other *Geodia* in the region, with the shallowest sample collected between 9–18 m depth, and the deepest at 66 m depth. The type material was dredged from several locations around Santa Barbara Island and Anacapa Island, Southern California. New samples were collected on SCUBA in kelp forest habitat in Carmel Bay, Central California and off Point Loma, San Diego. It is not widespread at diving depths within this range, as recent sponge collections have been made by the author at 113 locations across Central and Southern California, and it was found at only two of these sites.

#### Remarks

Though he had only four samples, Lendenfeld erected 3 varieties when describing this species: var. *microana* (with only small-cladded anatriaenes), var. *megana* (for two samples with larger maximum anatriaene clad lengths), and var. *orthotriaena* (with intermediate maximum anatriaene clad lengths, and with orthotriaenes instead of plagiotriaenes). I isolated and measured new spicules from three of Lendenfeld’s type samples, one from each variety (I also examined some of Lendenfeld’s archived slides, but these contained too few spicules to be of much use). Three new samples were also examined, providing greater context on spicule variability.

In most cases, the quantitative spicule data reported by Lendenfeld was a good match to the new spicules measured from the type samples. A notable exception was that I was unable to replicate differences among varieties in the shape of the plagiotriaenes, and therefore did not find support for var. *orthotriaena*. In Lendenfeld’s data, var. *orthotriaena* had rhabd:clad angles averaging 93.7°, while the other samples averaged 101.3°, 103°, and 105.6°. Lendenfeld did not specify how he measured the rhabd:clad angle, and the task is not straightforward because the rhabds are often curved throughout their length and/or with tips that curve back towards the rhabd axis. Despite trying several methods, I did not find that var. *orthotriaena* had lower angles than the other samples (var. *orthotriaena* mean 105°, var. *megana* mean 104°, var. *microana* mean 109°, new sample TLT1176 mean 107°). Though there are significant differences in angles among samples (ANOVA p= 4.91e-05), var. *orthotriaena* is not the lowest, nor did there seem to be categorical types worthy of erecting named varieties. The other notable spicule variation among Lendenfeld’s samples was in the lengths of the anatriaene clads, where longer values were found in the var. *megana* samples (var. *megana* 45–210 μm; var. *orthotriaena* 33–80 μm; var. *microana* 30–50 μm). Consistent with these data, I found only short-cladded anatriaenes in var. *orthotriaena* and var. *microana*; however, no anatriaenes were seen in the newly collected samples or the new spicule prep from var. *megana*. Lendenfeld describes these spicules as occurring within the spicule fur on the outside of the sponge, and there was little of this present on the subsamples loaned to me, and none on one of the newly collected samples. The two newly collected samples from Carmel had abundant spicule fur, but multiple spicule preps from different regions of the sponge still failed to find any anatriaenes. It seems likely that they are sometimes absent, or perhaps are limited to the basal region of the sponge, as the newly collected samples are fragments that do not include where the sponge was attached to the substrate.

Several other differences were found between var. *megana* and the other samples by Lendenfeld, and reproduced by me, with the newly collected samples matching the var. *orthotriaena/microana* samples. These differences are that the var. *megana* sample has longer oxeas I, longer oxeas II, longer clads on the plagiotriaenes, and larger sterrasters. It therefore seems possible that this variant is a different species than the other samples. On the other hand, I was able to sequence 140 bp of the cox1 locus from one var. *megana* sample, and it was identical to a var. *microana* sample and two of the recently collected samples. Such a small region of a single gene is hardly conclusive, but without more data and additional samples, I agree with Lendenfeld that evidence for an additional species is not yet sufficient.

de Laubenfels (1932) considered *G. bicolor* (Lendenfeld 1910) a synonym of *G. angulata*, despite the fact that he did not examine any material from either species. His justification was that the only declared differences between them were in external color and the presence of bent megascleres in *G. angulata*, but this assertion is inaccurate. The names of these species are based on those characters, but Lendenfeld also indicated that *G. angulata* has smaller sterrasters than *G. bicolor*, multiple differences in aster morphology including the lack of spines on oxyasters in *G. angulata*, and anatriaenes that were only present in *G. angulata*. Anatriaenes are rare and may be absent in some *G. angulata*, so this character is difficult to use in diagnosis. The bent oxeas are also a difficult feature, as the bend is subtle and present on only some oxeas, but it does seem to occur in at least a few oxeas of all *G. angulata* samples and was absent in the two *G. bicolor* samples I reexamined. The sterrasters, on the other hand, are 60% larger in *G. bicolor* (t-test p-value < 2.2e-16); they are also more oval-shaped in *G. bicolor* (length/width = 1.1 in all *G. angulata* samples examined, but 1.4 in both *G. bicolor* samples examined). SEM data from one sample of each species was consistent with Lendenfeld’s assertion that the oxyasters differ as well, with spines present only in *G. bicolor,* and a higher proportion of the *G. angulata* oxyasters having a prominent centrum. I agree with Lendenfeld that the differences in sterrasters and oxyasters are sufficient to consider both species valid, pending the analysis of further samples. DNA sequencing cannot yet come to bear on the question, as I was unable to amplify DNA from historical *G. bicolor* samples, and did not locate any new samples.

It is interesting that the newly collected sample from San Diego did not have protruding spicules in situ, while the two newly collected samples from Carmel were covered in a very long and thick spicule fur. Variation in the presence of protruding spicules is frequently noted in tetractinellids, but often assumed to be caused by rough handling during the collection process, or "due to such external causes as, for example, overlying bryozoans" (de Laubenfels 1932). The San Diego sample may have been scoured in the past, but was not buried or overgrown when found, so it seems worth considering that the spicule fur could be a plastic response to environmental conditions, or a trait that is periodically shed and regrown, perhaps as a response to predation or fouling.

This species is one of the few California astrophorids that previously had some sequence data available: a partial cox1 sequence (EU442203, figure S1). This was generated as part of a previous phylogenetic analysis of the Astrophorina, where it was found to be more closely related to species of *Rhabdastrella* than to other *Geodia*, albeit with low statistical support (Cárdenas *et al*. 2011). Here, I have generated full-length ribosomal data with Illumina sequencing, and confirm this placement. In the ribosomal tree, the species is most closely related to *Rhabdastrella globostellata* (Carter 1883), with 98% bootstrap support (figure S2). Placement in the cox1 tree is similar, but less well supported (figure S1).

### *Geodia bicolor* (Lendenfeld 1910)

Figure 20

**Figure 20.**
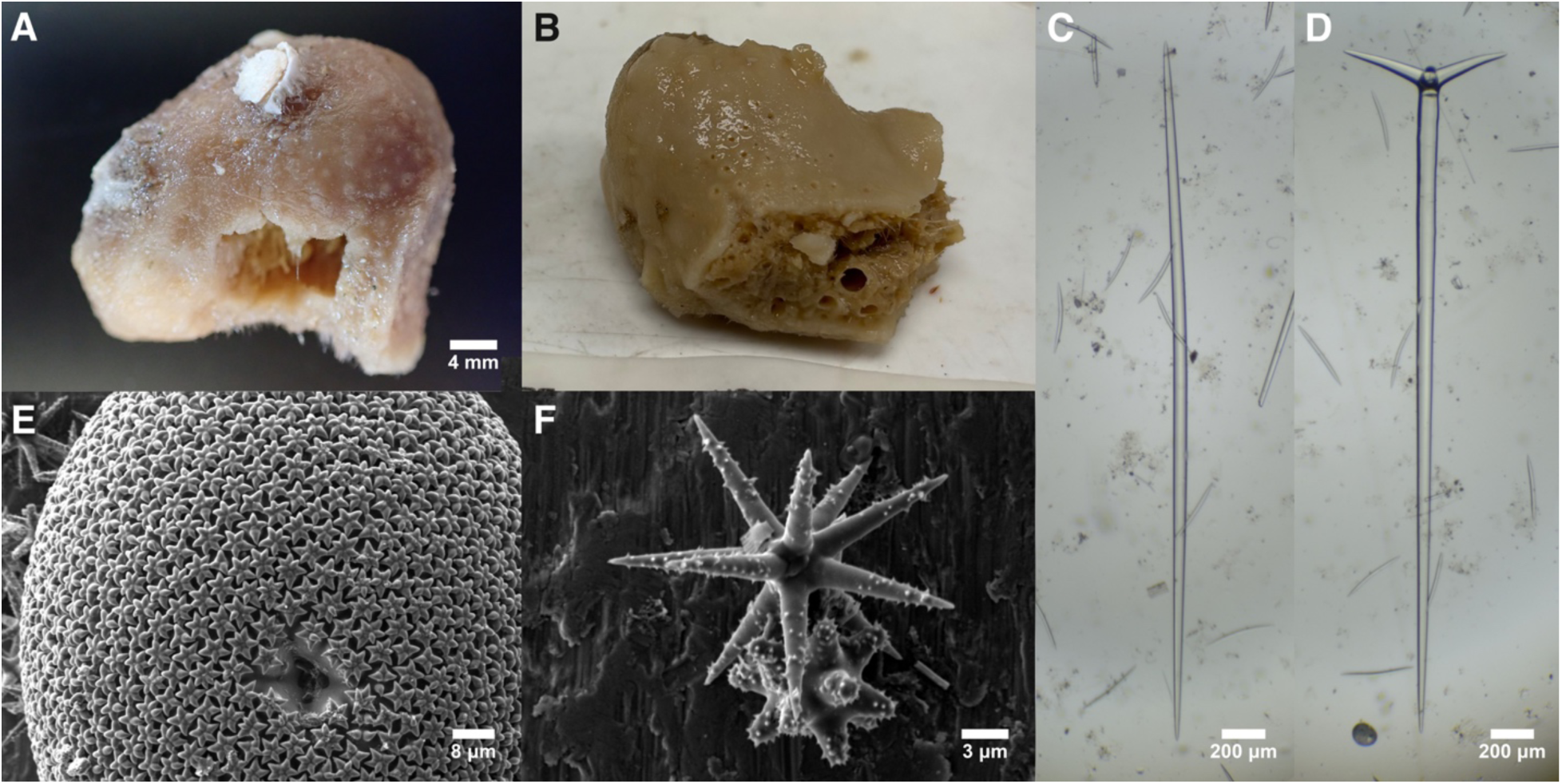
Geodia bicolor. A: Loaned portion of syntype USNM 8399. B: Entirety of syntype USNM 8398; photo by Abigail Reft, Smithsonian Museum. C: Oxea I from USNM 8398. D: Plagiotriaene from USNM 8398. E: Sterraster from USNM 8399. F: Acanthose oxyaster (top) and acanthose strongylospheraster (bottom).

#### Material examined

Syntypes: USNM 8398, San Nicolas Isl., California, (33.30, -119.40), 82 m, 13-Feb-1889; USNM 8399, E of San Nicolas Isl., California, 59-60 m, 12-Apr-1904.

#### Morphology

Only small fragments of two type specimens were reexamined, but the morphology is described in detail by Lendenfeld (1910). His account notes that sponges are thickly encrusting, up to 101 mm in horizontal extent and 40 mm thick. They are white, red, or purple-brown post-preservation, and darker on the upper surface. Spicule fur is present, but less extensive in areas with oscula; these appear as many small pores, as in *G. angulata* described above.

#### Skeleton

Not investigated here, but described in detail by Lendenfeld (1910). Briefly, the choanosome contains bundles of oxeas I (and possibly oxeas II) extending to the surface and widening towards the periphery, where they pierce the surface and create a spicule fur of oxeas. Near the surface, plagiotriaenes are added to bundles. The cladomes of the plagiotriaenes occur mainly below or within the cortical layer at the surface, but some extend into the spicule fur. The cortical layer contains the sterrasters. Oxyasters are reported to be limited to the choanosome, with oxyspherasters primarily in cortical canals and subcortical cavities. Strongylospherasters populate the outer layer of the cortex.

#### Spicules

Shown in figure 20 except where noted.

Oxeas I: stout, somewhat fusiform, and tapering to points at both ends; very rarely seen as styles. Variable in size across samples. Newly isolated spicules are 1790–3037–4726 × 31–54–66 μm (n=14). Reported as 2300–5600 × 35–105 μm in the type description, with maximum lengths per sample varying from 3700–5600 μm (Lendenfeld 1910).

Oxeas II: differentiated from oxeas I by being thinner relative to their length and less fusiform. Few were seen in the small subsamples used to isolate new spicules here, likely due to having few protruding spicules. Newly isolated spicules are 4511–4799–5283 × 22–29–33 μm (n=3). Reported as 3500–9000 × 15–40 μm in the type description (Lendenfeld 1910). Not shown in figure 20; see figure 19D for similar example.

Short oxeas: short oxeas (300–400 μm) are abundant in syntype USNM 8398 (visible in figure 20 C and D), but not in the other syntype; they were not mentioned by Lendenfeld (1910) and are presumed to be foreign.

Plagiotriaenes: with long rhabds usually tapering to sharp points, but a few are short and strongylote. The clads are frequently unequal in length, and only the largest clad was measured. Rhabds 2022–2787–3383 μm (n=13) × 52–80–96 μm (n=20). Clads 340–435–537 (n=18) × 38–61–76 (n=20) μm. Reported with rhabds 2100–4000 × 62–110 μm and clads 280–700 μm in the type description (Lendenfeld 1910). The rhabd:clad angles were 104°–110°–113° (n=9), similar to the 103°–122° reported by Lendenfeld.

Sterrasters: Oval when seen from above, with length:width ratios of 1.4 (Lendenfeld examined a larger number of samples, and found this to vary from 1.2–1.4). Rosettes are not warty. Mature sterrasters (those with rays ending in rosettes) are 108–146–173 (n=164) × 79–111–136 (n=98) × 79–95–125 (n=27) μm. Reported as 130–170 × 100–133 × 77–97 μm in the type description (Lendenfeld 1910).

Oxyasters/oxyspherasters: Acanthose asters with sharply pointed rays. Lendenfeld divides them into oxyasters and oxyspherasters, while noting that they form a continuous spectrum. He also noted that the oxyspherasters have a less pronounced centrum than seen in *G. angulata*. Newly measured spicules were considered one class, 18–22–29 μm (n=56). Reported in the type description as 19–34 μm for oxyasters and 14–23 μm for oxyspherasters (Lendenfeld 1910).

Strongylospherasters: with a large, smooth centrum and short, heavily spined, strongylote rays. In the syntype examined with SEM, total diameters were 8–11–14 μm (n=26). Reported as 9–22 μm in the type description, which analyzed more samples and includes additional detail about shape and size (Lendenfeld 1910).

#### Distribution and habitat

Known from Central and Southern California, from 48–135 m depth. The 15 syntypes were dredged from near San Nicolas and San Miguel Islands in Southern California and Carmel Bay in Central California. A sample collected near Cordell Bank in 1949 (not examined here) is the only other sample assigned to this name, to my knowledge.

#### Remarks

I isolated and measured new spicules from two of Lendenfeld’s 15 syntypes. The new data are consistent with the data from Lendenfeld, but useful because Lendenfeld did not report mean values. Having complete distributions also allowed me to verify that the sterrasters of this species are significantly larger than those from *G. angulata* (t-test p-value < 2.2e-16). This trait, together with the acanthose oxyasters (unspined in *G. angulata*), allows us to differentiate these two similar species. In contrast to *G. angulata*, I was unable to obtain any DNA data from the syntypes of this species to verify species differentiation.

I was unable to locate any new samples of this species at diving depths, despite searching over 100 locations in Central and Southern California, including both islands where Lendenfeld’s samples were dredged. This may indicate that depth partially differentiates these two species, as *G. angulata* has been found from 9–66 m, and *G. bicolor* from 48–135 m

### *Geodia carolae* (Lendenfeld, 1910)

Figure 21

**Figure 21.**
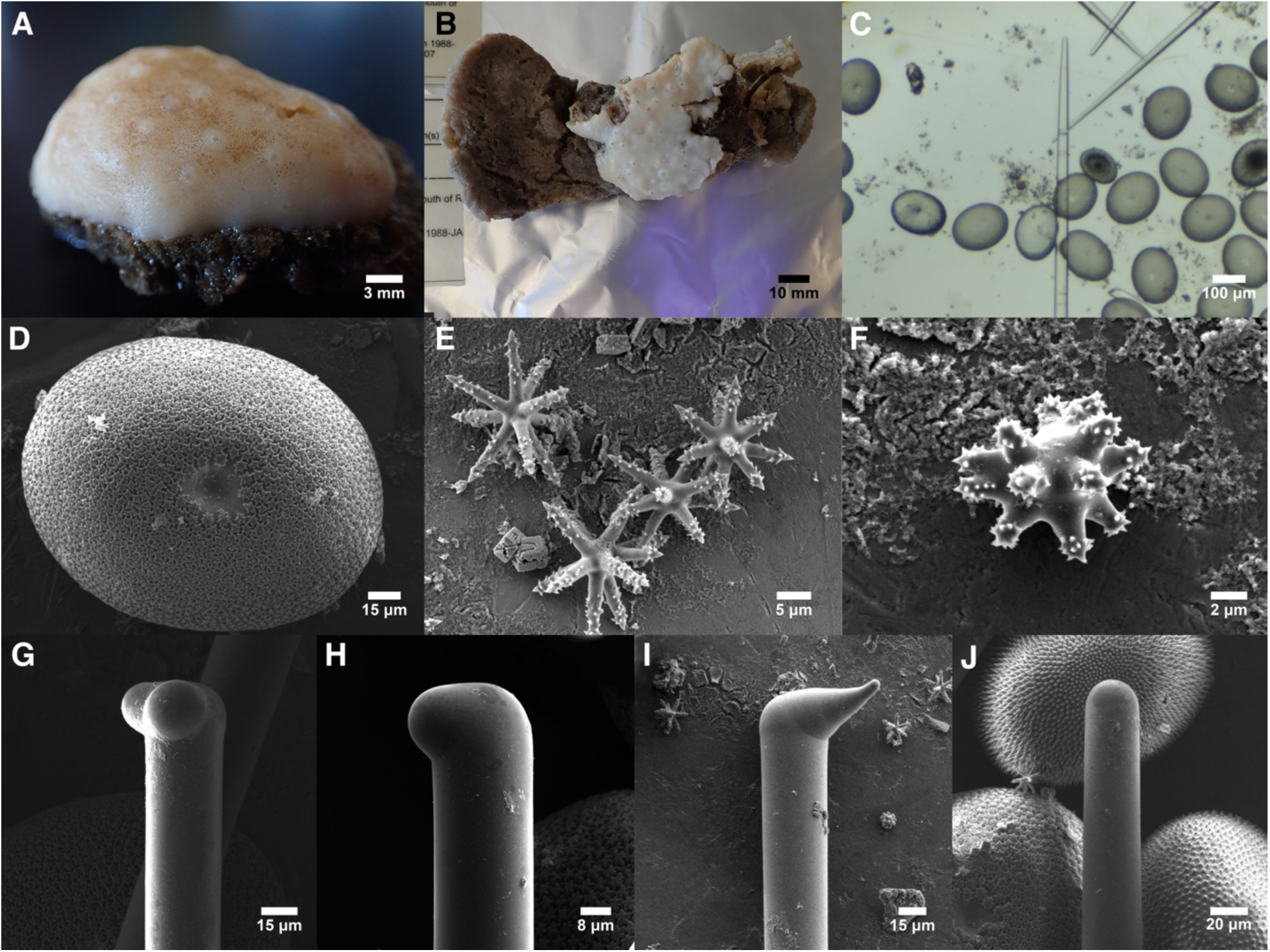
Geodia carolae. A: Preserved sample RBCM 980-00232-008 on glass sponge skeleton. B: Preserved sample RBCM 988-00002-094 (white) encrusting on Poecillastra rickettsi sample RBCM 988-00002-095 (brown). C: Tip of strongyloxea with sterrasters. D: Sterraster. E: Oxyasters. F: Small spheraster. G: Diaene. H-I: Monaenes. J: Style with sterrasters. Immature sterrasters (those without rosettes, which appear with dark centers in C, and with protruding rays in J top and lower right) were not included in the size measurements for this spicule type. C-J are from the syntype, USNM 8391.

#### Synonyms

*Geodinella robusta* Lendenfeld, 1910

*Geodia lendenfeldi* Stone *et al*. 2011

#### Material examined

Syntype: USNM 8391, Revillagigedo Isl., Behm Canal, Naha Bay, Indian Point, Alaska, 75-245 m, 7-Jul-1903. Other samples: RBCM 980-00257-005, Greenway Sound, off Sutlej Channel, British Columbia, (50.86167, -126.72333), 150-220 m, 23-Mar-1980; RBCM 980-00232-008, Pt. Upwood, Texada Isl., British Columbia, (49.48833, -124.12833), 20-60 m, 18-Mar-1980; RBCM 988-00002-094, Fitz Hugh Sound; Rivers Inlet, British Columbia, (51.45000, -127.81667), 125 m, 19-Jan-1988.

#### Morphology

Some samples are encrusting to cushion-shaped, up to 45 × 35 × 15 mm, with many small (∼400 μm) scattered oscula. Others are upright and digitate, with an apical opening up to 1.5 mm wide. Post-preservation, white externally and beige internally. The external surface is a very hard cortex, but the internal choanosome is soft and yielding. For further information, see Lendenfeld (1910) and Stone *et al*. (2011).

#### Skeleton

Not investigated here, but described by Lendenfeld (1910). The choanosomal skeleton seems somewhat vestigial compared to the other *Geodia* in the region. It is composed of both types of megascleres (oxeas/styles/strongyloxeas/strongyles and plagiomonaenes/dienes). In upright forms, these form a loose longitudinal column, with oblique, plumose bundles projecting towards the surface. Sterrasters densely populate the central cortex, with some in the choanosome. Oxyasters are found mainly in the choanosome, with strongylospherasters mainly in the outer cortex.

#### Spicules

Shown in figure 21 except where noted.

Oxeas/styles/strongyloxeas/strongyles: Varying from oxeas with two pointed tips, to strongyloxeas which taper to one pointed and one rounded tip, to strongyles with two rounded ends, to styles with one pointed tip and the other end thick and rounded and sometimes even subtylote. Dimensions of all shapes are similar and are pooled here. 1025–1935–2887 × 12–34–51 μm (n=58). Reported as 370–2800 × 30–80 μm in the type description (Lendenfeld 1910). Oxea not shown; see figure 20C for similar example.

Plagiodiaenes/plagiomonaenes: with long rhabds usually tapering to sharp points, but occasionally with strongylote rhabds. A few have one or two tapering clads, and are clearly derived from plagiotriaenes, but most have only one or two rounded protuberances at the cladomal end of the spicule. They are similar in length to the other class of megascleres, and some styles included above may be derived from triaenes that have completely lost all clads. Rhabds 700–1563–2070 μm (n=36) × 11–30–43 μm (n=40), clads 10–44–135 (n=36). Reported with rhabds 1100–2100 × 26–42 μm and clads 30–105 μm in the type description (Lendenfeld 1910).

Sterrasters: Oval when seen from above, with length:width ratios of 1.25; a minority are somewhat triangular or otherwise slightly irregular in shape. Rosettes are not warty, with 3–9 spines per rosette. Mature sterrasters (those with rays ending in rosettes) are 116–169–200 (n=225) × 101–135–158 (n=56) × 66–99–120 (n=23) μm. In the type description, these were reported as 180–195 × 130–160 × 80–115 μm for var. *carolae,* and the sterrasters were said to be uniformly elliptical. Shapes were mostly elliptical but included some irregular forms in the other two varieties, with var. *megaclada* being 190–217 μm in maximum diameter, and var. *megasterra* 220–237 μm in maximum diameter (Lendenfeld 1910).

Oxyasters: Asters with sharply pointed acanthose rays. In the syntype examined with SEM, spicules are 10–19–26 μm (n=61) in total diameter. Reported in the type description as 9–38 μm (Lendenfeld 1910).

Small spherasters: Smaller asters with a large centrum and acanthose rays. Called strongylospherasters by Lendenfeld, but some appear to have oxyote rays. Newly measured spicules are 7–11–15 μm (n=40) in total diameter. Reported in the type description as 7–13 μm (Lendenfeld 1910).

#### Distribution and habitat

Excepting one sample, this species is known from the Northern Sea of Japan (Koltun 1966), throughout the Aleutians Islands, Southeast Alaska, and British Columbia, with the shallowest sample collected between 20–60 m and the deepest between 75–245 m (samples are confirmed to at least 190 m). One outlying sample was collected from 274 m near Santa Cruz Island, in Southern California (see remarks). Samples were collected from cobble, pebble, and sand bottoms, with several epizoic on other tetractinellid sponges (one on *Erylus aleuticus*, another on *Poecillastra rickettsi*) and one was encrusting the skeleton of a glass sponge.

#### Remarks

This species was originally described as a member of *Geodinella* Lendenfeld 1903, but this genus was later synonymized with *Sidonops* Sollas, 1889 (Uriz 2002b), and *Sidonops* was then synonymized with *Geodia* (Cárdenas *et al*. 2010). These changes made *G. robusta* (Lendenfeld, 1910) a junior homonym of *G. robusta* Lendenfeld, 1907, and the name *G. lendenfeldi* was proposed as a resolution (Stone *et al*., 2011). However, because there were named varieties, the rules of zoological nomenclature dictate that the new name should be taken from the varieties, and the species is now known as *G. carolae* (van Soest *et al*. 2020), based on the previous name *Geodinella robusta* var. *carolae*, which Lendenfeld says was named after Queen Charlotte Sound, British Columbia.

Lendenfeld allocated four samples to this species, but divided them into three varieties. I reexamined only one type sample, one of two syntypes for var. *carolae*, and my spicule measurements agreed well with Lendenfeld’s data. I also examined three more recently collected samples from British Columbia; these mixed characteristics of Lendenfeld’s var. *megaclada* (larger maximum plagiomonaene clad lengths, some irregularly shaped sterrasters) and characteristics of Lendenfeld’s var. *carolae* (smaller maximum sterraster lengths, some plagiodiaenes present). There is therefore no strong evidence supporting these varieties as valid taxonomic units. The final variety, var. *megasterra*, was not examined here, but it seems possible it constitutes a different species. This variety is known from a single Southern California sample and is notable because its distribution of sterraster diameter does not overlap those of other known samples. Samples from the Aleutians are described as having sterrasters 170–200 μm (Stone *et al*. 2011); samples from British Columbia, measured here, had sterrasters 116–169–200 μm, and Lendenfeld’s samples from British Columbia and Southeast Alaska had sterrasters 180–217 μm. In contrast, the Southern California sample had sterrasters 220–237 μm. Pending genetic data or additional samples, this sample remains within the concept of *G. carolae*, but this small difference appears worthy of further attention when combined with the apparently disjunct distribution.

Unfortunately, I was unable to amplify DNA from the syntype. I was able to amplify very short (135 bp) regions of the cox1 locus from two British Columbian samples, which allowed me to place this species in the cox1 phylogeny (figure S1). The placement is surprising, as these sequences fall within a clade otherwise populated with *Penares* and *Erylus*. It is important to note that this placement is highly preliminary due to the modest bootstrap values, the short length of the sequences, and the lack of any closely related taxa with longer sequences. This species is not present in figure 4 because no data was successfully generated at the 28S locus.

### Geodia agassizii Lendenfeld, 1910

Figure 22

**Figure 22.**
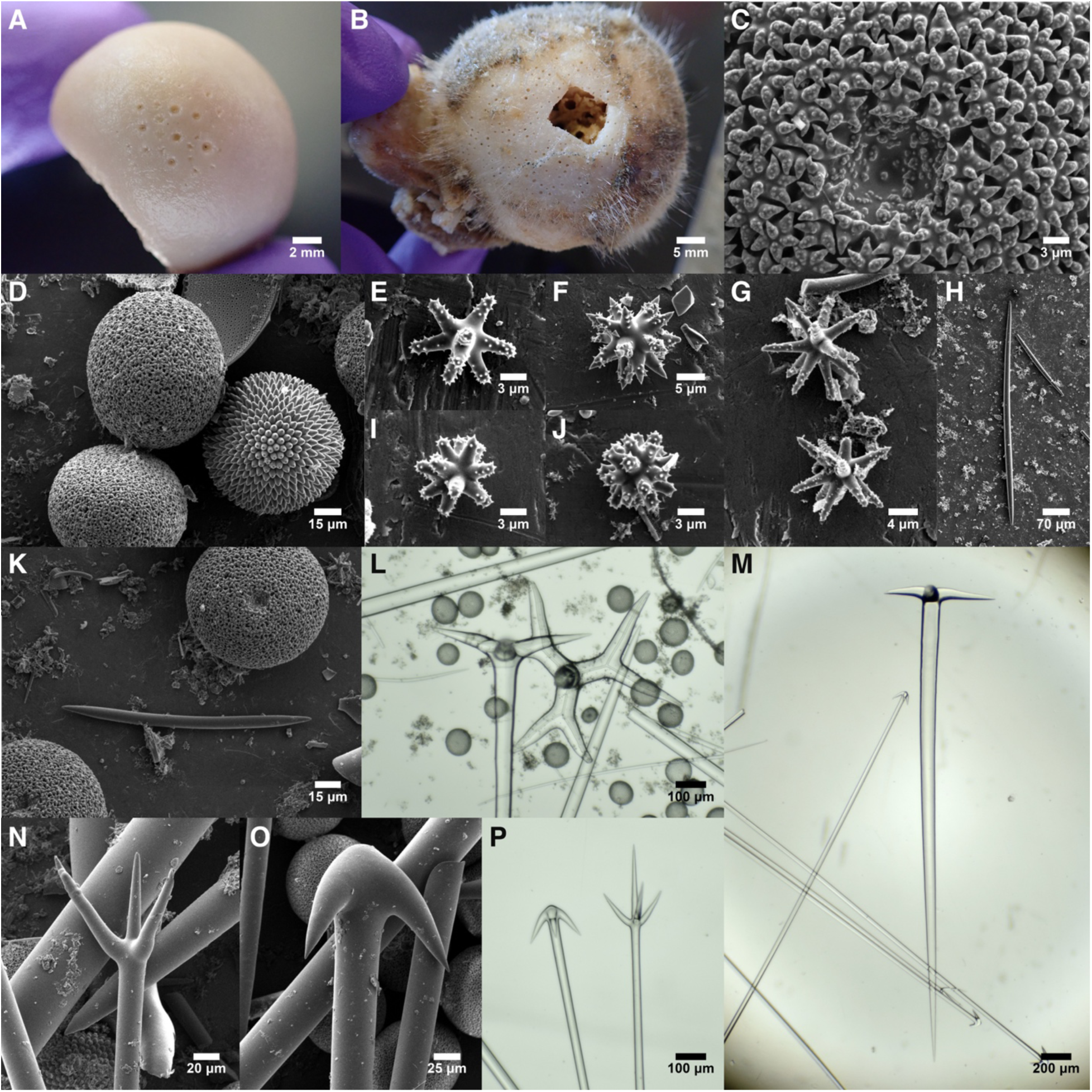
Geodia agassizii. A: Preserved syntype USNM 8381. B: Preserved sample RBCM 019-00567-003; cut-out was made by a previous taxonomist. C: Sterraster detail. D: Sterrasters including one that is immature (right). E, I, J: Small strongylasters. F, G: Large oxyasters. H: Oxea I. K: Short oxea with sterrasters. L: Plagiotriaene with dichotriaene cladome that has broken from its rhabd. M: Plagiotriaene with anatriaenes and a mesoprotriaene from RBCM 019-00567-003. N: Protriaene. O: Anatriaene. P: Anatriaene (left) with mesoprotriaene (right) from RBCM 019-00567-003. All spicules from syntype USNM 8381 except where indicated.

#### Synonyms

*Geodia japonica* sensu Lehnert & Stone (2016)

#### Material examined

Syntypes: USNM 8362, Strait of Georgia, Bowen Island, British Columbia, 33-40 m, 20-Jun-1903; USNM 8381, Fort Rupert, Vancouver Island, British Columbia, 124-196 m, 25-Jun-1903; USNM 8384, Dunes City, Oregon, (43.98330, -124.94200), 91 m, 19-Oct-1888.

Other samples: RBCM 019-00567-003, Malcolm Island, British Columbia, (50.64417, - 126.99806), depth unknown, 28-Sep-2007; RBCM 007-00011-004, West of Nootka Sound, British Columbia, (49.81667, -127.64333), 164.6 m, 6-Oct-1999; CASIZ 229525, Nehalem Bank vicinity, Oregon, (45.93333, -124.60000), 176 m, 14-Oct-1969; USNM 1478670, Shutter Ridge, West of Cape Ommaney, Alaska, (56.19660, -135.10200), depth 82-92 m, 8-Jun-2015; CASIZ 233435, Offshore of Newport, Oregon, (44.64333, -124.43667), 100 m, 16-Apr-1962.

#### Morphology

Most are roughly spherical, but some are massive and more irregular; the largest known sample was reported by Lendenfeld (1910) as 130 × 105 × 100 mm. He also reports that most were attached by a small base, but others were cushion-shaped with a larger base. Many are smooth, without projecting spicules, or with projecting spicules only in sheltered cavities, but some are covered in projecting spicules, with fewer projecting in the oscular pore field. One or two oscular pore fields are usually evident, each with many small (< 1 mm) pores. Samples are mostly white or tan, but some are dark and bluish (e.g. syntype USNM 8384), perhaps because they have been stained with foreign pigments. Additional details available in Lendenfeld (1910).

#### Skeleton

Not investigated here, but described in detail by Lendenfeld (1910). Briefly, the choanosome contains loose radial bundles of oxeas I, widening as they extend to the surface, with some bundles piercing the surface. Mesoprotriaenes and anatriaenes are also radially arranged, with their cladomes near the surface or protruding and forming the spicule fur.

Plagiotriaenes are arranged with the rhabds directed inwards; the cladomes occur primarily just below the cortical layer. Sterrasters are found mainly in the central cortex, but some are scattered in the choanosome. Oxyasters are reported to be limited to the choanosome, with oxyspherasters primarily in the inner cortex and strongylospherasters in the outer cortex. The short oxeas are found in upright "tuft-like groups" in the dermal layer, with their ends protruding slightly beyond the sponge surface.

#### Spicules

Shown in figure 22 except where noted. See table 2 for comparisons in average values among species.

**Table 2.** Key features distinguishing species of *Geodia* with mesotriaenes. Range of means across samples is shown, in microns.

|  | Plagiotriaene<br>rhabd length | Sterraster<br>length | Mesotriaene<br>epirhabd<br>length | Anatriaene<br>clad length | Oxea I length |
| --- | --- | --- | --- | --- | --- |
| *G. mesotriaena* | 4950–6600 | 94–106 | 143–200 | 153–183 | 4900–5800 |
| *G. agassizii* | 2100–3350 | 85–96 | 141–193 | 101–150 | 2450–3950 |
| *G. ovis* | 5100 | 82 | 80 | 124 | 5600 |
| *G. mesotriaenella* | 1650–1900 | 81–82 | 92–105 | 106 | 1700–1850 |

Oxeas I: stout; tapering to points at both ends, but occasional styles or strongyles also occur. Highly variable in size, with mean lengths per sponge 2451–3902 μm. All spicules pooled, 1658–3259–5684 × 24–69–118 μm (n=159). Reported as 2300–4800 × 60–112 μm in the type description, with maximum lengths per sample varying from 3400–4800 μm (Lendenfeld 1910).

Oxeas II: separated from oxeas I by being thinner relative to their length. Only found in low numbers, and only in half the samples analyzed, but this includes two syntypes. They were not mentioned by Lendenfeld and may be foreign. 769–2523–4466 × 6–14–37 μm (n=13). Not shown in figure 22; see figure 19D for similar example.

Short oxeas: straight or slightly curved. 122–324–791 × 6–10–25 μm (n=106). Reported as 160–480 × 5–12 μm in the type description (Lendenfeld 1910).

Plagiotriaenes/dichotriaenes: with long rhabds usually tapering to sharp points, but strongylote or tylote rhabds are a considerable minority in some samples. The clads are frequently unequal in length, and only the largest clad was measured. Forked or bent clads are common in many samples, and some have four clads instead of three. In some cases, all clads are forked and these samples therefore include dichotriaenes, which are pooled here. Rhabds 1105–2888–4313 μm (n=131) × 34–101–144 μm (n=144). Clads 148–413–630 (n=127) × 39–79–111 (n=127) μm.

Reported with rhabds 1500–4200 × 65–150 μm and clads 240–560 μm in the type description (Lendenfeld 1910). The rhabd:clad angles were 74°–102°–123° (n=103), similar to the 73°–117° previously reported; these triaenes were referred to as "orthoplagiotriaenes" in the type description (Lendenfeld 1910).

Mesoprotriaenes/protriaenes/mesoprodiaenes: Mesoprotriaenes with three curved clads and a straight central epirhabd are present in all samples. Many samples also have protriaenes (lacking the epirhabd) or mesoprodiaenes/mesopromonaenes (appearing to have the epirhabd but lacking one or more clads; these are sometimes hard to differentiate from protriaenes). In mesoprotriaenes, the epirhabd is usually longer or equal to the clads in length, but it is shorter in some spicules, and the epirhabd:clad length ratio is reported as highly variable across samples by Lendenfeld (1910). When all spicules are pooled, rhabds are 2576–5244–7106 μm (n=18) × 12–26–47 μm (n=96); epirhabds are 68–169–275 μm (n=53), and the longest clads are 66–169–380 μm (n=90) × 9–21–33 μm (n=88) μm. For mesoprotriaenes alone, the epirhadb-to-longest-clad ratios were 0.71–1.10–1.79 (n=49). Lendenfeld reported rhabds 2000–6000 × 7–40 μm, epirhabds 25–320 μm, and clads 60–250 μm in the type description (Lendenfeld 1910). The rhabd:clad angles of newly isolated spicules were 28°–40°–51° (n=47), nearly identical to the 22°–38°–55° previously reported (Lendenfeld 1910).

Anatriaenes/anadiaenes: Seen in all specimens. Rhabds are long and usually taper to a filament-like ending. Clad size varies greatly within a sample, but they do not seem to form size classes. Forked clads are sometimes seen, as are anadiaenes, sometimes with both remaining clads close together. When all spicules are pooled, rhabds are 5353–8167–11355 μm (n=10) × 10–30–44 μm (n=112); clads are 56–124–175 (n=112) × 16–34–58 (n=102) μm, and rhabd:clad angles are 31°–39°–48° (n=70). Lendenfeld (1910) reported rhabds 4000–9000 × 10–50 μm, clads 40–155 μm, and rhabd:clad angles of 32°–46°–65°. He also reported variation among samples, with a small sample having maximum clad lengths of only 100 μm.

Small anatriaenes: Much smaller than the primary anatriaenes, and possibly immature. Not seen in any new spicule preparations, but reported as present in the smallest examined sample of this species by Lendenfeld (1910). He reported rhabds 290 × 1.0–1.5 μm, clads 4–6 μm, and rhabd:clad angles of 38°–62°. Not shown.

Sterrasters: Nearly round when seen from above, with length:width ratios of 1.1. Rosettes are warty, with 3–10 spines per rosette. When averaging 10 or more rosettes per sterraster the number of spines per rosette is 6–7–9 (n=4 sterrasters). Mature sterrasters (those with rays ending in rosettes) are 57–91–109 (n=508) × 64–80–93 (n=127) × 52–62–76 (n=52) μm.

Reported as 82–118 × 75–100 × 58–83 μm in the type description; see Lendenfeld (1910) for additional details, including discussion of rare forms he dubs "sterroids".

Large sterrasters: A very small minority of sterrasters in syntype USNM 8381 were much larger than the others. One large sterraster was also seen in one sponge from voucher RBCM 019-00567-003. None were seen in the other samples, or reported by Lendenfeld (1910), so these may be rare aberrations or foreign spicules, but care should be taken to avoid misidentifications relative to *G. starkii*, which has two size classes of sterrasters. 145–173–184 × 123–136–147 μm (n=6).

Oxyasters/oxyspherasters: Acanthose asters with sharply pointed rays. While he noted that they form a continuous spectrum, Lendenfeld still divided them into oxyasters and oxyspherasters. I found this division to be difficult, and all are pooled here. Newly measured spicules are 8–14–28 μm (n=75) in total diameter. Reported in the type description as 9–31 μm for oxyasters and 10–21 μm for oxyspherasters; see Lendenfeld (1910) for additional details.

Strongylasters/strongylospherasters: Acanthose asters with blunt, strongylote rays. These were difficult to separate from oxyasters. Newly measured spicules are 6–9–15 μm (n=66) in total diameter. Reported as 3.5–11 μm in the type description, which analyzed more samples and includes additional detail about shape and size (Lendenfeld 1910).

Oxyasters/strongylasters combined: Defining classes of asters based on shape was difficult, so I also present their combined distribution here. Together, total diameters are 6–12–28 μm (n=141). The distribution is continuous but with major modes at 7 and 11 μm, and a smaller mode at 16 μm.

#### Distribution and habitat

Known from the Gulf of Alaska to Anacapa Island, Southern California. The shallowest sample was collected between 33–40 m depth, and the deepest was between 75–245 m. Some were found growing on the skeletons of dead glass sponges, including multiple instances where sponges were found inside tubular hexactinellid skeletons. One was found growing on *Penares orientalis* comb. nov.

#### Remarks

*Geodia agassizii* was originally described based on 22 samples, considerably more than the other *Geodia* from the region, and these spanned a broad latitudinal range, from Southeast Alaska to Southern California (Lendenfeld 1910). Despite this relative abundance, no sponges have been assigned to the species since, based on records in GBIF. This is likely due to uncertainty regarding the number of *Geodia* species present in the region and how to differentiate them. In his monographic work on the sponges of California, de Laubenfels (1932) declared that all of Lendenfeld’s *Geodia* species that possessed mesotriaenes should be lumped under the name *G. mesotriaena*. This reduced *G. agassizii*, *G. ovis*, *G. mesotriaenella,* and *G. breviana* to synonyms. As of this writing, these names are once again considered valid in the World Porifera Database (de Voogd, *et al*. 2025), but no further work has been published on any of these species, as far as I am aware.

I was able to sequence short (∼140 bp) fragments of the *cox1* locus from two *G. agassizii* syntypes and three *G. mesotriaena* syntypes. These sequences are quite different (divergence (Dxy) = 8.7%; Fst = 1.0) and verify that the species are distinct. DNA sequencing failed from the type material of *G. ovis*, *G. mesotriaenella*, and *G. breviana*, but I was able to differentiate *G. agassizii* from all of these species morphologically. Of the many spicule measurements taken, I focused on five that were easy to reproduce, visible under light microscopy, and appeared to vary among species: plagiotriaene rhabd length, sterraster length (maximum diameter), mesotriaene epirhabd length, anatriaene clad length, and oxea I length. When the mean values of these traits are compared in a principal components analysis, *G. agassizii* is differentiated from all other sympatric *Geodia* (figure 23). The plagiotriaene and sterraster lengths appeared to be the most informative traits, and a two-dimensional plot of these features is sufficient to differentiate *G. agassizii* from sympatric species (figure S3). The sterrasters of *G. agassizii* are larger than *G. ovis* and *G. mesotriaenella* (*G. breviana* is synonymized with *G. mesotriaenella* below), and the plagiotriaenes are shorter than *G. mesotriaena* and *G. ovis.* As shown in table 2, differences in additional spicule types add confidence to morphological species discrimination in this group. Lendenfeld (1910) also states that *G. mesotriaena* has preoscular cavities but *G. agassizii* does not (this was not investigated here). He additionally found the choanosomal oxyasters to be smaller in *G. agassizii* and the strongylospherasters to have a different shape, when this species is compared to *G. mesotriaena*. I measured oxyasters from two samples of each species using SEM, and though the overall trend in size agrees with Lendenfeld’s observation, the *G. agassizii* with the largest oxyasters appeared to be indistinguishable from the *G. mesotriaena* with the smallest oxyasters. I was also unable to differentiate samples based on the shapes of the strongylospherasters. Gross morphology may also be useful, if not diagnostic, in differentiating these species, as *G. agassizii* is nearly spherical while *G. mesotriaena* and *G. ovis* are cake-shaped (though some *G. mesotriaenella* are spherical). Finally, note that I could not confirm any cases of *G. ovis* and *G. mesotriaena* North of Point Conception, California, though this may change as more samples are reexamined with the methods detailed here (see *G. mesotriaena* remarks for further information about that species’ uncertain range).

**Figure 23.**
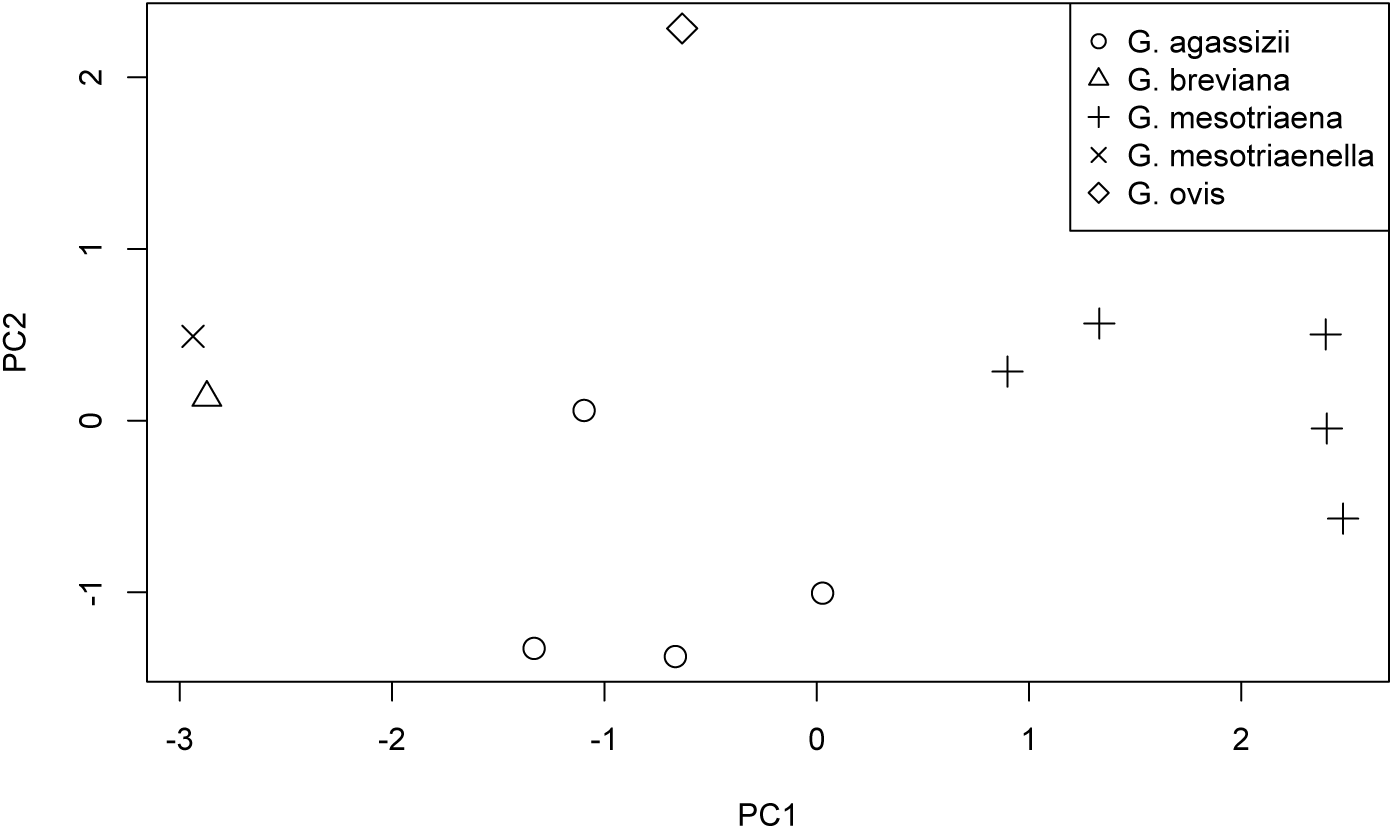
Principal Components Analysis of morphological variation in *Geodia* species with mesoprotriaenes. When the average values of five key spicule traits (shown in table 2) are plotted with a PCA, all species are distinguishable except *G. mesotriaenella* and *G. breviana*. Below, it is proposed that *G. breviana* is a junior synonym of *G. mesotriaenella*.

Though I did not locate any new samples of *G. agassizii* at diving depths, I was able to use the morphological and DNA differences described above to reassign several sponges previously described as *G. mesotriaena* or *Geodia* sp. to *G. agassizii.* I likewise found that a sponge previously described as *G. japonica* (Sollas 1888) is in fact *G. agassizii*. Seven spherical *Geodia* were dredged from a single location in Southeast Alaska in 2015, and assigned to the name *G. japonica* (Lehnert & Stone 2016). However, a published *cox1* sequence of one of these sponges (MT815825, from Cárdenas (2020)) matches the sequences of *G. agassizii* produced here. I obtained one sponge from this lot (USNM 1478670) and verified that both the *cox1* and *28S* loci match *G. agassizii* (figure 4). I also isolated and measured spicules from this sponge. There are some differences between the spicule data I obtained and the data previously published, perhaps because the previous data were from a different sponge in the lot (Lehnert & Stone 2016). Regardless, based on either dataset, the spicules are a better match to *G. agassizii* than *G. japonica*. According to Lendenfeld (1910), "*Geodia japonica* is distinguished from *G. agassizii* by being cup shaped, having rounded, lobate protuberances on the outer surface, and possessing, although very large in size, considerably smaller megascleres; the large [oxeas] of this species are less than half as thick as those of *G. agassizii*" (Lendenfeld, 1910, p. 150). He goes on to say that the "sigmoid clades" on the anatriaenes are "very characteristic" of *G. japonica*. The Alaskan samples do not have these sigmoid clades, and they are all spherical like *G. agassizii*, rather than cup-shaped with a lobed outer surface. It is true that the oxea widths published by Lehnert and Stone (2016) are thin for *G. agassizii* (32–38–62 μm), and in this sense, fit *G. japonica* better. However, when I measured the oxeas in the sponge archived as USNM 1478670, I found them to have widths of 57–72–89 μm, fitting *G. agassizii*. Also note that the thinnest *G. agassizii* described by Lendenfeld had oxeas of only 20–66 μm wide (Lendenfeld, 1910, p. 120), which match the published data for these new Alaskan samples. Other spicule data in both reports is a marginally better fit to *G. agassizii* as well. The oxyasters as measured by me (8–13–21 μm) and the previous authors (6–9–11 μm & 16–19–23 μm) are a better fit to *G. agassizii* (9–31 μm according to Lendenfeld) than *G. japonica* (17–46 μm according to Lendenfeld). The same is true of sterraster size: (mine, 82–94–104 μm; previous authors, 65–90–102 μm; *G. agassizii* according to Lendenfeld, 82–118 μm; *G. japonica* according to Lendenfeld, 80–89 μm), and strongylaster shape (having a very large centrum in *G. japonica*, but being similar to small oxyasters in *G. agassizii* (Lendenfeld 1910)). Additionally, *G. agassizii* is expected in Southeast Alaska, while *G. japonica* is described from the warm temperate waters in central Japan. Together, the DNA data, gross morphology, spicules, and biogeographic data all support reassigning these samples to *G. agassizii*. There is one other claim of *G. japonica* in the Eastern Pacific: a sponge collected in the Gulf of California was assigned to this name by Dickinson (1945). The limited morphological data included in the associated report is insufficient to identify this sample, but it seems to match *G. mesotriaena* at least as well as *G. japonica*. I therefore do not think there is any solid evidence of *G. japonica* in the Eastern Pacific.

### Geodia mesotriaena Lendenfeld, 1910

Figure 24

**Figure 24.**
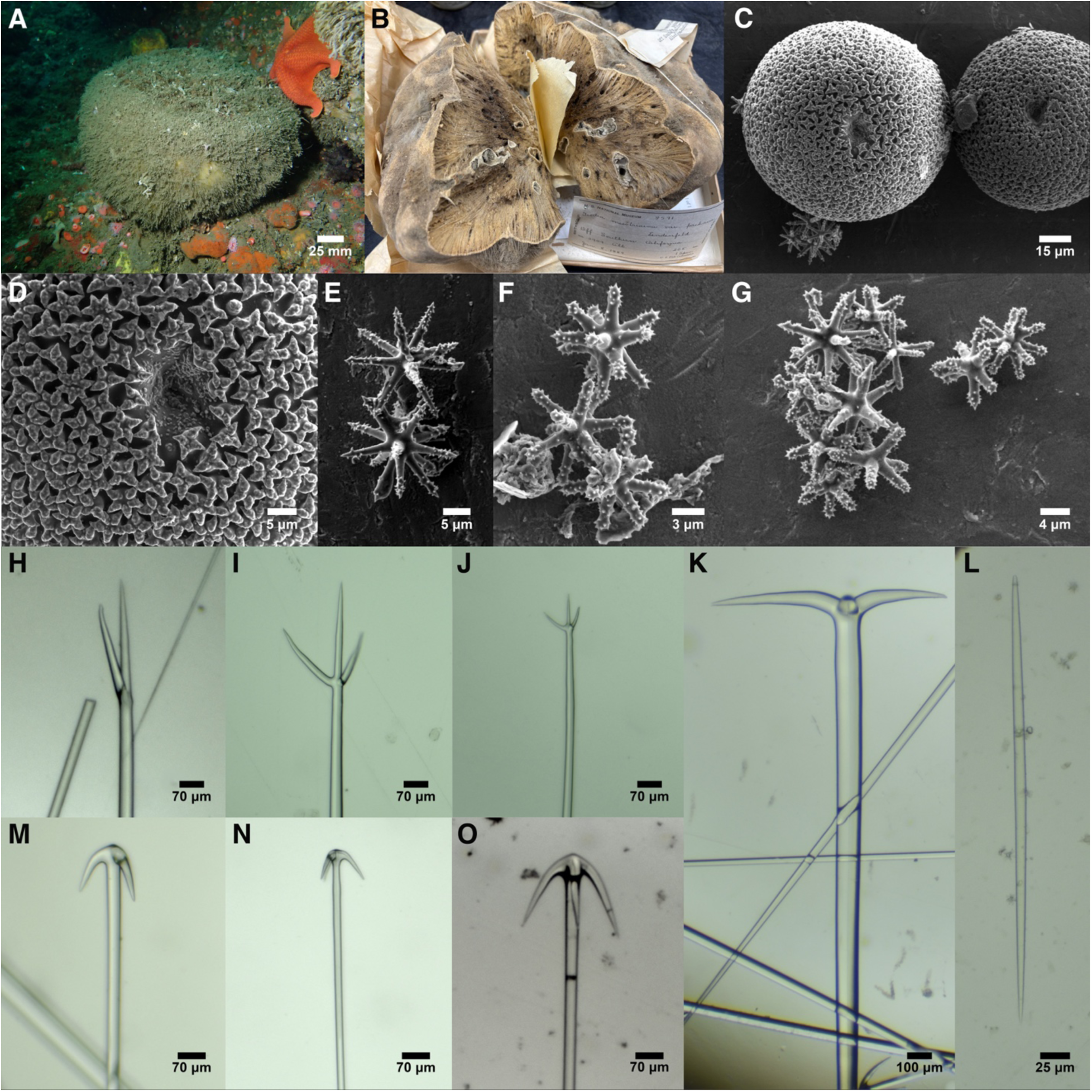
Geodia mesotriaena. A: TLT589 in situ. B: Dried var. pachana holotype, USNM 8571, photo curtesy of Abigail Reft, Smithsonian Institution. C-D: Sterrasters. E: Large oxyaster. F: Small strongylasters. G: Aster variety, including both types. H: Promesomonanine. I-J: Promesotriaenes. K: Plagiotriaene. L. Short oxea; note that it is slightly asymmetrical. C-L from TLT589. M: Anatriaene from var. pachana holotype, USNM 8571. N: Anatriaene from var. microana holotype, USNM 8410. O: Anatriaene from var. megana holotype, USNM 8409.

#### Material examined

Type material: var. *pachana* holotype USNM 8571, Santa Barbara Channel, Santa Barbara, California, (34.36670, -120.14200), 375 m, 8-Jan-1889; var. *microana* holotype USNM 8410, San Pedro Channel, Los Angeles, California, (33.64580, -118.22900), 37 m, 5-Feb-1889; var. *megana* holotype USNM 8409, NE of San Miguel Isl., California, (34.06670, - 120.32500), 47 m, 9-Feb-1889. Other samples: TLT1444, Off Torrey Pines, San Diego, California, (32.92505, -117.29567), 67 m, 13-Sep-2023; TLT586 & TLT589, Train wheels, San Diego, California, (32.65205, -117.26243), 25-29 m, 19-Sep-2020; SBMNH 693792, SE of Point Conception, Santa Barbara, California, (34.42110, -120.36330), 50-55 m, 13-Jan-1983.

#### Morphology

Large sponges, up to at least 23 cm in diameter and 12 cm high. Shaped like cakes or truckles of cheese; upper surfaces are flat and undersides taper somewhat to a broad attachment region. Covered in a thick spicule fur up to a centimeter thick. Yellowish-white alive, though sediment trapped in the spicule fur leads to many appearing brown; beige after preservation. Small oscula are grouped in one or more shallow depressions (preoscular cavities). Additional details available in Lendenfeld (1910).

#### Skeleton

Described in detail by Lendenfeld (1910), and not differing markedly from other sympatric *Geodia* with mesotriaenes. Briefly, the choanosome contains loose radial bundles of oxeas I, with some piercing the surface to contribute to spicule fur. Plagiotriaenes are added to bundles near the surface, with their cladomes located just below the sterraster layer in the cortex. Anatriaenes also occur near the surface, with their clads in the cortex or protruding beyond it.

Mesoprotriaenes pierce the surface of the sponge and are the primary component of the spicule fur. Sterrasters densely populate the central cortex, with immature sterrasters in the outer choanosome and inner cortex. Oxyasters are found throughout, with larger asters in the choanosome and smaller asters more common in the inner cortex. The outer cortex contains mainly strongylospherasters. The short oxeas form upright bouquets in the dermal layer, but are also scattered throughout the choanosome. Spicule fur includes extruded spicules, as some unbroken oxeas and mesotriaenes lie loose within it.

#### Spicules

Shown in figure 24 except where noted. See table 2 for comparisons in average values among species.

Oxeas I: stout; most taper to points at both ends, but occasional styles occur, and some are intermediate, with a slightly rounded tip at one end. Oxeas with bends are occasionally seen, as in *G. angulata*. All spicules pooled, 2472–5313–8116 × 12–60–94 μm (n=95); mean lengths per sponge 4910–5797 μm. Reported as 4300–8200 × 50–105 μm in the type description (Lendenfeld 1910). Not shown in figure 24; see figure 22H for similar example.

Short oxeas: straight or slightly curved. Most are symmetrical or very nearly so, but they tend towards anisoxeas in at least one sample. 193–419–678 × 6–13–22 μm (n=98). Reported as 380–680 × 9–19 μm for ectosomal oxeas, 170–440 × 7–17 μm for choanosomal oxeas in the type description (Lendenfeld 1910).

Plagiotriaenes: with long rhabds usually tapering to sharp points, but strongylote rhabds and/or clads are a notable minority in some samples. The clads are frequently unequal in length, and only the largest clad was measured. Rhabds 2427–5932–8325 μm (n=68) × 28–80–123 μm (n=78). Clads 144–393–774 (n=75) × 21–57–100 (n=76) μm. Reported with rhabds 4600–7200 × 75–120 μm and clads 200–670 μm in the type description (Lendenfeld 1910). The rhabd:clad angles were 92°–104°–125° (n=103), similar to the 85°–99°–117° previously reported; these triaenes were referred to as "ortho/plagiotriaenes" in the type description (Lendenfeld 1910).

Mesoprotriaenes/protriaenes/mesoprodiaenes/mesopromonaenes: Mesoprotriaenes with three curved or straight clads and a straight central epirhabd are present in all samples. Some samples also have protriaenes (lacking the epirhabd) or mesoprodiaenes/mesopromonaenes (appearing to have the epirhabd but lacking one or more clad; these can be hard to differentiate from protriaenes). In mesoprotriaenes, the epirhabd usually averages longer or equal to the clads in length, but is shorter in some spicules. Highly variable in shape, with some having more than three clads or misshapen clads. When all spicules are pooled, rhabds are 5827–9478–11188 μm (n=17) × 10–28–56 μm (n=90); epirhabds are 33–166–350 μm (n=70), and longest clads are 16–148–311 μm (n=76) × 7–18–36 μm (n=87) μm. For mesoprotriaenes alone, the epirhadb-to-longest-clad ratios were 0.50–1.13–2.12 (n=70). Lendenfeld reported rhabds 6000–14000 × 15–40 μm, epirhabds 95–330 μm, and clads 90–310 μm in the type description (Lendenfeld 1910). The rhabd:clad angles of newly isolated spicules were 17°–45°–57° (n=67), similar to the 29°–42°–54° previously reported (Lendenfeld 1910).

Anatriaenes/anadiaenes: Seen in all specimens. Rhabds are long and usually taper to a filament-like ending. Clad size varies within and among samples, but they do not seem to form size classes. Anadiaenes are sometimes seen. When all spicules are pooled, rhabds are 7153–11808–13600 μm (n=11) × 10–28–46 μm (n=187); clads are 61–169–254 (n=193) × 10–31–51 (n=193) μm, and rhabd:clad angles are 20°–36°–55° (n=70). Lendenfeld (1910) reported rhabds 11000–16000 × 22–40 μm, clads 90–270 μm, and rhabd:clad angles of 34°–58°.

Small anatriaenes: Much smaller than the primary anatriaenes, and possibly immature. Not reported by Lendenfeld (1910), and only a single case was seen in one of the newly collected samples (TLT1444). The rhabd was 7 μm wide, with the clad 17 × 6 μm. Not shown.

Sterrasters: Nearly round when seen from above, with length:width ratios of 1.1. Rosettes are warty, with 3–10 spines per rosette. When averaging 10 or more rosettes per sterraster the number of spines per rosette is 5–5–6 (n=7 sterrasters). Mature sterrasters (those with rays ending in rosettes) are 73–100–125 (n=718) × 58–88–109 (n=113) × 52–67–81 (n=76) μm.

Reported as 92–125 × 78–107 × 67–82 μm in the type description; see Lendenfeld (1910) for additional details, including discussion of rare forms he dubs "sterroids".

Oxyasters/oxyspherasters: Acanthose asters with sharply pointed rays. While he noted that they form a continuous spectrum, Lendenfeld divided them into oxyasters and oxyspherasters. I found this division to be fraught, and all are pooled here. Newly measured spicules are 11–20–38 μm (n=83) in total diameter. Two sponges were examined with SEM, and the mean values differed considerably (TLT589 = 17 μm, USNM 8409 = 23 μm, n=41 for each). Reported in the type description as 11–54 μm for oxyasters and 19–32 μm for oxyspherasters; see Lendenfeld (1910) for additional details.

Strongylasters/strongylospherasters: Acanthose asters with blunt, strongylote rays. These were difficult to separate from oxyasters. Newly measured spicules are 5–10–13 μm (n=54) in total diameter. Reported as 6–14.5 μm in the type description, which analyzed more samples and includes additional detail about shape and size (Lendenfeld 1910).

Oxyasters/strongylasters combined: Defining classes of asters based on shape was difficult, so I also present their combined distribution here. Together, total diameters are 5–16–38 μm (n=137). The distribution is continuous but with major modes at about 10 and 21 μm.

#### Distribution and habitat

Samples verified as *G. mesotriaena* in this work were spread throughout Southern California, from just Southeast of Point Conception to off Point Loma. Published morphological data are consistent with a range extending South into Pacific Mexico to at least Cabo San Lucas and the Gulf of California (Dickinson 1945). The shallowest samples verified here were collected between 25–29 m by divers, while dredged samples were up to 375 m deep. The species appears abundant at some locations off Point Loma, San Diego, below 25 meters: I searched six locations at this depth, and found numerous truckle-shaped sponges at all six (samples were taken at only one location, with the others hypothesized to be this species based on gross morphology only; note that some could prove to be *G. ovis*). I searched for sponges at 93 other locations throughout Southern California between 2019–2026, and did not find this species at any, but only three of these locations included depths below 25 m.

Museum vouchers compiled at GBIF indicate that sponges have been identified as *G. mesotriaena* across a broad latitudinal gradient, from British Columbia to the Mexican Tropical Pacific; additional records without listed vouchers span an even greater range, from the Aleutians to the Galapagos Islands. However, it is likely that some of these records are confounded with other *Geodia* species. I examined two sponges identified as *G. mesotriaena* from British Columbia, and both proved to be *G. agassizii*. Future work is needed to reexamine additional museum vouchers in light of the data published here, in order to determine the true range of this species.

#### Remarks

*Geodia mesotriaena* was described from only three samples, all from Southern California, with other mesotriaene-bearing *Geodia* from the temperate Northeastern Pacific allocated to *G. agassizii*, *G. ovis*, *G. mesotriaenella,* and *G. breviana* (Lendenfeld 1910). In his monographic work on the sponges of California, de Laubenfels (1932) thought it likely that all these names were synonyms, and because *G. mesotriaena* was listed first among these species, it became the senior synonym. Adding to the potential for confusion, an additional species from the Atlantic was later named *Sidonops meostriaena* Hentschel, 1929, and when *Sidonops* was merged with *Geodia* (Cárdenas *et al*. 2010), this became a junior homonym of *G. mesotriaena* Lendenfeld, 1910 (Cárdenas *et al*. 2010; Koltun 1966). This homonymy has now been resolved, with the Atlantic species known as *G. hentscheli* Cárdenas, Rapp, Schander & Tendal 2010.

As discussed in the *G. agassizii* remarks above, *G. agassizii* is confirmed here as a valid species, with considerable genetic differentiation between it and *G. mesotriaena*. These two species are also morphologically differentiated, with *G. mesotriaena* having longer oxeas, longer plagiotriaene rhabds (but not longer clads), and larger sterrasters (table 2, see also *G. agassizii* remarks). The only species in the region known to have megascleres as long as *G. mesotriaena* are *G. ovis* (which has smaller sterrasters, shorter anatriaene clads, and shorter mesotriaene epirhabds) and *G. starki* (which has longer megascleres, no mesotriaenes, and two size classes of sterrasters). Further morphological differences pointed out by Lendenfeld (1910) include that *G. agassizii* and *G. ovis* both lack the preoscular cavities seen in *G. mesotriaena* (this was not investigated here). He also found that the small asters of these species had subtle differences in size and shape, though I was unable to differentiate them based on the asters.

Lendenfeld erected three varieties of this species, one for each sample he examined (Lendenfeld 1910). The primary difference between them was in their anatriaene clad sizes: var. *pachana* clads were short and thick, var. *microana* clads were short and thin, and var. *megana* were long and thick. These differences are reproduced in the new spicules isolated and measured from the type samples, but increasing the number of samples from three to seven makes clad length and width look like continuous spectra of variation rather than categorical types. I therefore think it is unlikely that these varieties will prove to be valid taxonomic units. No DNA differences were seen between the holotypes of the three varieties, though only a very short region of a single gene was successfully sequenced from the types. Longer sequences were generated from the samples collected by the author, including a complete nuclear ribosomal sequence generated from Illumina data.

### *Geodia ovis* Lendenfeld, 1910

Figure 25

**Figure 25.**
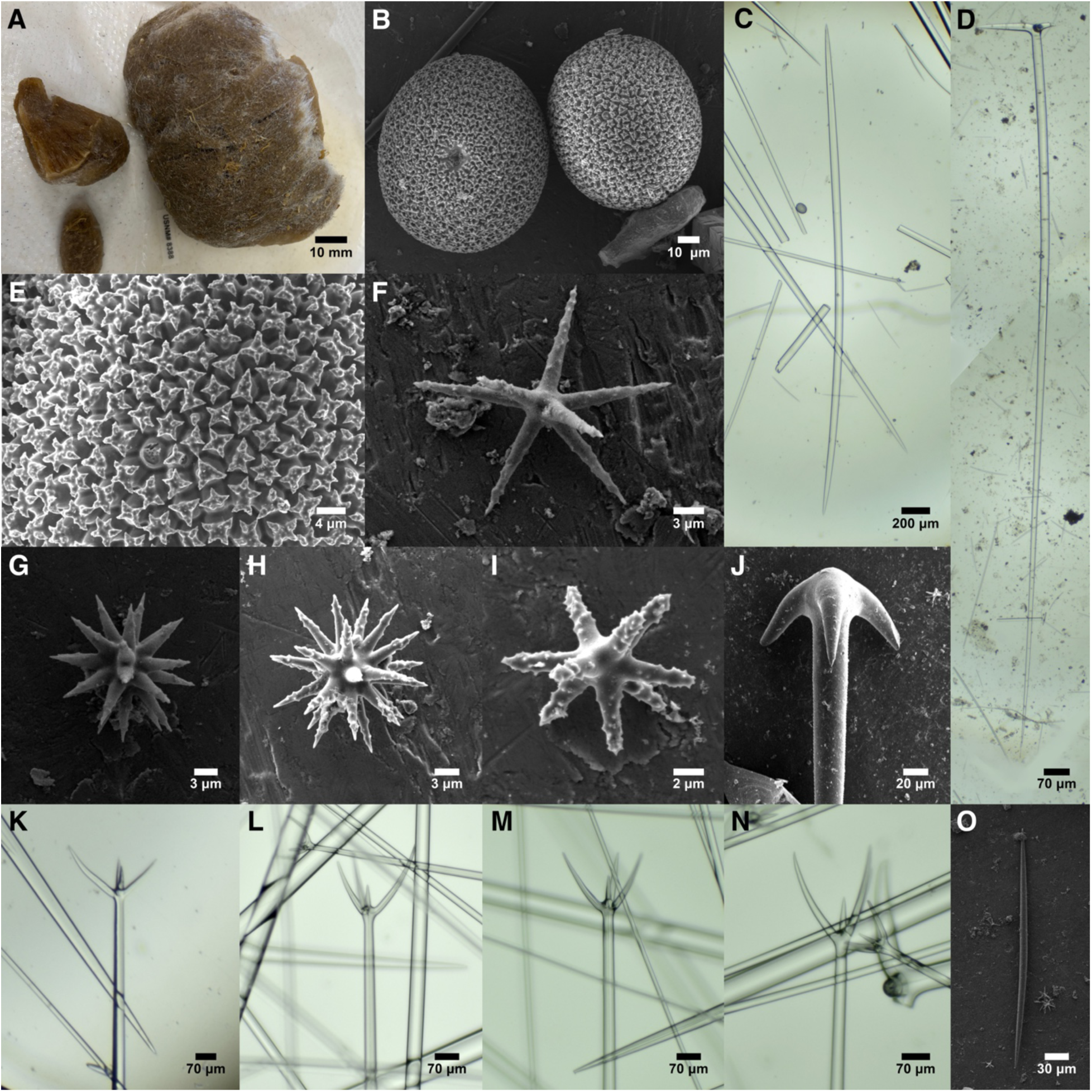
Geodia ovis. A: Preserved holotype; the smaller fragments were loaned to me and reexamined here. B: Sterrasters. C: Oxea I. D: Plagiotriaene. E: Sterraster detail. F: Large thin-rayed aster. G-H: Many-rayed large thick-rayed asters. I: Small aster. J: Anatriaene. K-N: Example mesoprotriaenes showing short epirhabds. O: Short oxea. All images from holotype and only known sample.

#### Material examined

Holotype: USNM 8388, W of Anacapa Isl., California, (34.02500, - 119.48300), 66 m, 12-Feb-1889.

#### Morphology

Known from only one sample, and only a fragment was collected. The fragment was 12.7 × 5.0 × 2.7 cm, but proposed to have come from a cake-shaped sponge approximately 4 cm thick (Lendenfeld 1910). Said to have especially long and thick spicule fur, up to 2 cm thick, thus the name; the original description says the sample is light brown, though it is now dark brown. Oscula not observed. See Lendenfeld (1910) for further details.

#### Skeleton

Described in detail by Lendenfeld (1910), and not differing substantially from the other *Geodia* with mesotriaenes included here. Briefly, the choanosome contains radial bundles of oxeas I, with some piercing the surface to contribute to spicule fur. Plagiotriaenes are added to bundles near the surface, with their cladomes located just below the sterraster layer in the cortex. Anatriaenes also occur near the surface, with their clads in the cortex or protruding beyond it. Mesoprotriaenes are mostly found piercing the surface of the sponge and are the primary component of the spicule fur. Sterrasters densely populate the central cortex, with a few in the choanosome. Oxyasters are found throughout, with larger asters in the choanosome and smaller ones in the cortex. The outer cortex is dense with small asters, most said to be strongylospherasters. The short oxeas form upright bouquets in the dermal layer, but are also scattered throughout the choanosome.

#### Spicules

Shown in figure 25 except where noted. See table 2 for comparisons in average values among species.

Oxeas I: stout; most taper to points at both ends, but occasional styles or tylostyles occur. 1294–5584–8423 × 24–65–98 μm (n=30). Reported as 4000–9000 × 30–100 μm in the type description (Lendenfeld 1910).

Short oxeas: straight or slightly curved; most are symmetrical or very nearly so, but some have one rounded tip and one pointed tip. 264–321–393 × 7–10–13 μm (n=20). Reported as 270–550 × 8–13 μm in the type description (Lendenfeld 1910).

Plagiotriaenes: with long rhabds usually tapering to sharp points, but strongylote rhabds or clads are found as a minority. The clads are frequently unequal in length, and only the longest clad was measured. Rhabds 2586–5071–7349 μm (n=25) × 31–68–93 μm (n=28). Clads 130–348–612 (n=27) × 29–57–86 (n=30) μm. Reported with rhabds 5000–8000 × 77–110 μm and clads 310–640 μm in the type description (Lendenfeld 1910). The rhabd:clad angles were 93°–103°–112° (n=103), similar to the 86°–94°–109° previously reported; these triaenes were referred to as ortho/plagiotriaenes in the type description (Lendenfeld 1910).

Mesoprotriaenes/protriaenes/mesoprodiaenes/mesopromonaenes: Mesoprotriaenes with three curved clads and a straight central epirhabd predominate, but protriaenes (lacking the epirhabd), or mesoprodiaenes/mesopromonaenes (appearing to have the epirhabd but lacking one or more clads) also occur. In mesoprotriaenes, the epirhabd is usually shorter than the longest clad but is equal in some spicules. Only one unbroken rhabd was found, 8784 μm long; widths 13–23–35 μm (n=18); epirhabds are 33–80–122 μm (n=16), and the longest clads are 64–148–265 μm (n=19) × 9–17–29 μm (n=19) μm. For mesoprotriaenes alone, the epirhadb-to-longest-clad ratios were 0.31–0.54–1.02 (n=16). Lendenfeld reported rhabds 6000–17000 × 20–41 μm, epirhabds 70–110 μm, and clads 40–360 μm in the type description (Lendenfeld 1910). The rhabd:clad angles of newly isolated spicules were 28°–46°–63° (n=67); previously reported as about 45° (Lendenfeld 1910).

Anatriaenes/anadiaenes/anamonaenes: Rhabds are long and usually taper to a filament-like ending. Most are anatriaenes, but some anadiaenes/anamonaenes seen. Measurements are highly variable and nearly continuous, but rhabd widths were bimodal in the newly measured spicules and also reported as such in type description. I therefore divide them into large and small classes here.

Large anatriaenes: One complete rhabd was measured at 12622 μm; widths 11–24–37 μm (n=28); clads are 52–124–207 (n=28) × 12–28–46 (n=28) μm, and rhabd:clad angles are 29°–38°–52° (n=28). Lendenfeld (1910) reported rhabds up to 23000 × 17–45 μm, clads 70–205 μm, and rhabd:clad angles of 36°–42.5°–55°.

Small anatriaenes: No unbroken rhabds were seen, but widths are 5–8–10 μm (n=4); clads are 21–28–33 (n=3) × 5–8–11 (n=4) μm, and rhabd:clad angles are 36°–41°–44° (n=3). Lendenfeld (1910) reported rhabds up to 670–2500 × 2–7 μm, clads 6–43 μm, and rhabd:clad angles of 41°–65°. Lendenfeld (1910) reports that some of these were extruded into the spicule fur, and therefore not still developing into larger anatriaenes. Not shown.

Sterrasters: Nearly round when seen from above, with length:width ratios of 1.1. Rosettes are warty, with 3–7 spines per rosette. The number of spines per rosette are 4–5–6 when 10 or more rosettes are measured and averaged per sterraster; n=4 sterrasters. Mature sterrasters (those with rays ending in rosettes) are 73–100–125 (n=718) × 67–71–75 (n=27) × 55–58–61 (n=3) μm.

Reported as 82–92 × 70–83 × 54–61 μm in the type description; see Lendenfeld (1910) for additional details, including discussion of rare forms he dubs "sterroids".

Asters: All asters are acanthose, and most are clearly oxyasters. While he noted that they form a continuous spectrum, Lendenfeld still divided them into large thin-rayed oxyasters, many-rayed large thick-rayed oxyasters, and small thick-rayed asters. While asters are greatly varying in size, spine number, and ray thickness, I found it difficult to categorize them, and all are pooled here.

Newly measured spicules are 8–16–25 μm (n=68) in total diameter. The distribution of diameter is continuous but multi-modal, with a major mode at 18 μm and smaller modes at 11 and 25 μm. Previously reported as 20–34.5 μm for large thin-rayed asters, 28–45 μm for many-rayed large thick-rayed asters, and 11–24 μm for small thick-rayed asters (Lendenfeld 1910).

#### Distribution and habitat

The only known sample was trawled from 66 m depth in Southern California, West of Anacapa Island. It was found growing on a bed of gravel and broken shells (Lendenfeld 1910).

#### Remarks

Though very similar to *G. mesotriaena*, Lendenfeld distinguished the one sample he attributed to this species by its lack of preoscular cavities and having thicker spicule fur, smaller sterrasters, and a small size class of anatriaenes. I found a single small anatriaene in one sample of *G. mesotriaena*, so this character is somewhat suspect. I also doubt that the thickness of the spicule fur is a reliable taxonomic character. However, I was able to replicate the difference in sterraster diameter; when this is combined with the long megascleres in this sample, it is distinguishable from both *G. mesotriaena* and *G. agassizii* (table 2, figure 23). Also, though not mentioned as a distinguishing feature by Lendenfeld, his measurements and mine indicate that the mesoprotriaenes in this sample have short epirhabds, with epirhabd:clad ratios averaging 0.54, similar to *G. mesotriaenella* but differing from *G. mesotriaena* (1.13) and *G. agassizii* (1.10). Lendenfeld also felt that it was distinguished by several traits in the asters, with smaller dermal asters and "the large oxyasters with stout, regularly conical, and sharp-pointed rays, so abundant in *G. ovis*, are wanting in *G. mesotriaena*". I did not examine aster type in specific tissues, and cannot speak to the differences in dermal asters. I did find that the many-rayed asters with conical spines were more abundant in *G. ovis* than *G. mesotriaena*, but they are present in both, and the large variation in aster shapes and sizes made quantifying types difficult.

Unfortunately, I was unable to amplify DNA to confirm that this sample is a unique species. However, the multiple spicular traits that differentiate this sample in both Lendenfeld’s analysis and mine indicate that it is morphologically distinguishable from the other *Geodia* in the region. It is possible that a larger morphological dataset will show that there is a continuous range of variation in *G. mesotriaena* that encompasses this sample, but until such data are found, I suggest retaining the name *G. ovis* for this morphotype.

### Geodia mesotriaenella Lendenfeld, 1910

Figure 26

**Figure 26.**
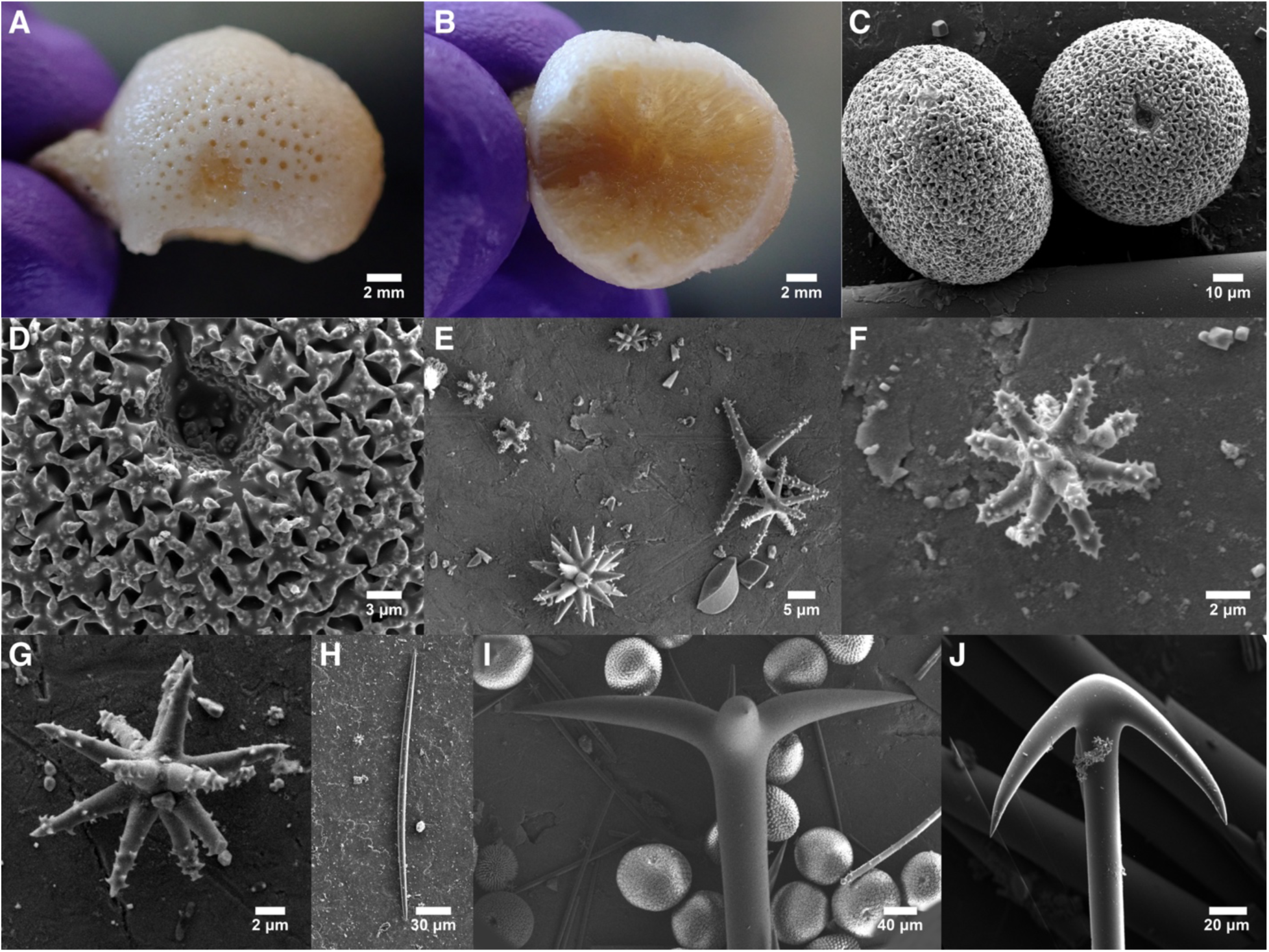
Geodia mesotriaenella. A-B: Preserved holotype. C-D: Sterrasters. E: Small strongylasters (upper left) with large oxyasters of varying shape. F: Small strongylaster. G: Oxyaster. H: Short oxea. I: Plagiotriaene. J: Anatriaene. All are from holotype except J, which is from USNM 8367, the former G. breviana holotype.

#### Synonyms

*Geodia breviana* Lendenfeld, 1910

#### Material examined

Holotype: USNM 8387, NE of Santa Barbara Isl., California, (33.45000, - 119.08333), 53 m, 12-Apr-1904. Holotype of *G. breviana*: USNM 8367, NW of San Miguel Isl., California, (34.11670, -120.55800), 97 m, 5-Jan-1889.

#### Morphology

The holotype is a small sphere, 19 × 15 mm, with only a small amount of spicule fur near the base. Oscules are numerous and small, approximately 400 μm across. White preserved. Samples previously assigned to *G. breviana,* which is synonymized here, also include a thick-walled cup-shaped sponge 6 cm high (not examined here). See Lendenfeld (1910) for further details.

#### Skeleton

Described by Lendenfeld (1910), and not differing substantially from the other *Geodia* with mesotriaenes included here. The choanosome contains radial bundles of oxeas I, with some piercing the surface to contribute to spicule fur. Plagiotriaenes are added to bundles near the surface, with their cladomes located within the cortex. Anatriaenes also occur near the surface, with their clads in the cortex or protruding beyond it. Mesoprotriaenes are mostly found with their clads piercing the surface of the sponge. Sterrasters densely populate the central cortex, with some in the choanosome. Oxyasters are found throughout the choanosome and in the inner cortex. The outer cortex is densely populated with strongylospherasters. The short oxeas are found in the dermal layer, upright with their tips piercing the surface.

#### Spicules

Shown in figure 26 except where noted. See table 2 for comparisons in average values among species. A few spicules measured here are from the slides prepared and archived by Lendenfeld, but most were isolated from new spicule preparations.

Oxeas I: stout; most taper to points at both ends, but occasional styles or tylostyles occur. 801–1765–2560 × 12–37–62 μm (n=88). No significant difference seen between the *G. mesotriaenella* holotype (mean 1807 μm) and the *G. breviana* holotype (mean 1720 μm).

Reported as 2000–2600 × 20–50 μm in the type description (Lendenfeld 1910). Not shown in figure 26; see figure 25C for similar example.

Short oxeas: straight or slightly curved; many are asymmetrical due to tapering more gradually at one end, but symmetrical oxeas also seen. 166–341–581 × 5–8–11 μm (n=51). Significantly longer in the *G. mesotriaenella* holotype (mean 386 μm) and the *G. breviana* holotype (mean 294 μm). Reported as 196–260 × 4–5 μm in the type description (Lendenfeld 1910).

Plagiotriaenes: with long rhabds usually tapering to sharp points. The clads are frequently unequal in length, and only the longest clad was measured. Rhabds 517–1766–2481 μm (n=68) × 18–68–113 μm (n=72). Clads 75–322–589 (n=64) × 14–53–100 (n=72) μm. Reported with rhabds 2100–2400 × 75–120 μm and clads 350–600 μm in the type description (Lendenfeld 1910). The rhabd:clad angles were 94°–108°–119° (n=58), similar to the 90°–93°–96° previously reported; referred to as orthotriaenes in the type description (Lendenfeld 1910). All measurements were extremely similar in the *G. mesotriaenella* and *G. breviana* holotypes.

Mesoprotriaenes: With three curved clads and a straight central epirhabd; the epirhabd is shorter than the longest clad. Few of these spicules could be found in new spicule preparations, which were limited by the small size of the subsamples loaned and the lack of protruding spicule fur upon them. Only one was found on the existing slides archived by Lendenfeld, and one new example was isolated from the *G. mesotriaenella* holotype. One complete rhabd was measured at 2106 μm long; widths 13–14–15 μm (n=2); epirhabds are 79–92–106 μm (n=2), and the longest clads are 120–135–149 μm (n=2) × 10–11–11 μm (n=2) μm. The epirhabd to longest clad ratios were 0.66–0.68–0.71 (n=2). Lendenfeld reported rhabds 2800–3400 × 9–19 μm, epirhabds 70–165 μm, and clads 100–220 μm in the type description (Lendenfeld 1910). The epirhabd:clad length ratios were previously reported as 0.43-0.77. The rhabd:clad angles of newly isolated spicules are 20°–27°–35° (n=2); previously reported as 30°–36°–47° (Lendenfeld 1910). Not shown in figure 25; see figure 25 K-N for similar examples.

Anatriaenes/anadiaenes/anamonaenes: Rhabds are long and usually taper to a filament-like ending. Most are anatriaenes, but some anadiaenes/anamonaenes seen. Measurements are highly variable and continuous, but spicules were divided into size classes based on rhabd widths here.

Large anatriaenes: Rhabds 2384–2867–3284 μm (n=7) × 11–23–37 μm (n=40); clads are 58–106–147 (n=38) × 12–25–40 (n=40) μm, and rhabd:clad angles are 32°–39°–49° (n=28).

Lendenfeld (1910) reported rhabds up to 3700 × 18–30 μm, clads 87–140 μm, and rhabd:clad angles of 41°–48°–57°. Clad lengths and clad angles were the same in the *G. mesotriaenella* and *G. breviana* holotypes, but there are slight, but significant, differences in rhabd thickness (18 vs. 25 μm) and clad thickness (22 vs. 27 μm), both of which are smaller in the *G. mesotriaenella* holotype.

Small anatriaenes: Not seen in the *G. mesotriaenella* holotype, but present in the *G. breviana* holotype. Rhabds 1996–2360–2770 μm (n=3) × 5–8–10 μm (n=6); clads are 21–31–47 (n=5) × 5–9–11 (n=6) μm, and rhabd:clad angles are 39°–49°–55° (n=5). Lendenfeld (1910) reported none in the *G. mesotriaenella* holotype, but even smaller spicules in the *G. breviana* holotype. They were found to have rhabds 350–610 × 1–4.5 μm, clads 2–12 μm, and rhabd:clad angles of 42°–60°; anadiaenes and anamonaenes were seen among them. Not shown.

Sterrasters: Nearly round when seen from above, with length:width ratios of 1.1. Rosettes are warty, with 3–7 spines per rosette. When averaging 10 or more rosettes per sterraster, the number of spines per rosette is 4–5–6 (n=5 sterrasters). Mature sterrasters (those with rays ending in rosettes) are 63–82–103 (n=181) × 61–75–88 (n=56) × 49–57–68 (n=11) μm. No differences were seen between the *G. mesotriaenella* and *G. breviana* holotypes. Reported as 87–107 × 70–92 × 58–69 μm in the type description.

Oxyasters/oxyspherasters: Asters with sharply pointed rays, varying from sparsely acanthose to heavily spined, and varying from many conical rays to fewer long thin rays. Lendenfeld divides them into oxyasters and oxyspherasters. I found this division to be difficult, and all are pooled here. Newly measured spicules are 13–18–29 μm (n=83) in total diameter, with no differences between *G. mesotriaenella* and *G. breviana* holotypes. Reported in the type description as 17–26 μm for oxyasters and 20–21 μm for oxyspherasters; see Lendenfeld (1910) for additional details.

Strongylasters/strongylospherasters: Called strongylospherasters by Lendenfeld, but more easily differentiated from oxyasters by size than shape. Small and heavily spined asters, with rays that are fairly straight and blunt, but most are arguably oxyasters due to pointed spines on the ends of the rays. Newly measured spicules are 5–7–10 μm (n=60) in total diameter, with no differences between *G. mesotriaenella* and *G. breviana* holotypes. Lendenfeld states that strongylospherasters are larger in *G. breviana,* but this is not clear based on the numbers he reports. The two *G. breviana* samples were said to have total diameters of 6–9 μm and 7–12 μm vs. 6–11 μm in *G. mesotriaenella* (Lendenfeld 1910).

Oxyasters/strongylasters combined: defining classes of asters based on ray shape was difficult, but there were two clear classes based on size. If considered together, total diameters of all asters are 5–13–29 μm (n=121).

#### Distribution and habitat

Two samples are known from near the Northern Channel Islands in Southern California, from 53–97 m depth. A third sample was collected near Comox, near Vancouver Island, British Columbia, at 7 m.

#### Remarks

When describing *G. mesotriaenella* and *G. breviana*, Lendenfeld stated that "in *G. breviana* minute, dermal anaclades are present, the clades of the large anatriaenes are much stouter and the strongylosphaerasters larger" compared to *G. mesotriaenella* (Lendenfeld 1910). In the newly isolated spicules measured here, the small anatriaenes were again found only in *G. breviana.* However, Lendenfeld found these in *G. ovis* and *G. agassizii*, and I found one in a sample of *G. mesotriaena*. It seems possible that these are either variable across these species and/or immature spicules of the larger anatriaenes. I also did not find any differences in the size of the smallest asters (p = 0.424), and the sizes reported by Lendenfeld appear to be the same, so this reported difference may have been an error. I did, however, replicate the difference in the thickness of the larger anatriaene clads (p = 0.015); I also found the short oxeas to be shorter in *G. breviana* (p=0.0002). These differences are slight, however, and I think they are unable to support the concept of two species, when based on so few samples. In contrast to the other mesotriaene-bearing *Geodia*, the principal components analysis of multiple variable characters also failed to distinguish the holotypes (figure 23). I therefore propose that *G. breviana* be considered a junior synonym of *G. mesotriaenella*. I was unable to amplify DNA from either holotype.

Lendenfeld acknowledges that these samples may be small, young individuals of another species, and thinks *G. agassizii* or *G. mesotriaena* would be the most likely conspecifics. I also think this is possible, maybe even probable, but disagree about the likely congener. Based on sterraster size and having mesotriaenes with epirhabds shorter than clads, these samples seem to be more closely allied to *G. ovis* than to *G. agassizii* or *G. mesotriaena*. Despite this possibility, I recommend retaining the name *G. mesotriaenella* until new samples can be found and genotyped, or alternative methods are able to extract usable DNA from the type samples.

### *Penares cortius* de Laubenfels, 1930

Figure 27

**Figure 27.**
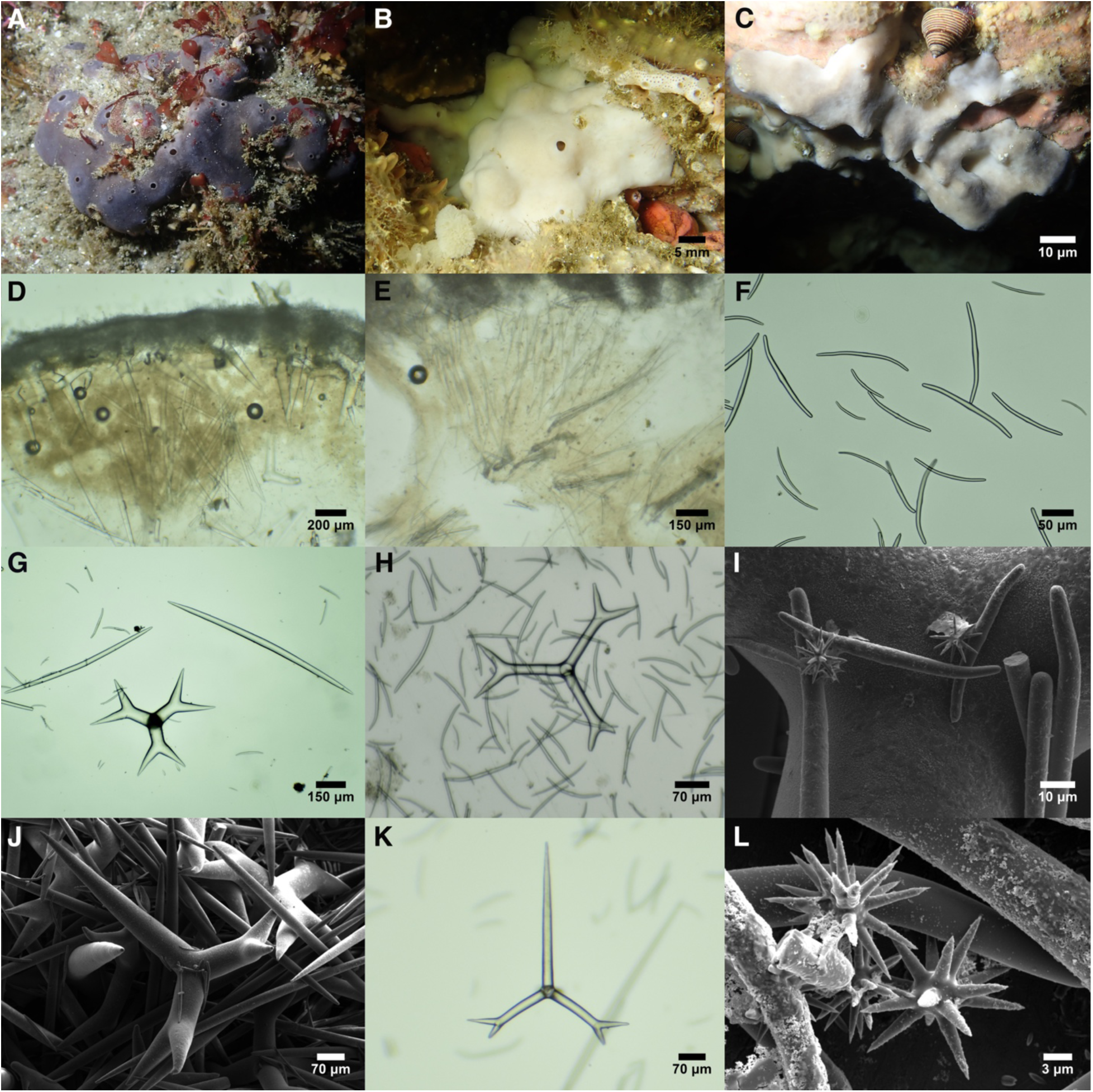
Penares cortius. A: TLT946 in situ, showing the gray surface of samples exposed to sunlight. B: TLT468 in situ, showing the white surface of samples growing in caves. C: TLT368 in situ, growing under a ledge in an intermediate condition to A and B. D: Perpendicular section at the sponge surface, showing the dense surface layer of microstrongyles supported by triaenes, with oxeas in confusion; from TLT368. E: Perpendicular section showing oxeas in confusion but also in tracts; the surface layer of microstrongyles is visible at the top of the image; from TLT1497. F: Microstrongyle variation in TLT325. G: Thick-rayed dichotriaene cladome with oxeas from TLT368. H: Thin-rayed dichotriaene cladome from TLT468. I: Microstrongyles with asters from TLT368. J: Thick-rayed dichotriaene from TLT368. K: Thinner dichotriaene from TLT118. L: Asters from TLT368.

#### Material examined

TLT368, Middle Reef, Point Lobos, Monterey, California, (36.52172, - 121.93894), 6-15 m, 23-Nov-2019; TLT67, Landslide cove, Anacapa Isl., California, (34.01657, -119.36126), 7-11 m, 25-Apr-2019; TLT946, La Jolla Cove Reef, San Diego, California, (32.85227, -117.27239), 10-16 m, 17-May-2021; USNM 31321, San Clemente Isl., California, (33.00170, -118.55100), low intertidal, 20-Dec-1976; TLT1548, Sealion Cave, Anacapa Isl., California, (34.01642, -119.42679), 4.6 m, 8-Feb-2026; TLT1247, Naples Reef, Santa Barbara, California, (34.42212, -119.95154), 6-16 m, 29-Oct-2021; TLT325, Naples Reef, Santa Barbara, California, (34.42212, -119.95154), 6-16 m, 26-Sep-2019; TLT118, Naples Reef, Santa Barbara, California, (34.42212, -119.95154), 6-16 m, 31-Jul-2019; 15517 & 15518, Marineland, Los Angeles, California, (33.75200, -118.41700), depth not known, 21-Aug-2019; TLT437, Prince Island, San Miguel Isl., California, (34.05700, -120.33109), depth not recorded, 8-Oct-2019; TLT468, Big Rock, Santa Cruz Isl., California, (34.05220, -119.57360), 6-14 m, 19-Jan-2020; TLT56, West End, Anacapa Isl., California, (34.01352, -119.44570), 9-12 m, 25-Apr-2019; TLT1466, Daytona, San Nicolas Isl., California, (33.21688, -119.44412), 9-14 m, 9-Oct-2024; TLT1497, Point Dume, Ventura, California, (33.99837, -118.80735), 7-17 m, 9-Nov-2024.

#### Morphology

Small sponges are often roughly spherical and cushion-shaped, but sometimes irregular in outline. Larger samples have undulating surfaces and highly variable thickness, with oscula mostly atop elevations. The holotype was 4 cm thick and 10 cm in diameter; most new samples collected were not measured in situ but appeared smaller than the holotype. Oscula are not grouped, and are scattered across the surface, with slightly raised rims, circular to oval, 1.0–2.5 mm in diameter. On some large samples, oscula are atop raised mounds; in rare cases, each mound has 3-4 small oscula in groups. Sponges collected from caves or under ledges are white inside and outside, but sponges exposed to more direct light have a dark gray surface and white interior; colors remain after preservation in ethanol. Surface is smooth and leather-like.

#### Skeleton

The ectosomal skeleton is densely populated with tangential microstrongyles. Asters are visible just below this layer, but may also occur within it and be obscured by the strongyles. The ectosome is supported by the cladomes of the dichotriaenes, with the rhabds pointing inward; a few dichotriaenes are also scattered within the choanosome. The choanosomal skeleton includes ascending tracts 2–10 oxeas wide but also has many oxeas in confusion. Some scattered microstrongyles are also found in the interior.

#### Spicules

Shown in figure 27 except where noted. See table 3 for a comparison of spicules among *Penares* species.

**Table 3.** Key features distinguishing species of *Penares* in the region. Range of means across samples is shown, in microns.

|  | Oxea length | Microxea /<br>strongyle<br>length | Dichotriaene<br>rhabd length | Dichotriaene<br>clad length | Aster<br>diameter |
| --- | --- | --- | --- | --- | --- |
| *P. orientalis* | 1178 - 1712 | 83 - 103 | 584 - 941 | 154 - 257 | 24 - 40 |
| *P. cortius* | 692 - 1046 | 89 - 115 | 242 - 526 | 135 - 327 | 9 - 16 |
| *P. anyapax* | 530 - 724 | 67 - 84 | 186 - 251 | 168 - 191 | 12 - 19 |
| *P. foxi* | 577 | 71 | 225 | 224 | 15 |
| *P. saccharis* | 696- 788 | 89 - 120 | 274 - 331 | 239 - 270 | none |

Oxeas/strongyloxeas/styles: the primary choanosomal spicules are diactinal, but vary on a continuum from oxeas with sharply pointed tips, to strongyloxeas with one or both tips blunt and rounded, to styles with one end untapered and the other end pointed. All are about the same size and are pooled here. 340–828–1245 (n=286) × 4–16–37 (n=200) μm. Oxea shown in figure 27; for a more strongylote example, see figure 28E.

**Figure 28.**
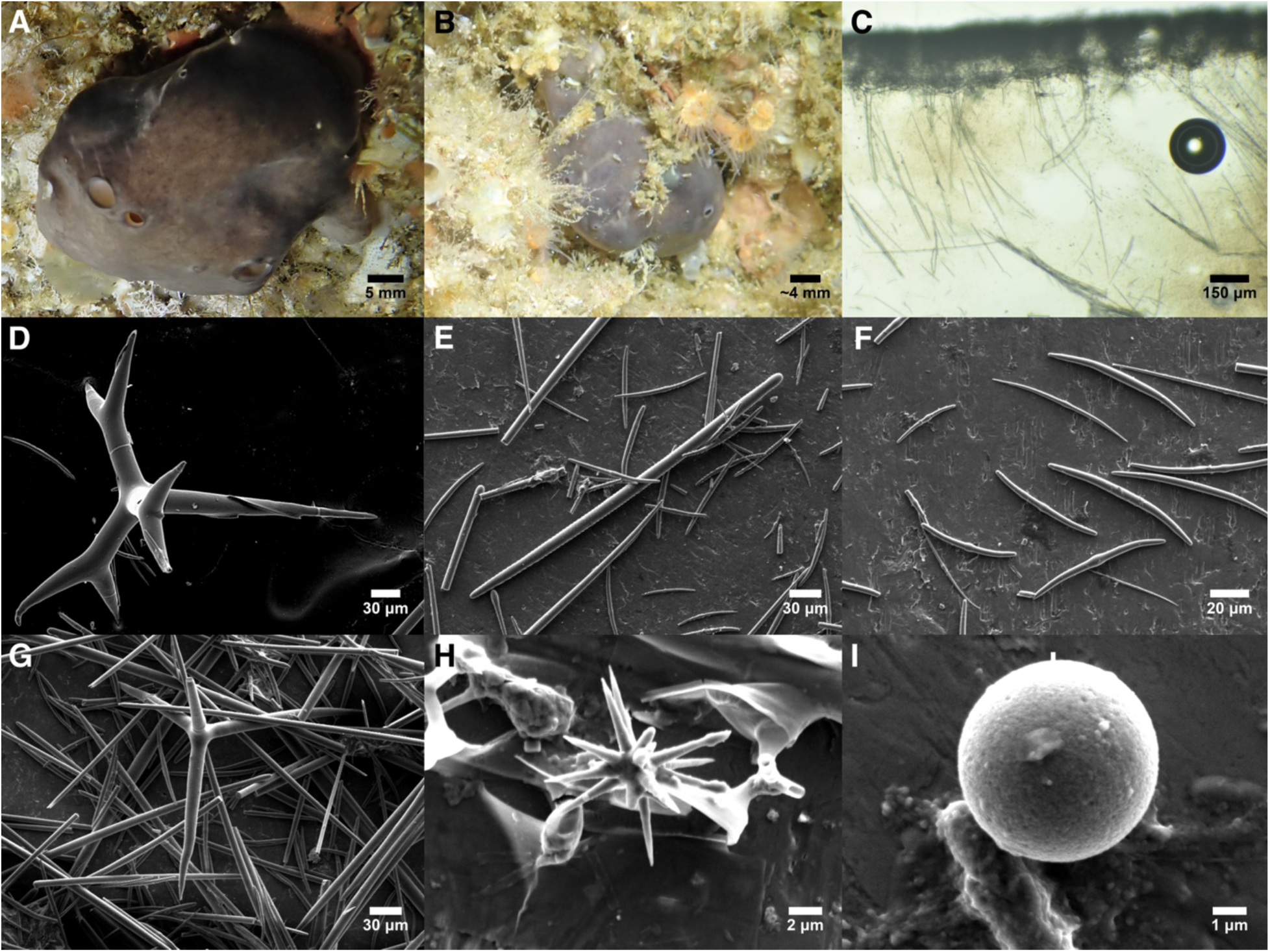
Penares anyapax. A: Holotype in situ. B: TLT61 in situ; scale is approximate. C: Perpendicular section at sponge surface, showing surface crust of microstrongyles supported by triaenes, with single oxeas and tracts of oxeas in the choanosome below; from holotype. D: Dichotriaene. E: Strongylote megasclere with microstrongyles. F: Variability in microstrongyles, including several that are relatively oxeote. G: Plagiotriaene with broken rhabd. H: Oxyaster. I: Sphere. All SEM images from TLT75.

Dichotriaenes/plagiotriaenes: Dichotriaenes with long pointed rhabds and three forked clads are present in all samples, but simple triaenes with no forked clads are also common in many samples. Irregular forms with some clads forked and some unforked, having irregular bends and twists, or with strongylote rhabds and/or clads are common in many samples. The clads are sometimes unequal in length, and only the longest was measured. Rhabds 144–380–683 μm (n=102) × 5–30–93 μm (n=103). Clads 91–239–528 (n=177) × 7–25–88 (n=136) μm. The one sample from Central California (TLT368) had larger dichotriaenes than the Southern California samples, especially in terms of width (mean clad width 68 μm for sample TLT368, 12–34 μm for the 10 measured samples from Southern California). Sample TLT368 alone: rhabds 377–526–614 μm (n=13) × 60–75–93 μm (n=13). Clads 238–327–528 (n=28) × 46–68–88 (n=28) μm. All Southern California samples measured: rhabds 144–359–683 μm (n=89) × 5–24–45 μm (n=90). Clads 91–223–381 (n=149) × 7–20–48 (n=121) μm. The type description states that rhabds are up to 400 × 50 μm, clads up to 310 × 50 μm. Plagiotriaene not shown in figure 27; for similar example, see figure 28G.

Microstrongyles: smooth; thickness tapers somewhat at the ends but most end in blunt, rounded tips. A small minority are oxeas, generally thinner than strongyles, and gradually tapering to sharply pointed tips. Variable in shape, but many are slightly thickened in the center and bent at each end to form the "bicurvate microstrongyles" described in the type description (de Laubenfels 1932). However, many are gently curved, straight, or with weak bends. Many (especially the smaller ones) have a centrotylote ball rather than just merely a slight centrotylote swelling. Variable in length, but the distribution is continuous without clear size classes. When all spicules are pooled, the length distribution is slightly trimodal, with modes at about 60, 100, and 165 μm. Small, medium, and large spicules were found in every sample, with minimum lengths per sponge varying from 28–62 μm, mean lengths per sponge varying from 89–125 μm, and maximum lengths per sponge varying from 151–204 μm. All spicules pooled: 28–98–192 (n=621) × 2–5–12 (n=483) μm.

Oxyspherasters: smooth, with a round centrum and many (∼10–20) conical, sharply pointed rays. Common in some samples but rare in others; at least one was found in every sample, but considerable searching was sometimes required. Total diameter 4–12–23 μm (n=82).

Spheres: simple glass spheres are present in low numbers; only a few were measured, and they were 5–25 μm in diameter. Not shown in figure 27; see figure 28I for similar example.

#### Distribution and habitat

This species ranges from Carmel Bay, Central California, to San Diego, Southern California, and likely into temperate Mexican waters. The type samples were collected in the intertidal, but I know of no other intertidal samples. I did not find the species at any of the 14 intertidal sites I surveyed in California, and no pictures posted on the site iNaturalist.org show intertidal *Penares* in the region. The deepest sample I have verified as this species was collected between 10–16 m.

*P. cortius* has previously been said to range as far north as the Gulf of Alaska (Austin 1985; Stone *et al*. 2011), but the samples I examined from Oregon, British Columbia, and Alaska are here reassigned to *P. orientalis* comb. nov. (see below). More samples should be examined to determine the Northern range limit of *P. cortius*, but its distribution within California suggests it is a warm-water affiliated species. Though it was described from Carmel, in Central California, I found it at only 2/18 subtidal sites surveyed in Central California, both of which were in Carmel Bay. In Southern California, *Penares* sp. were more common in the warmer waters of Anacapa Island, Catalina Island, Los Angeles, and San Diego, and less common in the colder waters of Santa Barbara and the other Northern Channel Islands (many of these samples were identified in photo surveys only, which can’t differentiate between *P. cortius*, *P. anyapax* sp. nov., and *P. foxi* sp. nov.). *Penares* sp. in Southern California also appear to favor clearer, less silty conditions, as they were common on the island side of the Santa Barbara Channel, but absent on the mainland side, except for Naples Reef, which has the clearest water on the mainland side of the channel.

The Southern range limit of *P. cortius* should also be investigated further. The species likely extends into the Temperate Mexican Pacific, but the only published record from Baja California does not include a morphological description (Dickinson 1945). It has also been said to be in the Tropical Mexican Pacific (Carballo & Vega 2022; Gómez *et al*. 2002), but this is unlikely, as explained below.

#### Remarks

Genotyping of *Penares* in the region revealed a higher-than-expected diversity. Two species were previously known, and validated here, but an additional three species were found. It is therefore important that the correct genotype is associated with the name *P. cortius.* I did not examine the type material of this species, which was collected in the intertidal at Pescadero Point, Carmel Bay, California. However, sample TLT368 was collected from Point Lobos, which is also within Carmel Bay, and is a good morphological fit to the type description. As I have been unable to successfully sequence DNA from any of de Laubenfels’ vouchered material, this sample is a good voucher for associating the name with a genotype.

*P. cortius* is distinguished from *P. anyapax* sp. nov. and *P. foxi* sp. nov. by having longer oxeas and longer microstrongyles. It is distinguished from *P. orientalis* comb. nov. by having smaller spicules of all types, especially the oxyasters, where the mean values in all *P. orientalis* comb. nov. are larger than the maximum values seen in *P. cortius.* The final species, *P. saccharis*, is distinguished by having no asters, but this is a difficult taxonomic character because asters were rare and hard to find in some *P. cortius* samples. It is also distinguished by having primarily microxeas instead of primarily microstrongyles.

*P. cortius* appears to be the most common and widespread *Penares* species in California. *P. saccharis* is known only from the intertidal zone on the Monterey Peninsula, where it co-occurs with *P. cortius*. The other Californian species were only found at Anacapa Island; of the eight samples collected around the island, 50% were *P. anyapax* sp. nov., 37.5% were *P. cortius*, and 12.5% were *P. foxi* sp. nov.

Gómez *et al*. (2002) describe a sponge from Guerrero, Mexico as *P. cortius*. However, the asters revealed by their SEM images can now be compared to the SEM images generated here and by Lee et al. (2007), and they are clearly different. The tropical sample has acanthose asters which also have fewer and thicker rays. The oxeas are also considerably longer in this tropical sample, which is likely from an undescribed species.

### Penares anyapax sp. nov

Figure 28

#### Material examined

Holotype, TLT1549, Goldfish Bowl, Anacapa Isl., California, (34.01539, - 119.43560), 3-11 m, 8-Feb-2026. Paratypes, TLT61, TLT75, and TLT76, all collected at West End, Anacapa Isl., California, (34.01352, -119.44570), 9-12 m, 25-Apr-2019.

#### Etymology

Name inspired by the island on which it was found, Anacapa Island, which is itself derived from the Chumash name ’anyapax. The name means "mirage", because when the island is viewed from the mainland, it is frequently distorted due to atmospheric phenomena.

#### Morphology

Cushion-shaped to irregular in outline. The holotype is the largest sample, and was approximately 5 cm in horizontal extent; the sampled portion is 2 cm thick, but the sponge was approximately twice as thick in life. Oscula are single and scattered across the surface, with slightly raised rims, circular to oval, 0.5–4.5 mm in diameter. All known samples have a dark gray surface and white interior, and these colors remain after preservation in ethanol. Surface is smooth and leather-like.

#### Skeleton

The ectosomal skeleton is densely populated with tangential microstrongyles, some of which are also scattered about the interior. The ectosome is supported by the cladomes of the dichotriaenes, with the rhabds pointing inward; a few dichotriaenes are also seen in the choanosome. The choanosomal skeleton includes tracts of oxeas 2–10 spicules wide which are mostly parallel to the sponge surface, and also many oxeas and microstrongyles in confusion. The location of the asters was not determined in tissue sections, as they are small, sparse, and difficult to distinguish from sand grains.

#### Spicules

Shown in figure 28 except where noted. See table 3 for a comparison of spicules among *Penares* species.

Oxeas/strongyloxeas/styles: the primary choanosomal spicules are diactinal, but vary continuously from oxeas with sharply pointed tips, strongyloxeas with one or both tips blunt and rounded, or styles with one end untapered and the other end pointed. Holotype: 356–616–951 (n=23) × 5–9–17 (n=23) μm. All samples pooled: 301–630–1069 (n=109) × 4–10–18 (n=97) μm. Strongylote example shown in figure 28E; see figure 27G for similar example oxeas.

Dichotriaenes/plagiotriaenes: Dichotriaenes with long pointed rhabds and three forked clads are present in all samples, but simple triaenes with no forked clads are also common. Irregular forms are present, with only some clads forked, with irregular bends, or with strongylote rhabds and/or clads (strongylote rhabds are especially common in the holotype). The clads are sometimes unequal in length, and only the longest was measured. Holotype: rhabds 138–251–342 μm (n=20) × 11–18–22 μm (n=20). Clads 128–191–244 (n=20) × 11–18–26 (n=20) μm. All samples pooled: rhabds 91–228–362 μm (n=56) × 11–19–30 μm (n=56). Clads 104–176–264 (n=123) × 10–19–31 (n=123) μm.

Microstrongyles: smooth; thickness tapers at the ends but most end in rounded and blunt tips. The doubly-bent form common in *P. cortius* is present, but the most common form is smoothly curved or with weak double bends. Some are more severely tapered and approach oxeas, with a few having oxeote tips. Most smaller spicules have a prominent centrotylote ball, with intermediate-sized spicules having a more subtle central swelling, and the largest spicules lacking a central swelling. Highly variable in length, but the distribution is continuous without clear size classes. When all samples are pooled, the distribution is weakly trimodal, with modes at about 40, 70, and 110 μm. Holotype: 35–66–103 (n=51) × 2–3–5 (n=23) μm. All samples pooled: 29–72–121 (n=259) × 1–3–7 (n=203) μm.

Oxyspherasters: smooth, with a round centrum and many (∼10–20) conical, sharply pointed rays. Present in all samples, but not abundant. Total diameter, holotype 9–12–14 μm (n=6); all samples pooled 8–14–27 μm (n=21).

Spheres: simple spheres are usually present in low numbers; few were measured, but they were 5–15 μm in diameter.

#### Distribution and habitat

Known only from the shallow waters of the western portion of Anacapa Island, Southern California. Exact collection depths not recorded, but all were between 3–12 m.

#### Remarks

Half of the samples of *Penares* collected at Anacapa Island were found to have small spicules and very distinct genotypes at the 28S locus (4% divergence to *P. cortius*, Fst = 0.99). The primers used to amplify the cox1 locus in other species failed in all of these samples, despite working in all other freshly collected *Penares*, likely because these samples have species-specific mismatches in the primer regions. Using additional custom primers, I was able to obtain a very small (84 bp) sequence from cox1 from two of these samples. In this small fragment, there are two unique genetic differences compared to other *Penares* (2% divergence to *P. cortius*, Fst = 1). *P. cortius* was also collected from Anacapa Island — in one case, on the same dive as *P. anyapax* sp. nov. — so differentiation at two unlinked loci supports the existence of reproductive isolation in nature (Jennings 1917; Rannala & Yang 2020). The samples designated here as *P. anyapax* sp. nov. are the sister species to *P. cortius*, based on samples with sequence data available.

*P. anyapax* sp. nov. is distinguished from *P. cortius* and *P. orientalis* comb. nov. by having shorter oxeas and microstrongyles, and from *P. saccharis* by having asters and strongyles instead of microxeas (the asters were found in all samples of *P. anyapax* sp. nov., but they were uncommon and often required considerable searching). Differentiation from *P. foxi* sp. nov. is more difficult. *P. anyapax* sp. nov. has more prominent bends and centrotylote swelling on the microstrongyles than *P. foxi* sp. nov., and the asters that are uncommon in *P. anyapax* sp. nov. and abundant in *P. foxi* sp. nov. The maximum lengths of the microstrongyles is also different: despite measuring 259 spicules, and searching for long examples, the longest microstrongyle seen in *P. anyapax* was 121 μm. Ten percent of the strongyles in *P. foxi* were longer than this, with a maximum of 147 μm. Additionally, the dichotriaene clad lengths are shorter (168–191 μm) in *P. anyapax* than *P. foxi* (224 μm). However, with only one sample of *P. foxi* sp. nov., variation in that species is uncharacterized, so it is highly recommended that genotyping be included in future discrimination of these species (they have 4% divergence at both the 28S and cox1 loci).

Outside of the temperate Northeast Pacific, no other *Penares* are likely to be conspecific with *P. foxi* sp. nov. based on biogeographic considerations, and all other species from the North Pacific are easily excluded based on spicular differences. The closest species are described from the Galapagos Islands; *P. angeli* Sim-Smith *et al*., 2021 and *P. apicospinatus* Desqueyroux-Faúndez & van Soest, 1997 have orthotriaenes instead of dichotriaenes, and *P. scabiosus* Desqueyroux-Faúndez & van Soest, 1997 and *P. foliaformis* Wilson, 1904 have much larger oxeas and dichotriaenes. Several additional species from Korea and Japan are also differentiated by having much larger or smaller spicules, as detailed in the comparative table in Sim-Smith and Kelly (2019).

### Penares orientalis comb. nov

Figure 29

**Figure 29.**
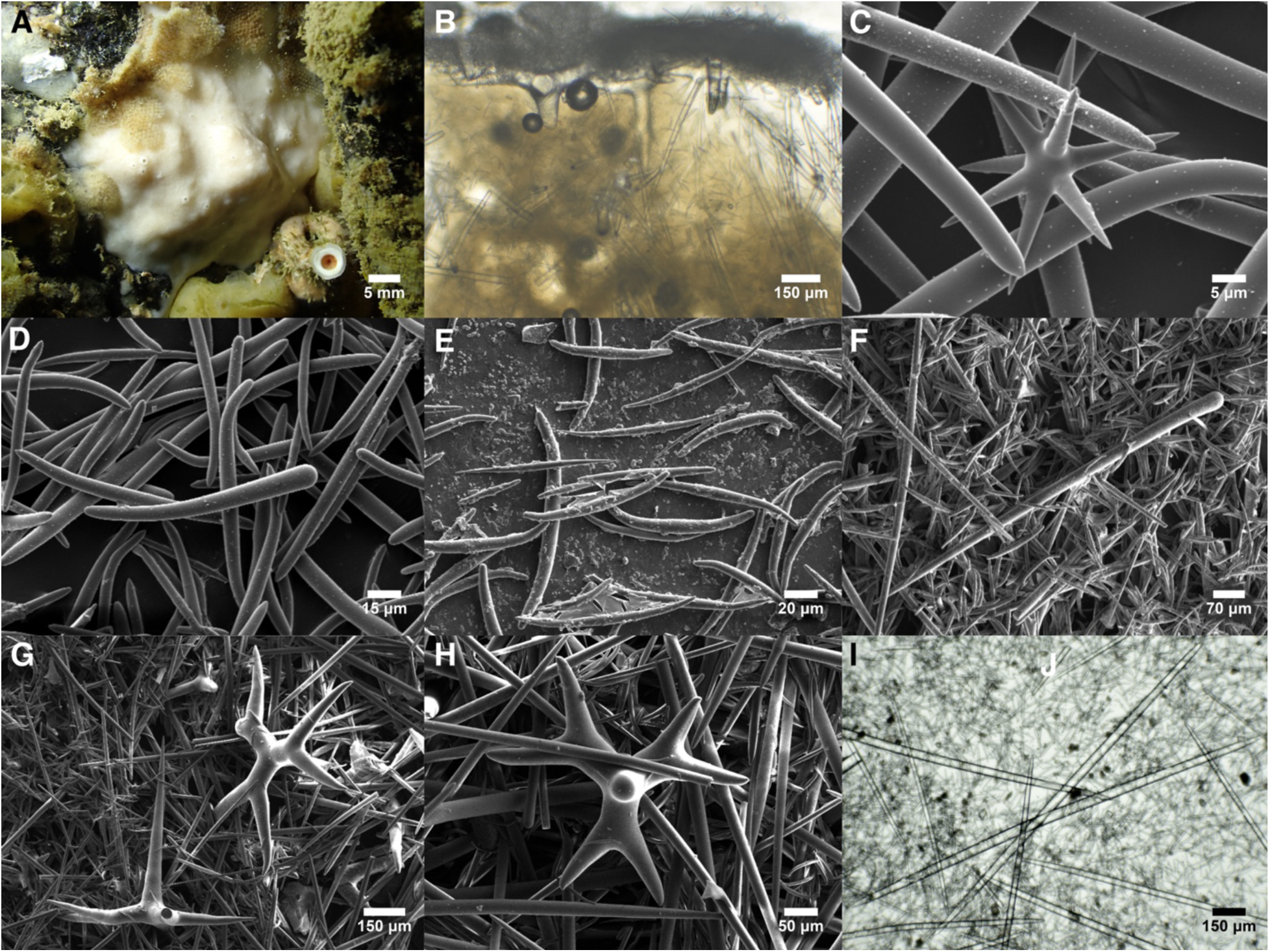
Penares orientalis. A: TLT1616 in situ. B: Perpendicular section at sponge surface showing a surface crust of tangential microstrongyles supported by dichotriaenes, with oxeas and microstrongyles confused in the interior; from TLT1616. C: Aster from CASIZ 302613. D: Club-shaped microsclere, with microstrongyles behind it, from CASIZ 302613. E: Microstrongyle variation from TLT1616, including some oxeote forms. F: Subtylostyle from TLT1616. G-H: Dichotriaenes from TLT1616. I: Oxeas from CASIZ 233435

#### Synonyms

*Penares* cortius ssp. orientalis Koltun, 1966

#### Material examined

TLT1616, Bird Island, Juneau, Alaska, (58.49035, -134.85168), 34 m, 28-Jun-2025; CASIZ 302613, Silver Bay, Sitka, Alaska, (57.02067, -135.16933), 24 m, 2-Sep-2010; RBCM 018-00912-001, SGaan Kinghlas-Bowie Seamount, British Columbia, (53.30020, -135.65290), to 67 m, 12-Jul-2018; CASIZ 233435, Offshore of Newport, Oregon, (44.64333, -124.43667), 100 m, 16-Apr-1962.

#### Morphology

Cushion-shaped to irregularly massive with an uneven thickness; oscula are about 0.5 mm in diameter, round with slightly raised rims, and scattered across the surface, especially at elevated points. One sponge collected at 34 m was white inside and outside, and the colors remained after preservation, but previously collected vouchers had a gray surface and white interior. All samples examined are fragments of larger sponges, but these were up to 5 cm in horizontal extent and up to 2 cm thick. Surface is smooth and leather-like.

#### Skeleton

The ectosomal skeleton is densely populated with tangential microstrongyles. The ectosome is supported by the cladomes of the dichotriaenes, with the rhabds pointing inward. The choanosomal skeleton includes tracts of oxeas 2–10 wide, parallel to the surface internally, but curving to meet the surface at the ectosome. The choanosome also contains scattered singular oxeas, asters, and microstrongyles in confusion. The skeleton did not differ markedly from *P. cortius*, though it seemed to have more microstrongyles scattered within the choanosome.

#### Spicules

Shown in figure 29 except where noted. See table 3 for a comparison of spicules among *Penares* species.

Oxeas: gently curved, with tips that vary from sharply pointed to conical and blunt. All samples pooled: 685–1420–2587 (n=126) × 12–27–53 (n=126) μm. Sample TLT1616 has relatively small oxeas, only 685–1178–1452 (n=39) × 12–22–31 (n=39) μm, while the other sample mean lengths vary from 1324–1726 μm. The description of the type sample states 1800–2400 × 33–50 μm (Koltun 1966).

Subtylostyles: with rounded heads and sharp tips. Rare in all samples except TLT1616, where they were less common than oxeas but still numerous. 824–919–1028 (n=10) × 16–32–60 (n=10) μm.

Dichotriaenes/plagiotriaenes: Most are well-formed dichotriaenes with long pointed rhabds and three forked clads, but a few simple triaenes were seen. Compared to the other *Penares* in the region, the clads appear to fork closer to the rhabd (the deuteroclads make up a larger proportion of the cladome). Strongylote rhabds and/or clads are sometimes present. The clads are sometimes unequal in length, and only the longest was measured. All samples pooled, rhabds 508–803–1221 μm (n=47) × 49–76–123 μm (n=47). Clads 154–413–695 (n=108) × 23–68–122 (n=108) μm. The type description has rhabds 670–920 μm × 93–120 μm, clads 268–335 μm (Koltun 1966).

Microstrongyles: smooth; thickness tapers at the tips but most end in rounded, blunt tips. As in *P. cortius*, the most common form is doubly-bent, but a large minority are smoothly curved or straight. Centrotylote swelling is weak or absent. A small minority are oxeas with sharply pointed tips. A larger minority are club-shaped styles, with one bend at the tapered end and the other end untapered (figure 29D). These club-shaped spicules were very rarely seen in other *Penares* from the region. The length distribution is continuous without obvious size classes. When all samples are pooled, the distribution is slightly trimodal, with modes at about 70, 120, and 150 μm. All samples pooled: 35–91–198 (n=350) × 3–6–12 (n=302) μm. Type description: 80–174 μm (Koltun 1966).

Oxyspherasters/oxyasters: small asters are smooth oxyspherasters, with a round centrum and many (∼10–20) conical, sharply pointed rays. Larger asters have a smaller centrum and fewer rays, until the largest asters have no centrum and as few as 4 rays. As they form a continuous series, small oxyspherasters and large oxyasters are pooled here. The distribution of total diameter is continuous but weakly multimodal, with a major mode at about 16 μm and minor modes at about 36 and 54 μm. Total diameter, all samples pooled 11–29–76 μm (n=167). Type description: 40–81 μm (Koltun 1966).

Spheres: simple glass spheres are present in low numbers; a few were measured at 10–70 μm in diameter. Not shown in figure 29; see figure 28I for example.

#### Distribution and habitat

The type sample was collected near Okusiri Island, Hokkaidō, Japan, with the depth stated only as "shallow waters". The samples examined here were collected from Southeast Alaska, British Columbia, and Oregon, from 24–100 m depth. An additional sample from Cordell Bank, Northern California is likely this species (YPM IZ 084200). This sample was not examined here, but photos of its spicules are available in GBIF. Too few spicules are imaged to make a confident identification, but the few dichotriaenes visualized appear to be too large relative to the microstrongules to be *P. cortius*. I therefore think it is likely that this species is found from Northern Japan to Northern California, from 24–135 m depth. Further work is needed to determine if the range of this species overlaps with *P. cortius*.

#### Remarks

*Penares cortius* was previously described as occurring as far north as Alaska (Austin 1985; Stone *et al*. 2011), but the samples I examined from Oregon, British Columbia, and Alaska were genetically and morphologically distinct compared to samples from Central and Southern California. Indeed, these samples were not sister groups in the phylogeny, with *P. cortius* from California more closely related to *P. anyapax* than to Northern samples.

In his work on the Tetractinellida from the Northwest Pacific, Koltun (1966) described a sponge as a new subspecies of *P. cortius*: *P. cortius orientalis*. The morphological description of this sample is a good fit to those determined here to be a different species than *P. cortius*, so I raise this subspecies to species status. Interestingly, I know of no *Penares* records between Northern Japan and the Gulf of Alaska. If the species ranges across this entire span, it would be expected to occur in the Aleutians, where it has not been reported (Stone *et al*. 2011). It therefore seems likely that the Japanese and North American populations are separate species, but as I am not able to differentiate them here, this remains as a question for future work. It also seems possible that there is additional diversity within this species concept in North America. Sample TLT1616 had short oxeas, more numerous styles, and few club-shaped microstrongyles compared to the other samples examined. It is also slightly differentiated at both sequenced loci (0.8% divergence at 28S, 0.4% divergence at cox1). As these genotypes are more closely related to each other than any other groups, and the morphological differences are slight, they are easily accommodated in a single species concept, pending further data.

*P. orientalis* comb. nov. is distinguished from the other species in the region by having larger spicules of all types. The most diagnostic difference is the presence of large oxyasters, which don’t occur in any other species in the region. *P. orientalis* comb. nov. also has a distinctive morph of club-shaped microstrongyle, which was present only very rarely in any other *Penares* from the region. The dichotriaenes are also subtly different in shape, with clads that fork closer to the rhabd (the deuteroclads make up a larger proportion of the cladome).

### Penares foxi sp. nov

Figure 30

**Figure 30.**
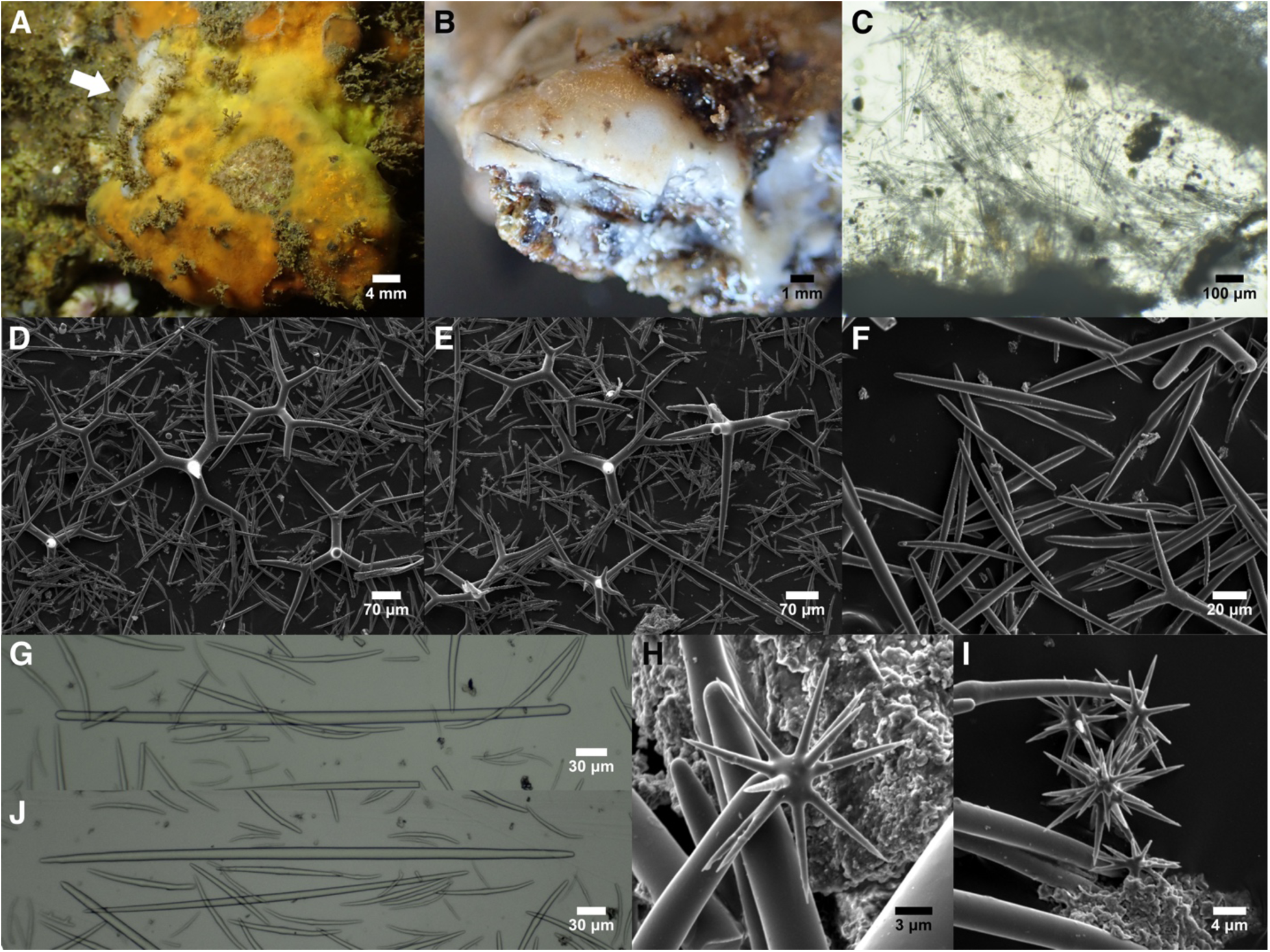
Penares foxi. A: In situ photograph; the small white sponge indicated by the arrow is the holotype, which was collected incidentally when collecting the more prominent *Hymedesmia*. B: Preserved sponge after a portion had been removed for analysis. C: Perpendicular section showing a surface crust of microstrongyles supported by dichotriaenes and oxeas in the choanosome. Dark areas at the base are where the sponge met the substrate. D-E: Dichotriaenes. F: Microstrongyles. G: Strongylote megasclere. H-I: Asters. J: Oxeote megasclere. All photos from holotype.

#### Material examined

Holotype, TLT1455B, Garbage Cove, Anacapa Isl., California, (34.01706, - 119.36391), 2-10 m, 28-Aug-2024.

#### Etymology

Named in memory of M. M. Fox (1932-1954), Coast Guard Engineman Petty Officer, 2nd Class. This species was found at Garbage Cove, which is so named because the Coast Guard formerly bulldozed garbage into a naturally concealed area at the site. Engineman Petty Officer Fox was bulldozing garbage into the site when the tractor he was operating went over the cliff. A memorial at the site reads "Here is where he fell. This man was talking when he should have been listening".

#### Morphology

The only known sample was found as a small encrustation, approximately 17 × 9 mm in horizontal extent and 2 mm thick. Several single, circular oscula are present, with slightly raised rims, about 0.5 mm in diameter. The sponge is entirely white, both alive and preserved.

Brownish discoloration on the surface appears to be caused by the presence of ∼1 mm long filamentous algae growing upright across much of the surface.

#### Skeleton

The ectosomal skeleton is densely populated with tangential microstrongyles. This layer is supported by the cladomes of the dichotriaenes, with the rhabds pointing inward. The base of the sponge, where it meets the substrate, is densely spiculated with both oxeas and microstrongyles. The oxeas rise from the base in disorganized tracts towards the surface, and there are also many oxeas in confusion. The asters are found in the choanosome.

#### Spicules

Shown in figure 30 except where noted. See table 3 for a comparison of spicules among *Penares* species.

Oxeas/strongyles/strongyloxeas/styles: the primary choanosomal spicules are diactinal, forming a continuous series from oxeas with sharply pointed tips, to strongyles with both tips blunt and rounded, or styles with one end untapered and the other end pointed. Some strongyles were faintly subtylote at the ends. 353–577–796 (n=51) × 4–11–18 (n=51) μm.

Dichotriaenes/plagiotriaenes: Most are well-formed dichotriaenes, but a few have one unforked clad, and one plagiotriaene was seen. A minority have strongylote rhabds or one or more strongylote clads. The clads are long in comparison to the rhabd, which means that few spicules are positioned in a way that allows for measurement of rhabd length. Rhabds 139–225–297 μm × 14–21–26 μm (n=8). Clads 124–224–342 × 10–22–33 (n=79) μm.

Microstrongyles: smooth; thickness tapers towards rounded and blunt ends. Spicules are straight or gently curved, though a minority have two slight bends, reminiscent of the doubly-bent form common in *P. cortius*. Most have a weak centrotylote swelling. The length distribution is continuous, but weakly multimodal, with modes at about 40, 65, and 130 μm. 30–71–147 (n=140) × 2–4–7 (n=140) μm.

Oxyspherasters: smooth, with a round centrum and many (∼10–20) conical, sharply pointed rays. Common. Total diameter 11–15–20 μm (n=27).

Spheres: simple glass spheres are present in low numbers; one was measured at 17 μm in diameter. Not shown in figure 30; see figure 28I.

#### Distribution and habitat

The only known sample was collected from a sea cave at Anacapa Island, Southern California. The cave is only partially submerged, with an entrance extending from 12 m above sea level to about 6 m below. The exact depth of the sample was not recorded, but it was collected between 2–6 m, on a small rock outcropping, where it was abutting a *Hymedesmia* sp. *Penares* sp. were abundant throughout the cave, including much larger samples, but only the one sample was collected, so it is uncertain if they were all *P. foxi* sp. nov.

#### Remarks

The spicules of this species are very similar to *P. anyapax* sp. nov., but genetic data from cox1 and 28S indicate that the species is highly distinct, and that it is the outgroup to the clade of *P. cortius, P. anyapax* sp. nov., and *P. orientalis* comb. nov. (figure 3). Despite having only one sample, this phylogenetic position clearly illustrates that a new name is needed.

*P. foxi* sp. nov. is distinguished from *P. cortius* and *P. orientalis* comb. nov. by having shorter oxeas and microstrongyles, and from *P. saccharis* by having asters and strongyles instead of microxeas. Differentiation from *P. anyapax* sp. nov. is more difficult based on spicules. The maximum length of the microstrongyles is longer in *P. foxi* sp. nov.: despite measuring 259 spicules, and searching for long examples, the longest microstrongyle seen in *P. anyapax* was 121 μm. Ten percent of the strongyles in *P. foxi* were longer than this, with a maximum of 147 μm. Additionally, the dichotriaene clad lengths are shorter (168–191 μm) in *P. anyapax* than *P. foxi* sp. nov. (224 μm). Indeed, the clads of *P. foxi* sp. nov. were the only *Penares* in the region to be as long as the rhabds. I suspect that many of the rhabds are actually shorter than the clads in *P. foxi* sp. nov., because nearly all of the dichotriaenes in this species were imaged with the rhabd facing up, probably because the long clads and short rhabds make this position more stable. Additional minor differences between these species include that *P. anyapax* sp. nov. have more prominent bends and centrotylote swelling on the microstrongyles, and that asters are abundant in *P. foxi* sp. nov. and uncommon in *P. anyapax* sp. nov. These differences are all slight, and we cannot characterize variation in *P. foxi* sp. nov. until more samples are collected. As such, it is highly recommended that genotyping be included in future discrimination of these species. They are not sister species, and have 4% divergence at both the 28S and cox1 loci.

Outside of the temperate Northeast Pacific, no other *Penares* are likely to be conspecific with *P. foxi* sp. nov. based on biogeographic considerations, and all other species from the North Pacific are easily excluded based on spicular differences. The closest species are described from the Galapagos Islands; *P. angeli* Sim-Smith *et al*., 2021 and *P. apicospinatus* Desqueyroux-Faúndez & van Soest, 1997 have orthotriaenes instead of dichotriaenes, and *P. scabiosus* Desqueyroux-Faúndez & van Soest, 1997 and *P. foliaformis* Wilson, 1904 have much larger oxeas and dichotriaenes. Several additional species from Korea and Japan are also differentiated by having much larger or small spicules, as detailed in the comparative table in Sim-Smith and Kelly (2019).

### *Penares saccharis* (de Laubenfels, 1930)

Figure 31

**Figure 31.**
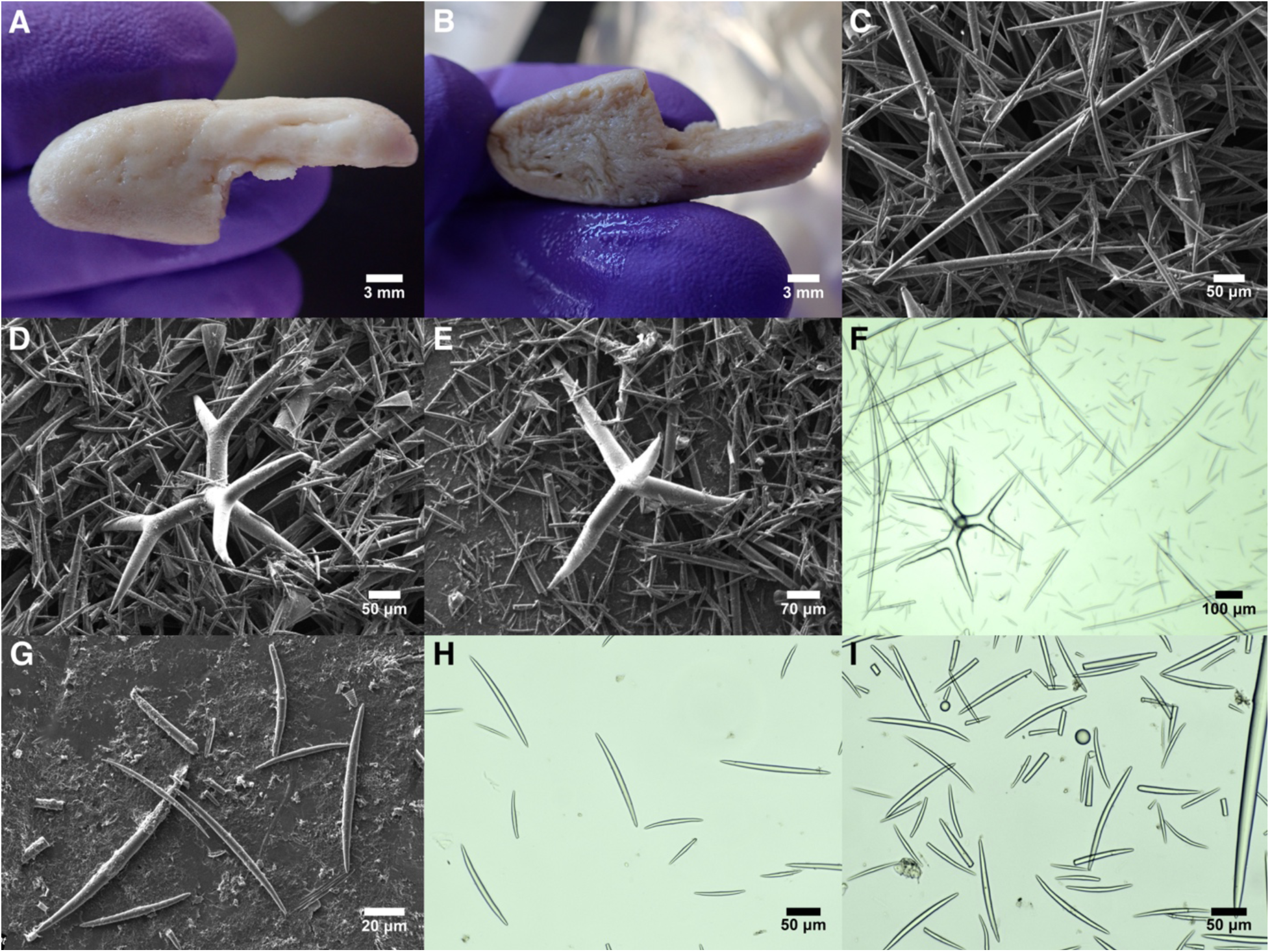
Penares saccharis. A-B: Preserved holotype. C: Oxea. D: Dichotriaene. E: Plagiotriaene. F: Dichotriaene, oxeas, and microxes imaged from a slide prepared from the holotype by de Laubenfels. G: Microxeas from the holotype, showing oxeote form and one strongylote example. H-I: Microxeas from the sequenced sample, CASIZ 450272, showing many oxeote forms but also styles and strongyles; spheres also visible in I. A-G are from the holotype.

#### Synonyms

*Papyrula saccharis* (de Laubenfels 1930, 1932)

#### Material examined

Holotype, USNM 21476, and paratype, USNM 21477, both collected at Point Pinos, Pacific Grove, Monterey, California, intertidal, 27-Jun-1926. Other sample: CASIZ 450272, Carmel Intertidal, Monterey, California, intertidal, 18-Jun-1947. Comparative material: USNM 37916, Gardner Island, Galapagos Islands, 22 m, Feb-1976.

#### Morphology

The holotype is the largest of the three samples, at 32 × 12 mm across and 9 mm thick, though it appears to have been sliced from a larger mass. Both of the other samples are also less than a centimeter thick. All three are white to off-white, smooth, with no visible oscula or other features, and all appear to be fragments of larger sponges. The original description states that samples can be up to 35 × 15 × 15 mm, samples were white alive, and that they contained pores 65–150 μm and round oscula up to 200 μm across (de Laubenfels 1932).

#### Skeleton

Not examined here, but the original species description does not differ markedly from the other *Penares* described above. The species was said to have a ectosomal skeleton of tangential microstrongyles supported by dichotriaenes, with a choanosomal skeleton containing tracts of oxeas up to 100 μm in diameter (de Laubenfels 1932).

#### Spicules

Shown in figure 31. See table 3 for a comparison of spicules among *Penares* species.

Oxeas/strongyloxeas/styles: most are oxeas, with tips varying from sharply pointed to more conical and blunt; styles are also seen. All are about the same size and are pooled here. Holotype: 496–788–1064 (n=26) × 10–18–27 (n=97) μm. All samples pooled: 469–724–1064 (n=97) × 8–19–32 (n=97) μm.

Dichotriaenes/plagiotriaenes: Dichotriaenes with long pointed rhabds and three forked clads are present in all samples, but simple triaenes with no forked clads (plagiotriaenes) are a significant minority, and intermediates with some forked clads are also seen. The clads are sometimes unequal in length, and only the longest was measured. Holotype: rhabds 296–331–386 μm (n=10) × 25–32–43 μm (n=10), clads 135–261–410 (n=28) × 22–32–48 (n=28) μm. All samples pooled: rhabds 165–291–386 μm (n=37) × 21–33–50 μm (n=37), clads 135–256–410 (n=88) × 16–31–48 (n=88) μm.

Microxeas: smooth, with only a small minority showing any centrotylote bulge. Many have sharply pointed tips, but many also have more rounded tips, and therefore resemble the strongyles seen in the other species of *Penares* from the region. Straight or slightly curved; occasionally with one bend, but only a few show any trace of the double bends common in *P. cortius*. Some samples also have a notable minority that are styles, with one oxeote end and one untapered rounded end; these are differently shaped compared to the more comma-shaped microstyles seen in *P. orientalis* comb. nov. Variable in length, but the distribution is continuous. When all spicules are pooled, the distribution is multimodal, with strong modes at about 80 and 150 μm, and a weak mode at 190 μm. Holotype: 43–89–181 (n=72) × 3–5–11 (n=72) μm; all spicules pooled: 43–105–205 (n=216) × 3–6–11 (n=181) μm.

Spheres: simple glass spheres are fairly common; diameter 11–20–26 (n=28) μm.

#### Distribution and habitat

Known with certainty only from the intertidal zone around the Monterey Peninsula, Central California. The type samples were found unattached, under stones near the low tide mark (de Laubenfels 1932), so it is unclear if they were intertidal sponges or washed ashore from greater depth. The only other sample I could assign to this name lists "Carmel intertidal" as the collection location, so it is from an unknown location on the southern side of the Monterey Peninsula. *P. saccharis* is not a common species in the region: I searched 14 intertidal areas across California, including the type locality and two other sites on the Monterey Peninsula, and was unable to find any *Penares*; there are also no pictures of intertidal *Penares* in California on the site iNaturalist.org. I saw only two subtidal *Penares* when surveying 14 subtidal sites around the Monterey Peninsula, and the only one that was collected proved to be *P. cortius*.

There are several scattered records of this species in other places around Southern California; I suspect these are misidentifications (see below). The species has also been reported from the Tropical Eastern Pacific, from Guerrero, Mexico and the Galapagos Islands, but as discussed below, I doubt the conspecificity of these samples as well.

#### Remarks

This species is differentiated from others in the region by its lack of asters and by having microxeas rather than microstrongyles. Both these characters are more difficult to assess than might be suspected. The microxeas are variable in shape, and many have fairly blunt tips. The strongyles in the other California species have blunter ends, but also taper somewhat, making them similar. Many *P. cortius* and *P. anyapax* sp. nov. also have few asters, and they are small and easy to miss. Nonetheless, careful examination revealed asters in every other *Penares* analyzed, and none in the holotype or paratype of *P. saccharis*. The type samples were definitely more oxeote than any sample assigned to the other species. Using these characteristics, I was able to associate one other sample with this name: a sample from the intertidal near Carmel, California, collected in 1947 (CASIZ 450272). This is fortuitous, because I have been unable to obtain DNA sequence from any of de Laubenfel’s type samples, and this species is no exception: even very small amplicons (100 bp) failed. I was, however, able to use these "mini-barcode" primers to generate 83 bp of cox1 sequence from this additional sample from 1947. Evolutionary relationships inferred from such a small fragment are admittedly preliminary. With this in mind, it is interesting that this species appears to be distantly related to the other *Penares* in the region (cox1 figure S1). Instead, it is sister to *P. candidatus*, which then forms a clade with *P. sphaera* and *P. alatus*. All of these species lack asters. The absence of asters was used to define the genus *Papyrula*, to which *P. saccharis*, *P. candidatus,* and *P. sphaera* were previously allocated. There are additional *Penares* that lack asters, but to my knowledge, none have DNA data available. As a result, it is possible that *Papyrula* will be revived for this clade in the future.

There are several scattered records of this species in other places around Southern California. I examined one of these (USNM 31321) and it has asters and spicules closer to microstrongyles than microxeas, so I reassigned it to *P. cortius*. I suspect the other records are also misidentifications caused by the rarity of asters and tapering nature of the microstrongyles in many *P. cortius*. *P. saccharis* has also been reported from the Tropical Eastern Pacific, from Guerrero, Mexico (Gómez *et al*. 2002) and the Galapagos Islands (Desqueyroux-Faúndez & van Soest 1997). I think these records are likely to be a different species on biogeographic grounds. The morphological data published for these samples is admittedly very similar to *P. saccharis*, however. I reexamined the sample from the Galapagos Islands (USNM 37916), and found that the oxeas were considerably longer than the three Californian *P. saccharis* (Galapagos sample: 759–1005–1319 (n=35); California samples: 469–724–1064 (n=97), with means of 696, 703, 788 μm). Furthermore, the Galapagos sample had two distinct classes of microxeas (my measurements: 36–64–87 (n=27) and 115–156–199 (n=39) μm) rather than a continuous distribution; this was also noted by the authors who described the sample (Desqueyroux-Faúndez & van Soest 1997). The dimensions of these and all other spicules in the Galapagos sample are included in the supplementary table S2. Genotyping of this Galapagos sample was attempted with a variety of primer sets, but was not successful. Future work is needed for confirmation, but I suggest reassigning this sample to *Penares* cf. *saccharis*.

### Family Theneidae Gray, 1872

#### Definition

Astrophorina with long-shafted triaenes (sometimes lost) in combination with diverse categories of streptasters: spirasters, metasters and plesiasters (sometimes with annulate actines). From Cárdenas *et al*. (2011).

### Genus *Thenea* Gray, 1867

#### Definition

Theneidae with tetraxonic megascleres and without cladotyles. From Cárdenas & Rapp (2012).

### Thenea diastra sp. nov

Figure 32

**Figure 32.**
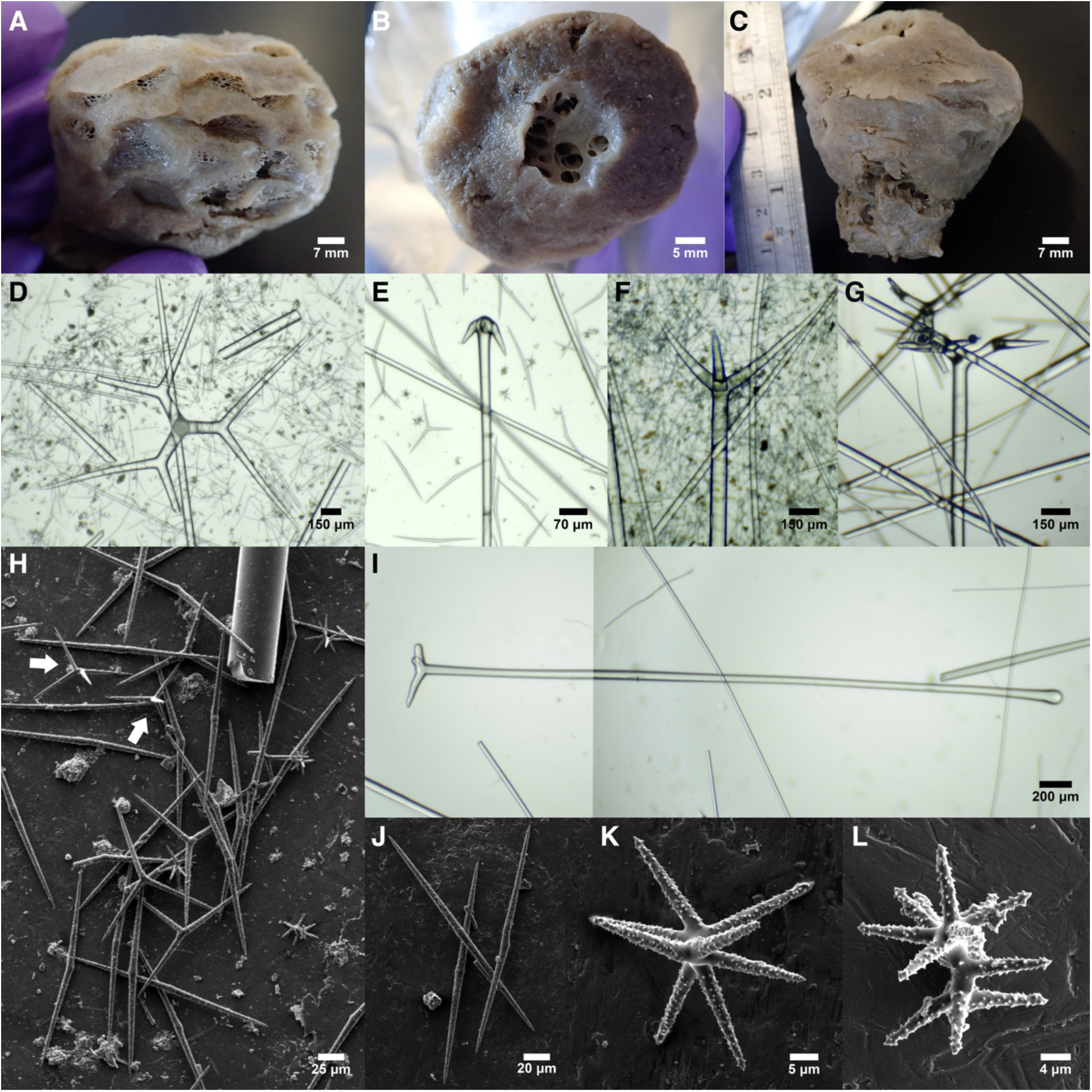
Thenea diastra. A-C: The preserved holotype viewed from the side with many pore areas (A), the top (B), and the side with fewer pore areas (C). D: Dichotriaene with long deuteroclads. E: Anatriaene. F: Protriaene. G: Dichotriaene with relatively short clads. H: Aster variation. The most common asters are large two-rayed plesiasters, but some large plesiasters with three rays are also visible, center. White arrows indicate examples of the slightly smaller class of four and five-rayed plesiasters. Smaller metasters are also visible. I: Example of an aberrant spicule, in this case a triaene with short clads; composite of two photos. Note the tylote rhabd, which was common in all types of triaenes. J: Large two-rayed plesiasters. K: Large metaster. L: Small metaster/spiratster. All photos from holotype.

#### Material examined

Holotype, RBCM 020-00056-001, Baby Bare Seamount, British Columbia, (47.70633, -127.78550), 2598 m, 8-Aug-1995.

#### Etymology

Named for the two-rayed asters that predominate among the large plesiasters.

#### Morphology

Shaped like an inverted cone, with a flat top and tapering sides. Top is 5 cm across, flat but tilted and off-center compared to the vertical axis of the sponge. A 1.8 × 1.6 cm oscular opening is present in the center of the flat top, with multiple internal channels leading into it.

Viewed from the side, the sponge is 6 cm across at the top, tapering to a short stalk 2.5 cm across. Many pore areas are present on the sides of the sponge, mainly on the side opposite to the sloping top, where hemispherical pore areas ∼ 10 × 4 mm across cover most of the sponge surface. Visibly hispid, but much less so than many other *Thenea*, with only a few long protruding oxeas scattered about on the surface. Beige in ethanol; color alive unknown.

#### Skeleton

The ectosomal skeleton is densely populated with large plesiasters (as most have only two rays, these are essentially microxeas) which are tangential to the surface. The ectosome is supported by dichotriaenes, which are oriented with their cladomes just below the ectosome and rhabds pointing inwards. Long oxeas are also oriented individually, in a roughly radial fashion, alongside the long rhabds of the dichotriaenes. The large two-rayed plesiasters are also very dense in the choanosome, in contrast to the megascleres, which are relatively sparse.

#### Spicules

Shown in figure 32 except where noted.

Oxeas: very long thin oxeas are abundant. The tips taper at both ends, but often to rounded, bunt tips. 5660–9797–11690 × 51–65–74 μm (n=11).

Tylostrongyles: two spicules were seen that had straight shafts and rounded, tylote ends; these were likely rare aberrations of another spicule type. 1990–2418–2847 × 47–59–71 μm (n=2). Not shown.

Dichotriaenes: The most common triaene. The protoclads diverge from the rhabds at an angle greater than 90 degrees, but the deuteroclads then flatten out to be orthogonal to the rhabd. The protoclads are invariably short, while the deuteroclads vary from short to very long. The rhabds and clads are long, thin, and frequently broken; those that aren’t broken frequently have subtylote endings. There was also a large number of aberrations seen, varying from orthomonaenes to spicules with one or more small numbs that appeared to be derived from triaene clads. Only one unbroken rhabd was seen, 7214 μm long; rhabd widths 47–77–96 μm (n=32). When measured from clad tip to rhabd, clads are 283–869–2300 × 31–64–138 (n=32) μm. Protoclads 128–206–358 (n=35) μm; deuteroclads 119–680–2056 (n=37) μm.

Protriaenes/plagiotriaenes: Protriaenes with curved clads are common, but these intergrade with plagiotriaenes that have a smaller angle with the rhabd. A large minority have one or more clads reduced to rounded nubs. All unbroken rhabds had subtylote endings, whereas clads that weren’t reduced to nubs generally had sharply pointed tips. Rhabds 3123–4499–6056 μm (n=5) × 20–66–96 μm (n=19). Longest clads 143–725–1094 × 22–54–76 (n=18) μm. Clad:rhabd angle 23–46°–70° (n=18).

Anatriaenes: only two examples were seen despite repeated spicule preps from multiple regions of the sponge. The only one with an unbroken rhabd ended in a subtylote ball rather than a sharp tip. Rhabd 5145 × 18–23–28 μm (n=2). Clads 60–85–110 × 17–24–31 μm (n=2). Clad:rhabd angle 40°–45°–49° (n=2).

Large plesiasters: the largest size class of plesiasters is mostly comprised of asters with only two rays, which therefore look like bent, microspined oxeas. The presence of three-rayed asters as a minority among them belies their true form. A few are also seen as one-rayed asters, which appear as acanthose subtylostyles. Ray lengths 67–90–122 μm, total diameter 116–170–219 μm (n=40).

Small plesiasters: A less common and slightly smaller class of plesiasters is seen, with 3–5 rays. These grade into the large plesiasters. Ray lengths 39–51–66 μm, total diameter 68–94–119 μm (n=10).

Metasters/spirasters: The smallest class of asters are primarily metasters, but the smallest asters among them tend towards spirasters. these have 6–12 rays, with smaller asters generally having more rays. Ray lengths 10–19–28 (n=29) μm, total diameter 28–40–61 (n=40) μm.

#### Distribution and habitat

The only known sample was collected at 2598 m at Baby Bare Seamount. Detailed collection data is not linked to the voucher, but it is noted as coming from "Verena Tunnicliffe’s collection", and it may have been collected at a cold seep.

#### Remarks

There are 38 described species of *Thenea* worldwide (de Voogd, *et al*. 2025). In the North Pacific, a remarkable nine species are described from warm temperate regions in Japan (Sollas 1886; Thiele 1898), though most have very little published data available and require revision. The only other North Pacific species, *T. wilsoni* van Soest *et al*., 2020, is described from the Mexican Tropical Pacific (Wilson 1904). No species are known to reside in the Temperate Northeast Pacific, nor did Koltun (1966) find any in the cold Northwest Pacific. However, recent deep-water visual surveys performed with ROVs in the region have reported thousands of records of apparent *Thenea*, from British Columbia to Southern California (GBIF). These *Thenea* are either reported as *Thenea* sp. or *T. muricata* (Bowerbank 1858), a species from the North Atlantic that is highly unlikely to occur in the region. Despite these thousands of visual observations, I know of only four collected vouchers from the region, none of which have any published analysis. I examined one of these, and found it to be markedly different than any described species. Most notably, the predominance of two-rayed plesiasters differentiates this sponge from all previously described species. Two-rayed plesiasters are sometimes present in North Atlantic *T. muricata,* but only as a small minority (Cárdenas & Rapp 2012; Despujols 2021). They are similarly present in the Indian Ocean species *T. corallophila* Dendy & Burton, 1926, but again as a minority, and these are larger than those reported here. Additionally, nearly all described species have pore areas that are limited to the equatorial region of the sponge, usually as one continuous area, in contrast to the many pore areas seen in the new species, spread across the surface of the sponge.

Despite the sample having been preserved in formalin, I was able to amplify a 152 bp "mini-barcode" from the 28S locus for this species. Despite its very short length, this was sufficient to confidently place it in a clade with other *Thenea* (figure S2), and to show that it was highly divergent from North Atlantic species (e.g., 6% sequence divergence vs. *T. muricata*). The only sequence data from Pacific *Thenea* are from samples collected from the Clarion-Clipperton Zone in the Tropical Pacific (Lim *et al*. 2017). These samples, which were not identified to species, appear to be from two species based on genetic data alone. Five sequenced vouchers from that study are closer to Atlantic *Thenea* than to *T. diastra* sp. nov. (6% divergence), but one voucher (RC0828) differs from *T. diastra* sp. nov. by only a single base pair (1% divergence). Further comparisons of these samples is a high priority for future work, as is examining the three other known vouchers of *Thenea* from the Northeast Pacific, and collecting additional samples. Post-preservation images of two samples collected by the E/V Nautilus in California were shared with me by Alana Rivera, curatorial assistant at the Museum of Comparative Zoology, Harvard; these samples show no resemblance to the species described here.

## Conclusions

The sponges of the North Pacific have been the focus of surprisingly little research, compared to the sponges of the North Atlantic, Caribbean, or Southwest Pacific. One potential contributor to this phenomenon is that the sponges in the North Pacific lack the solid systematic foundations that all other research relies upon. Uncharacterized diversity can confront researchers with a confusing tangle to decipher before other work is possible. Here, I have attempted to untangle the diversity of the Astrophorina for the region, so that this diversity presents an opportunity, rather than an intimidating quagmire. Whenever possible, I have gathered data on many aspects of each species, from traditional taxonomic characters, to in situ morphology, to DNA sequences, so that future researchers will have multiple levers available when trying to understand the ecology, evolution, or applied potential of sponges in this group.

This is not the end of the systematic work on this group in this region, or even the beginning of the end, but perhaps it is the end of the beginning. Building upon the classic (de Laubenfels 1932; Lendenfeld 1910) and the more recent work (Lee *et al*. 2007; Stone *et al*. 2011) of others, I have brought our understanding of astrophorid sponges in the region into the 21st century, but in many ways, these results highlight how much remains to be deciphered. For example, extensive SCUBA-based collections identified many new shallow-water species, but these were mainly in Southern California. This could be because shallow water diversity in the Tetractinellida is higher in warm temperate regions than cold temperate regions (Cárdenas *et al*. 2011), but it is also true that the majority of recent SCUBA collection effort has focused on Southern California. Expanding collecting efforts in other regions is likely to net additional species and better establish the ranges of the new species described here. I also discovered multiple new species inhabiting deep water by reexamining sponges vouchered in museum collections. It should be emphasized that finding new deep-water species was not my focus: the vouchers I acquired on loan were primarily intended to understand the status and distribution of species previously thought to be in the region, establish their diagnosability, and attempt to place them in molecular phylogenies. A more comprehensive look at museum collections, empowered by the data published here, will undoubtedly net additional species that have already been collected. Additional collecting in deep water is clearly needed as well. Some recent surveys in the region have been conducted primarily as ROV video surveys, but the results reported here makes it clear that there are more species present in the region than previously thought, some are likely to be undiagnosable from field images alone, and there are surely many new species that will only be identified with vouchered samples. I examined one *Thenea* sample and described one new species — there are very likely more species in the region requiring description. Our understanding of *Geodia* is also far from complete, with multiple species (*G. ovis, G. mesotriaenella, G. bicolor*) having no DNA data and very limited understanding of their morphology, range, or abundance. Despite the necessity for more data, I wish to highlight how replicable the *Geodia* data reported by Lendenfeld (1910) was, in contrast to the published claim to the contrary by de Laubenfels (1932). Though I propose that *G. breviana* a junior synonym of *G. mesotriaenella*, I support all of the other species’ concepts Lendenfeld proposed, and the quantitative data I collected on spicule dimensions was highly congruent to the numbers he reported over a century ago. Also, though I do not consider all of the Lendenfeld’s proposed subspecific variants to be useful taxonomic units, I do think that several should be investigated further as possible new species (i.e. *G. angulata* var. *megana* and *G. carolae* var. *megasterra*).

In addition to incomplete data on *Geodia* species, our understanding of *Poecillastra* diversity, though much improved, is far less certain than I would like. Excepting *P. alaskensis* sp. nov. in the far North, I found no evidence of multiple species being present, but there are hints from the mitochondrial DNA that genome-scale data would paint a different picture. I will also harbor continued concerns about whether *P. rickettsi* is truly the correct name for the widespread *Poecillastra* species in the region until samples with very long "coronal" oxeas, better matching the type description, can be found, collected, and characterized (including genotyping).

I wish to highlight that the Northeast Pacific temperate zone continues into Northern Mexico, and that Mexican waters are not included in this revision. The "Southern California Bight" marine ecoregion continues into Northern Baja, Mexico (Spalding *et al*. 2007), and many species included here are expected from this area as well. The warm-temperate Magdelenan and Cortezian marine ecoregion in Mexico include additional described species that have not been found in California (e.g. *Geodia californica* (Lendenfeld, 1910)), and these should be the focus of future revisions that build on this work. Additionally, working to revise sponge diversity in the Northeast Pacific serves to emphasize how valuable similar projects would be in the Northwest Pacific, where classical work is in similar need of updates that include genetic data and quantitative morphological data.

I hope the results presented here will motivate work on the ecology and evolution of the Astrophorina within this region and beyond. For example, I found that there were apparent "sponge grounds" of large *Geodia* at diving depths in Southern California. These seem likely to be both ecologically important and environmentally vulnerable, given their proximity to highly populated areas, but are entirely undescribed in any literature. I also present updated global phylogenies for the order, which include new possibilities for better aligning the taxonomy of the order with its evolutionary history, by, for example, resurrecting the genus *Papyrula*. Together with a recent revision of the Tetillidae (Turner 2026), the work presented here places the order Tetracinellida on a much more solid footing, but in the final analysis, only shows up how much more remains to be done.

## Key to the Astrophorina of the temperate Pacific coasts of the United States and Canada

1A: Sponges where all the triaenes are long-shafted: one ray (the rhabd) is much longer than all others (the clads): *Geodia, Stelletta, Thenea, Penares, Ancorina*. Clads may branched or unbranched: 7

1B: Sponges where most of the triaenes are short-shafted: no one clad is obviously a rhabd. *Poecillastra, Vulcanella, Dercitus*. Note that *Vulcanella* may include some long-shafted triaenes as well: 2

2A: No oxeas. Sponges contain only calthrops and sandiasters (*Dercitus*): 3

2B: Megascleres include oxeas; all but one species also has acanthose microxeas (*Poecillastra, Vulcanella*): 4

3A: Calthrop rays average less than 100 μm in length: *Dercitus (Stoeba) syrmatitus*. Known from the Southern California intertidal.

3B: Calthrop rays average more than 200 μm in length: *Dercitus (Stobea) giveni.* Known from shallow water in Southern California.

4A: Lacks acanthose microxeas: *Vulcanella explorata* sp. nov. Known from deep water in British Columbia.

4B: Has acanthose microxeas: 5

5A: Largest asters are more than 60 μm in diameter: *Poecillastra alaskensis* sp. nov. Known from Alaska and British Columbia.

5B: Largest asters are less than 60 μm in diameter: 6

6A: Acanthose microxeas in two size categories; triaenes include calthrops, long-shafted orthotriaenes, and dichotriaenes: *Vulcanella rupta* sp. nov. Sponges are subspherical to cup-shaped and known from deep waters in British Columbia.

6B: Acanthose microxeas in one size category; all triaenes are short-shafted and calthrops-like: *Poecillastra rickettsi*. Sponges vary from small and globular to large horizontal vases and possibly plate or fan shapes. Ranges from British Columbia to California, and possibly farther in both directions.

7A: Has sterrasters (*Geodia*) or aspidasters (sterraster-like but flattened, see Lehnert *et al*. (2006); *Erylus*): 23

7B: Lacks sterrasters and lacks aspidasters: 8

8A: Triaenes include dichotriaenes that have very long (>1000 μm) deuteroclads (figure 32D); the most abundant asters are acanthose plesiasters that mostly have only two rays and resemble bent oxeas, but some three-rayed examples reveal them to be asters; sponge roughly spherical / conical with an apical oscule: *Thenea diastra* sp. nov. Known from deep water in British Columbia.

8B: Lacking these features: 9

9A: Has sandiasters among the microscleres (see figure 4J in Lehnert and Stone, 2014): *Ancorina buldira*. Known from the Aleutian Islands.

9B: Lacks elongated sandiasters; all asters are roughly spherical euasters and/or anthasters that look like tiny, irregular sand grains in the light microscope (asters can also be rare or absent): 10

10A: The most abundant small spicules are smooth (non-acanthose) microstrongyles (figure 27F) or microxeas (figure 31H) which form a surface crust (Genus *Penares*): 11

10B: Lacking abundant microstrongyles/microxeas (Genus *Stelletta*): 15

11A: Lacks asters; has primarily microxeas rather than strongyles: *Penares saccharis*. Known from Central California.

11B: Has asters, though they may be rare; has primarily microstrongyles rather than microxeas: 12

12A: Mean aster diameter more than 20 μm, including some over 40 μm: *Penares orientalis* comb. nov. Known from Japan to Oregon, probably also ranges into Northern California and possibly farther South.

12B: Mean oxyaster diameter less than 20 μm, with maximum diameters less than 30 μm: 13

13A: Microstrongyle mean length over 85 μm, with maximum values over 150 μm; oxea mean length over 650 μm: *Penares cortius*. Known from California but may range farther North.

13B: Microstrongyle mean length less than 85 μm, with maximum values less than 150 μm; oxea mean length less than 650 μm: 14

14A: Mean dichotriaene clad length less than 200 μm; maximum microstrongyle length less than 130 μm: *Penares anyapax* sp. nov. Known from Anacapa Island, California.

14B: Mean dichotriaene clad length more than 200 μm; maximum microstrongyle length more than 130 μm: *Penares foxi* sp. nov. Known from Anacapa Island, California. (Note that this species is poorly known and genetic discrimination of *Penares anyapax* sp. nov. and *Penares foxi* sp. nov. is highly recommended.)

15A: Has anthasters (figure 12E; these look like tiny, irregular sand grains in the light microscope): 16

15B: No anthasters: 17

16A: Triaenes are primarily dichotriaenes; oxyasters are evenly acanthose: *Stelletta clarella.* Known from Southern Alaska to Southern California.

16B: Triaenes are primarily orthotriaenes/plagiotriaenes with recurved clads; oxyasters with spines concentrated at tips: *Stelletta anthastra.* Known from the Aleutian Islands.

17A: Oxea mean length more than 3500 μm; mean length of triaene rhabds over 2500 μm; only known from British Columbia and Alaska: 18

17B: Oxea mean length less than 3500 μm; mean length of triaene rhabds less than 2500 μm; only known from California: 19

18A: Triaene spicules are orthotriaenes/plagiotriaenes with recurved clads (clads sometimes reduced to nubs); asters consist only of acanthose oxyasters: *Stelletta makushina*. Known from the Aleutian Islands to British Columbia.

18B: Triaene spicules are dichotriaenes; asters include both small, unspined or lightly spined oxyasters and larger acanthose asters with blunt, conical rays: *Stelletta ovalae*. Known from Japan to British Columbia.

19A: Lacks dichotriaenes: 20 19B: Has dichotriaenes: 21

20A: Sponge surface is white (alive and preserved); maximum triaene rhabd lengths usually more than 2000 μm; acanthose asters are both oxyote and strongylote: *Stelletta estrella*. Known from Southern California.

20B: Sponge surface is gray (alive and preserved); maximum triaene rhabd lengths less than 2000 μm, acanthose asters are oxyote: *Stelletta cardenasi* sp. nov., in part. Known from Southern California. (Genetic confirmation is recommended when differentiating *Stelletta estrella* and *Stelletta cardenasi* sp. nov., as they are morphologically similar and variation may exceed what was found here)

21A: Mean dichotriaene rhabd length less than 1100 μm: 22

21B: Mean dichotriaene rhabd length more than 1100 μm; possesses small and large classes of unspined asters: *Stelletta nicolenya* sp. nov. Known from San Nicholas Island, Southern California.

22A: Orthotriaene/plagiotriaene rhabd mean length less than 700 μm; oxea mean length less than 1400 μm; possess small and large classes of unspined asters: *Stelletta limuwensis* sp. nov.

22B: Orthotriaene/plagiotriaene rhabd mean length more than 700 μm; oxea mean length more than 1400 μm; possess only a small size class of unspined asters: *Stelletta cardenasi* sp. nov. (in part). Known from Southern California.

23A: Has flattened aspidasters rather than spherical sterrasters; has centrotylote microrhabds: *Erylus aleuticus.* Known from the Aleutian Islands.

23B: Has spherical sterrasters and lacks centrotylote microrhabds (*Geodia*): 24

24A: Has plagiotriaenes: 25

24B: Lacks plagiotriaenes; has plagiomonaenes and/or plagiodiaenes, most of which are style-like with only small rounded nubs for clads: *Geodia carolae.* Known throughout the region and into the temperate Northwest Pacific.

25A: Lacks mesoprotriaenes (and/or derivatives like protriaenes, mesoprodiaenes, etc.); sterraster rosettes not warty; strongylasters with large, spherical centrum that makes them very distinct from oxyasters: 26

25B: Has mesoprotriaenes; sterraster rosettes warty; strongylasters lack dramatically inflated centrum and are difficult to separate from oxyasters: 28

26A: Sterrasters in two size classes, one class 110–115 μm in diameter, the other 165–203 μm in diameter: *Geodia starki*. Known from the Aleutian Islands.

26B: Sterrasters in one size class … 27

27A: Sterraster mean diameter less than 125 μm; oxyasters unspined: *Geodia angulata*. Known from California.

27B: Sterraster mean diameter more than 125 μm; oxyasters spined: *Geodia bicolor.* Known from California.

28A: Sterraster mean diameter less than 84 μm; epirhabds of mesotriaenes are shorter than clads: 29

28B: Sterraster mean diameter more than 84 μm; epirhabds of mesotriaenes are longer than or equal to the length of the clads: 30

29A: Plagiotriaene mean length less than 3700 μm: *Geodia mesotriaenella*. Known from British Columbia to Southern California.

29B: Plagiotriaene mean length more than 3700 μm: *Geodia ovis*. Known from Southern California.

30A: Plagiotriaene mean length less than 4150 μm: *Geodia agassizii*. Known from the Gulf of Alaska to Southern California.

30B: Plagiotriaene mean length more than 4150 μm: *Geodia mesotriaena.* Known from Southern California, but may range farther North.

## Acknowledgements

I am grateful for the help and support of many people in UCSB’s Marine Science Institute, especially Robert Miller, Clint Nelson, Christoph Pierre, and Christian Orsini. Steve Lonhart and The Monterey Bay National Marine Sanctuary were instrumental in facilitating collections in Central California. The Olympic Coast National Marine Sanctuary was instrumental in facilitating collections in Washington. The Natural History Museum of Los Angeles’ DISCO program facilitated collections in Los Angeles County. Allen Collins, Hugh MacIntosh, Christina Piotrowski, Kathy Omura, and Vanessa Delvanez graciously provided access to their collections at the Smithsonian National Museum of Natural History, the Royal British Columbia Museum, the California Academy of Sciences, the Natural History Museum of Los Angeles, and the Santa Barbara Natural History Museum, respectively. Thanks to Kristen Westfall (Fisheries and Oceans Canada) for the mitochondrial genome assembly of *P. rickettsi* sample RBCM 025-00144-001. Thanks to Lewis Sharpnack and Gareth Seward at the UCSB Earth Science Electron Microscopy and Microanalysis Lab for their assistance with SEM. Several of the California Academy of Sciences samples were collected by the Coral Reef Research Foundation under contract to the US National Cancer Institute. Finally, I thank the editor and reviewers for improving the manuscript.

## Funding Declaration

Financial support was provided by UCSB and by the National Aeronautics and Space Administration Biodiversity and Ecological Forecasting Program (Grant NNX14AR62A); the Bureau of Ocean Energy Management Environmental Studies Program (BOEM Agreement MC15AC00006); the National Oceanic and Atmospheric Administration (NOAA) in support of the Santa Barbara Channel Marine Biodiversity Observation Network; and the U.S. National Science Foundation (NSF) in support of the Santa Barbara Coastal Long Term Ecological Research program under Awards OCE-9982105, OCE-0620276, OCE-1232779, OCE-1831937, and the Tula Foundation. The funders had no role in study design, data collection and analysis, decision to publish, or preparation of the manuscript.

**Table 4.** Voucher and accession numbers of all samples. See table S1 for an expanded table with additional information.

| Species | Type status | Sample ID | Museum vouchers | Mitochondrial accession | Ribosomal accession |
| --- | --- | --- | --- | --- | --- |
| *Dercitus* (Stoebea) *giveni* | Holotype | n/a | SBMNH 659287 | n/a | n/a |
| *Dercitus* (Stoebea) *symmatitus* | Holotype | n/a | US 21438 | n/a | n/a |
| *Dercitus* (Stoebea) *symmatitus* | Paratype | n/a | US 21439 | n/a | n/a |
| *Geodia agassizii* | n/a | n/a | USNM 1478670 | TBD | TBD |
| *Geodia agassizii* | Syntype | n/a | USNM 8362 | n/a | n/a |
| *Geodia agassizii* | Syntype | n/a | USNM 8381 | TBD | n/a |
| *Geodia agassizii* | n/a | n/a | RBCM 019-00567-003 | TBD | n/a |
| *Geodia agassizii* | n/a | n/a | RBCM 007-00011-004 | TBD | n/a |
| *Geodia agassizii* | Syntype | n/a | USNM 8384 | TBD | n/a |
| *Geodia agassizii* | n/a | n/a | CASIZ 229525 | n/a | n/a |
| *Geodia agassizii* | n/a | n/a | CASIZ 233435 | n/a | n/a |
| *Geodia angulata* | n/a | TLT1175 | TBD | TBD | TBD |
| *Geodia angulata* | n/a | TLT1176 | TBD | n/a | n/a |
| *Geodia angulata* | n/a | TLT576 | TBD | TBD | TBD |
| *Geodia angulata* var. *megana* | Syntype | n/a | USNM 8403 | TBD | n/a |
| *Geodia angulata* var. *microana* | Holotype | n/a | USNM 8380 | TBD | n/a |
| *Geodia angulata* var. *orthotriaena* | Holotype | n/a | USNM 8402 | n/a | n/a |
| *Geodia bicolor* | Syntype | n/a | USNM 8398 | n/a | n/a |
| *Geodia bicolor* | Syntype | n/a | USNM 8399 | n/a | n/a |
| *Geodia breviana* | Holotype | n/a | USNM 8367 | n/a | n/a |
| *Geodia carolae* | n/a | n/a | RBCM 980-00257-005 | TBD | n/a |
| *Geodia carolae* | n/a | n/a | RBCM 980-00232-008 | n/a | n/a |
| *Geodia carolae* | n/a | n/a | RBCM 988-00002-094 | TBD | n/a |
| Geodia carolae var. carloae | Holotype | n/a | USNM 8391 | n/a | n/a |
| Geodia mesotriaena | n/a | TLT1444 | TBD | TBD | n/a |
| Geodia mesotriaena | n/a | TLT586 | TBD | TBD | TBD |
| Geodia mesotriaena | n/a | TLT589 | TBD | TBD | TBD |
| Geodia mesotriaena | n/a | n/a | SBMNH 693792 | n/a | n/a |
| Geodia mesotriaena var. megana | Holotype | n/a | USNM 8409 | TBD | n/a |
| Geodia mesotriaena var. microana | Holotype | n/a | USNM 8410 | TBD | n/a |
| Geodia mesotriaena var. pachana | Holotype | n/a | USNM 8571 | TBD | n/a |
| Geodia mesotriaenella | Holotype | n/a | USNM 8387 | n/a | n/a |
| Geodia ovis | Holotype | n/a | USNM 8388 | n/a | n/a |
| Penares anyapax | Paratype | TLT75 | TBD | TBD | TBD |
| Penares anyapax | Paratype | TLT76 | TBD | n/a | TBD |
| Penares anyapax | Paratype | TLT61 | TBD | TBD | TBD |
| Penares anyapax | Holotype | TLT1549 | TBD | n/a | TBD |
| Penares cf. saccharis | n/a | n/a | USNM 37916 | n/a | n/a |
| Penares cortius | n/a | n/a | USNM 31321 | n/a | n/a |
| Penares cortius | n/a | TLT67 | TBD | TBD | TBD |
| Penares cortius | n/a | TLT56 | TBD | TBD | TBD |
| Penares cortius | n/a | TLT118 | TBD | TBD | TBD |
| Penares cortius | n/a | n/a | NHMLA 15518 & BULA 227 | TBD | TBD |
| Penares cortius | n/a | n/a | NHMLA 15517 & BULA 226 | TBD | TBD |
| Penares cortius | n/a | TLT325 | TBD | TBD | TBD |
| Penares cortius | n/a | TLT437 | TBD | TBD | TBD |
| Penares cortius | n/a | TLT368 | TBD | TBD | TBD |
| Penares cortius | n/a | TLT468 | TBD | TBD | TBD |
| Penares cortius | n/a | TLT946 | TBD | TBD | TBD |
| Penares cortius | n/a | TLT1247 | TBD | TBD | TBD |
| Penares cortius | n/a | TLT1466 | TBD | TBD | TBD |
| Penares cortius | n/a | TLT1497 | TBD | TBD | TBD |
| Penares cortius | n/a | TLT1548 | TBD | n/a | n/a |
| Penares foxi | Holotype | TLT1455 B | TBD | TBD | TBD |
| Penares orientalis | n/a | n/a | CASIZ 302613 | TBD | TBD |
| Penares orientalis | n/a | TLT1616 | TBD | TBD | TBD |
| Penares orientalis | n/a | n/a | RBCM 018-00912-001 | TBD | TBD |
| Penares orientalis | n/a | n/a | CASIZ 233435 | n/a | n/a |
| Penares saccharis | Holotype | n/a | USNM 21476 | n/a | n/a |
| Penares saccharis | Paratype | n/a | USNM 21477 | n/a | n/a |
| Penares saccharis | n/a | n/a | CASIZ 450272 | TBD | n/a |
| Poecillastra alaskensis | Holotype | AB16-0156 | USNM 1512719 | n/a | n/a |
| Poecillastra alaskensis | Paratype | EX2306-D08-02B | USNM 1699915 | n/a | TBD |
| Poecillastra rickettsi | n/a | n/a | USNM 1479000 | n/a | n/a |
| Poecillastra rickettsi | n/a | n/a | USNM 1479051 | n/a | n/a |
| Poecillastra rickettsi | n/a | n/a | USNM 1478681 | n/a | TBD |
| Poecillastra rickettsi | n/a | n/a | RBCM 020-00056-002 | TBD | TBD |
| Poecillastra rickettsi | n/a | n/a | RBCM 018-00148-007 | TBD | TBD |
| Poecillastra rickettsi | n/a | n/a | RBCM 010-00247-002 | TBD | TBD |
| Poecillastra rickettsi | n/a | n/a | RBCM 988-00002-095 | TBD | TBD |
| Poecillastra rickettsi | n/a | n/a | RBCM 991-00332-069 | TBD | n/a |
| Poecillastra rickettsi | n/a | n/a | RBCM 999-00213-004 | TBD | n/a |
| Poecillastra rickettsi | n/a | PAC2024-041 | RBCM 025-00144-001 | TBD | TBD |
| Poecillastra rickettsi | n/a | n/a | RBCM 014-00176-006 | n/a | n/a |
| Poecillastra rickettsi | Holotype | n/a | USNM 21482 | n/a | n/a |
| Poecillastra rickettsi | n/a | n/a | CASIZ 179739 | TBD | TBD |
| Poecillastra rickettsi | n/a | n/a | CASIZ 165832 | TBD | n/a |
| Poecillastra rickettsi | n/a | n/a | CASIZ 180060 | TBD | n/a |
| Poecillastra rickettsi | n/a | n/a | USNM 21400 | n/a | n/a |
| Poecillastra rickettsi | n/a | 1987-3456 | SBMNH 467987 | n/a | n/a |
| Poecillastra rickettsi | n/a | n/a | CASIZ 235709 | n/a | n/a |
| Poecillastra rickettsi | n/a | BFHL 1225 | RBCM 018-00210-001 | TBD | TBD |
| Stelletta cardenasi | Holotype | TLT352 | NHMLA 16524 & TBD | TBD | TBD |
| Stelletta cardenasi | Paratype | TLT521 | TBD | TBD | n/a |
| Stelletta cardenasi | n/a | TLT271B | TBD | TBD | n/a |
| Stelletta cardenasi | n/a | TLT270 | TBD | TBD | n/a |
| Stelletta cardenasi | n/a | TLT776A | TBD | TBD | n/a |
| Stelletta cardenasi | n/a | TLT391 | TBD | TBD | n/a |
| Stelletta cardenasi | n/a | TLT1459 B | TBD | TBD | n/a |
| Stelletta cardenasi | n/a | n/a | CASIZ 302848 | TBD | n/a |
| Stelletta cardenasi | n/a | TLT786B | TBD | TBD | n/a |
| Stelletta clarella | n/a | n/a | CASIZ 302625 | TBD | TBD |
| Stelletta clarella | n/a | TLT1555 | TBD | TBD | TBD |
| Stelletta clarella | n/a | TLT1576 | TBD | TBD | TBD |
| Stelletta clarella | n/a | TLT1580 | TBD | TBD | TBD |
| Stelletta clarella | n/a | TLT1676 | TBD | TBD | TBD |
| Stelletta clarella | n/a | TLT1677 | TBD | TBD | TBD |
| Stelletta clarella | n/a | TLT459 | TBD | TBD | TBD |
| Stelletta clarella | n/a | TLT569 | TBD | TBD | TBD |
| Stelletta clarella | n/a | TLT1157 | TBD | TBD | TBD |
| Stelletta clarella | n/a | TLT1165 | TBD | TBD | TBD |
| Stelletta clarella | n/a | TLT1139 | UCSB-IZC00048471 | TBD | TBD |
| Stelletta clarella | n/a | TLT520 | TBD | TBD | TBD |
| Stelletta estrella | Holotype | n/a | USNM 21399 | n/a | n/a |
| Stelletta estrella | n/a | TLT286 | NHMLA 16544 | TBD | TBD |
| Stelletta estrella | n/a | TLT1343 | TBD | TBD | TBD |
| Stelletta estrella | n/a | TLT466 | TBD | TBD | TBD |
| Stelletta limuwensis | Holotype | TLT329 | TBD | n/a | TBD |
| Stelletta makushina | n/a | n/a | RBCM 019-00567-006 | TBD | TBD |
| Stelletta nicolenya | Holotype | TLT1492 B | TBD | TBD | TBD |
| Stelletta ovalae | n/a | n/a | CASIZ 302615 | TBD | TBD |
| Stelletta ovalae | n/a | n/a | CASIZ 302624 | TBD | TBD |
| Stelletta ovalae | n/a | TLT1674 | TBD | TBD | TBD |
| Stelletta ovalae | n/a | TLT1562 | TBD | TBD | TBD |
| Stelletta ovalae | n/a | n/a | RBCM 018-00914-001 | TBD | TBD |
| Thenia diastra | Holotype | n/a | RBCM 020-00056-001 | n/a | TBD |
| Vulcanella explorata | Holotype | n/a | RBCM 018-00975-006 | DSEP047-25.COI-5P | TBD |
| Vulcanella explorata | Paratype | n/a | RBCM 018-00975-005 | TBD & DSEP048-25.COI-5P | TBD |
| Vulcanella rupta | Holotype | n/a | RBCM 018-00898-006 | TBD | TBD |
| Vulcanella rupta | n/a | n/a | RBCM 018-00879-002 | TBD | TBD |
| Vulcanella rupta | n/a | n/a | RBCM 010-00538-001 | TBD | TBD |
| Vulcanella rupta | Paratype | n/a | RBCM 018-00898-002 | TBD | TBD |
| Vulcanella rupta | n/a | n/a | RBCM 023-00023-003 | TBD | TBD |
| Vulcanella rupta | n/a | n/a | RBCM 018-00967-001 | TBD | TBD |

## References

Austin, W.C. (1985) An Annotated Checklist of Marine Invertebrates in the Cold Temperate Northeast Pacific. Volume 1. Khoyatan Marine Laboratory, Cowichan Bay, B.C., 43 pp.

Bakus, G.J. & Green, K.D. (1987) The distribution of marine sponges collected from the 1976-1978 Bureau of Land Management Southern California Bight program. Bulletin of the Southern California Academy of Sciences, 86 (2), 57–88.

Beckmann, L.M., Vad, J. & Waller, R.G. (2026) Diversity and environmental drivers of deep-sea sponge and coral communities in Alaska. Deep Sea Research Part I: Oceanographic Research Papers, 229. 10.1016/j.dsr.2026.104683.

Bowerbank, J.S. (1858) On the anatomy and physiology of the Spongiadae. Part I. On the spicula. Philosophical Transactions of the Royal Society, 148 (2), 279–332.

Bowerbank, J.S. (1866) A Monograph of the British Spongiadae. Volume 2. Ray Society, London, 388 pp. Available from: https://biodiversitylibrary.org/page/1905089

Burton, M. (1930) Norwegian sponges from the Norman Collection. Proceedings of the Zoological Society of London, 2, 487–546.

Burton, M. (1959) Sponges. In: Scientific Reports. John Murray Expedition 1933-34. British Museum (Natural History), London, pp. 151–415.

Calcinai, B., Bastari, A., Makapedua, D.M. & Cerrano, C. (2017) Mangrove sponges from Bangka Island (North Sulawesi, Indonesia) with the description of a new species. Journal of the Marine Biological Association of the United Kingdom, 97, 1417–1422. 10.1017/s0025315416000710

Carballo, J.L. & Vega, C. (2022) Esponjas (Porifera). In: Invertebrados marinos y costeros del Pacífico sur de México. Universidad del Mar y Geomare, Puerto Ángel, Oaxaca, México, pp. 21–30.

Cárdenas, P. (2020) Surface microornamentation of demosponge sterraster spicules, phylogenetic and paleontological implications. Frontiers in Marine Science, 7, 613610. 10.3389/fmars.2020.613610

Cárdenas, P. & Moore, J.A. (2019) First records of Geodia demosponges from the New England seamounts, an opportunity to test the use of DNA mini-barcodes on museum specimens. Marine Biodiversity, 49 (1), 163–174. 10.1007/s12526-017-0775-3

Cárdenas, P. & Rapp, H.T. (2012) A review of Norwegian streptaster-bearing Astrophorida (Porifera: Demospon- giae: Tetractinellida), new records and a new species. Zootaxa, 3253, 1–52. 10.11646/zootaxa.3253.1.1

Cárdenas, P., Rapp, H.T., Schander, C. & Tendal, O.S. (2010) Molecular taxonomy and phylogeny of the Geodiidae (Porifera, Demospongiae, Astrophorida) – combining phylogenetic and Linnaean classification. Zoologica Scripta, 39 (1), 89–106. 10.1111/j.1463-6409.2009.00402.x

Cárdenas, P., Xavier, J.R., Reveillaud, J., Schander, C. & Rapp, H.T. (2011) Molecular phylogeny of the Astrophorida (Porifera, Demospongiae(p)) reveals an unexpected high level of spicule homoplasy. PloS One, 6 (4), e18318. 10.1371/journal.pone.0018318

Carter, H.J. (1883) Contributions to our knowledge of the Spongida. Pachytragida. Annals and Magazine of Natural History, 5 (11), 344–369. 10.1080/00222938309459163

Carvalho, M.S., Desqueyroux-Faúndez, R. & Hajdu, E. (2011) Taxonomic notes on Poecillastra sponges (Astrophorida: Pachastrellidae), with the description of three new bathyal southeastern Pacific species. Scientia Marina, 75 (3). 10.3989/scimar.2011.75n3477

Chen, S., Zhou, Y., Chen, Y. & Gu, J. (2018) fastp: an ultra-fast all-in-one FASTQ preprocessor. Bioinformatics, 34 (17), i884–i890. 10.1093/bioinformatics/bty560

Danecek, P., Bonfield, J.K., Liddle, J., Marshall, J., Ohan, V., Pollard, M.O., Whitwham, A., Keane, T., McCarthy, S.A., Davies, R.M. & Li, H. (2021) Twelve years of SAMtools and BCFtools. GigaScience, 10 (2), giab008. 10.1093/gigascience/giab008

Dendy, A. & Burton, M. (1926) Report on some deep-sea sponges from the Indian Museum collected by the R.I.M.S. ‘Investigator’. Part I. Hexactinellida and Tetractinellida (pars). Records of the Indian Museum, 28 (4), 225–248. 10.26515/rzsi/v28/i4/1926/163228

Despujols, D. (2021) Integrative taxonomy of the genus Thenea (Porifera, Demospongiae, Tetractinellida) of the Portuguese shelf and slope: new records and new species for science. Masters. University of Porto.

Desqueyroux-Faúndez, R. & van Soest, R.W.M. (1997) Shallow-water Demosponges of the Galapagos Islands. Revue suisse de Zoologie, 104 (2), 379–467.

Díaz, J.A., Ordines, F., Massutí, E. & Cárdenas, P. (2024) From caves to seamounts: the hidden diversity of tetractinellid sponges from the Balearic Islands, with the description of eight new species. PeerJ, 12 (e16584).

Dickinson, M.G. (1945) Sponges of the Gulf of California. In: Reports on the collections obtained by Allan Hancock Pacific Expeditions of Velero III off the coast of Mexico, Central America, South America, and Galapagos Islands in 1932, in 1933, in 1934, in 1935, in 1936, in 1937, in 1939, and 1940. Vol.11 The University of Southern California Press, Los Angeles, pp. 1–55.

Erpenbeck, D., Ekins, M., Enghuber, N., Hooper, J.N.A., Lehnert, H., Poliseno, A., Schuster, A., Setiawan, E., de Voogd, N.J., Worheide, G. & van Soest, R.W.M. (2015) Nothing in (sponge) biology makes sense – except when based on holotypes. Journal of the Marine Biological Association of the United Kingdom, 96 (2), 305–311.

Folmer, O., Black, M., Wr, H. & Vrijenhoek, R. (1994) DNA primers for amplification of mitochondrial Cytochrome C oxidase subunit I from diverse metazoan invertebrates. Molecular Marine Biology and Biotechnology, 3, 294–299.

Gómez, P., Carballo, J.L., Vázquez, L.E. & Cruz, J.A. (2002) New records for the sponge fauna (Porifera: Demospongiae) of the Pacific coast of Mexico (eastern Pacific Ocean). Proceedings of the Biological Society of Washington, 115 (1), 223–237.

Graiff, K. & Lipski, D. (2023) Characterization of Cordell Bank and continental shelf and slope: 2021 ROV surveys. National Marine Sanctuaries Conservation Series, National Oceanic and Atmospheric Administration.

Green, K.D. & Bakus, G.J. (1994) Taxonomic atlas of the benthic marine fauna of the Western Santa Maria Basin and the Western Santa Barbara Channel. Volume 2: The Porifera. The Santa Barbara Museum of Natural History, 82 pp.

Hoang, D., Chernomor, O., von Haeseler, A., Minh, B. & Vinh, L. (2018) UFBoot2: Improving the ultrafast bootstrap approximation. Molecular Biology and Evolution, 35, 518–522. 10.1093/molbev/msx281

Hooper, J.N.A. & van Soest, R. (2002) Order Astrophorida Sollas, 1888. In: Systema Porifera: A Guide to the Classification of Sponges. Kluwer Academic, Plenum Publishers, New York, pp. 105–107.

Huang, D., Meier, R., Todd, P.A. & Chou, L.M. (2008) Slow mitochondrial COI sequence evolution at the base of the Metazoan tree and its implications for DNA barcoding. Journal of Molecular Evolution, 66, 167–174. 10.1007/s00239-008-9069-5

Jennings, H.S. (1917) The numerical results of diverse systems of breeding, with respect to two pairs of characters, linked or independent, with special relation to the effects of linkage. Genetics, 2 (2), 97–154.

Kalyaanamoorthy, S., Minh, B.Q., Wong, T.K.F., von Haeseler, A. & Jermiin, L.S. (2017) ModelFinder: fast model selection for accurate phylogenetic estimates. Nature Methods, 14 (6), 587–589. 10.1038/nmeth.4285

Katoh, K., Rozewicki, J. & Yamada, K.D. (2017) MAFFT online service: multiple sequence alignment, interactive sequence choice and visualization. Briefings in Bioinformatics, 20 (4), 1160–1166. 10.1093/bib/bbx108

Kelly, M., Cárdenas, P., Rush, N., Sim-Smith, C., Macpherson, D., Page, M. & Bell, L.J. (2019) Molecular study supports the position of the New Zealand endemic genus Lamellomorpha in the family Vulcanellidae (Porifera, Demospongiae, Tetractinellida), with the description of three new species. European Journal of Taxonomy, 506. 10.5852/ejt.2019.506

Klitgaard, A.B. & Tendal, O.S. (2004) Distribution and species composition of mass occurrences of large-sized sponges in the northeast Atlantic. Progress in Oceanography, 61, 57–98. 10.1016/j.pocean.2004.06.002

Koltun, V.M. (1964) Sponges of the Antarctic. 1 Tetraxonida and Cornacuspongida. In: Biological Reports of the Soviet Antarctic Expedition (1955-1958). Akademya Nauk SSSR, pp. 6–133, 443–448.

Koltun, V.M. (1966) Four-rayed sponges of Northern and Far Eastern seas of the USSR (order Tetraxonida. Opredeliti Faunei SSSR, 90, 1–112.

Lambe, L.M. (1895) Sponges from the Western Coast of North America. Transactions of the Royal Society of Canada, 12 (4), 113–138.

Lampe, W. (1886) Tetilla japonica, eine neue Tetractinellidenform mit radiärem Bau. Archiv für Naturgeschichte, 52 (1).

de Laubenfels, M.W. (1930) The Sponges of California. Stanford University Bulletin, 5 (98), 24–29.

de Laubenfels, M.W. (1932) The marine and fresh-water sponges of California. Proceedings of the United States National Museum, 81 (2927), 1–140.

Lavrov, D.V., Diaz, M.C., Maldonado, M., Morrow, C.C., Pérez, T., Pomponi, S.A. & Thacker, R.W. (2023) Phylomitogenomics bolsters the high-level classification of Demospongiae (phylum Porifera). PLoS ONE, 18 (12), e0287281. 10.1371/journal.pone.0287281

Lavrov, D.V., Wang, X. & Kelly, M. (2008) Reconstructing ordinal relationships in the Demospongiae using mitochondrial genomic data. Molecular Phylogenetics and Evolution, 49 (1), 111–124.

Lebwohl, F. (1914) Japanische Tetraxonida, I. Sigmatophora und II. Astrophora metastrosa. *Journal of the College of Sciences*, Imperial University of Tokyo, 35 (2).

Lee, W., Elvin, D. & Reiswig, H. (2007) The sponges of California: A guide and key to the marine sponges of California. Monterey Bay Sanctuary Foundation, Monterey, CA, 395 pp.

Lehnert, H., Stone, R. & Heimler, W. (2006) Erylus aleuticus sp.nov. (Porifera: Demospongiae: Astrophorida: Geodiidae) from the Aleutian Islands, Alaska, USA. Journal of the Marine Biological Association of the United Kingdom, 86, 971–975. 10.1017/s0025315406013944

Lehnert, H. & Stone, R.P. (2014) Aleutian Ancorinidae (Porifera, Astrophorida): Description of three new species from the genera Stelletta and Ancorina. Zootaxa, 3826, 341–355. 10.11646/zootaxa.3826.2.4

Lehnert, H. & Stone, R.P. (2016) A comprehensive inventory of the Gulf of Alaska sponge fauna with the description of two new species and geographic range extensions. Zootaxa, 4144 (3), 365–382. 10.11646/zootaxa.4144.3.5

Lehnert, H., Stone, R.P. & Drumm, D. (2014) Geodia starki sp. nov. (Porifera, Demospongiae, Astrophorida) from the Aleutian Islands, Alaska, USA. Journal of the Marine Biological Association of the United Kingdom, 94 (2), 261–265. 10.1017/S002531541300101X

Lendenfeld, R. von (1907) Die Tetraxonia. Wissenschaftliche Ergebnisse der Deutschen Tiefsee-Expedition auf der Dampfer Valdivia 1898-1899, 11 (1–2), 59–374.

Lendenfeld, R. von. (1910) The Sponges. 1. The Geodidae. In: Reports on the Scientific Results of the Expedition to the Eastern Tropical Pacific, in charge of Alexander Agassiz, by the U.S. Fish Commission Steamer ‘Albatross’, from October, 1904, to March, 1905, Lieut. Commander L.M. Garrett, U.S.N., Commanding, and of other Expeditions of the Albatross, 1888-1904. Memoirs of the Museum of Comparative Zoology at Harvard College, 41 (1), 1–259.

Li, D., Liu, C.-M., Luo, R., Sadakane, K. & Lam, T.-W. (2015) MEGAHIT: an ultra-fast single-node solution for large and complex metagenomics assembly via succinct de Bruijn graph. Bioinformatics, 31 (10), 1674–1676. 10.1093/bioinformatics/btv033

Li, H. & Durbin, R. (2009) Fast and accurate short read alignment with Burrows-Wheeler transform. *Bioinformatics (Oxford*, England), 25 (14), 1754–1760. 10.1093/bioinformatics/btp324

Lim, S.-C., Wiklund, H., Glover, A.G., Dahlgren, T.G. & Tan, K.-S. (2017) A new genus and species of abyssal sponge commonly encrusting polymetallic nodules in the Clarion-Clipperton Zone, East Pacific Ocean. Systematics and Biodiversity, 15 (6), 507–519.

Lindholm, J., Cramer, A.N. & Braddock, A.M. (2013) Distribution and Abundance of Selected Corals and Sponges in the Channel Islands National Marine Sanctuary as Determined from ROV Video Imagery. National Oceanic and Atmospheric Administration Institutional Repository.

López-Legentil, S., Erwin, P.M., Henkel, T.P., Loh, T.L. & Pawlik, J.R. (2010) Phenotypic plasticity in the Caribbean sponge Callyspongia vaginalis (Porifera: Haplosclerida). Scientia Marina, 74 (3), 445–453. 10.3989/scimar.2010.74n3445

Maldonado, M. (2002) Family Pachastrellidae Carter, 1875. In: Systema Porifera. A Guide to the Classification of Sponges. Kluwer Academic/Plenum Publ., New York.

Maldonado, M., Aguilar, R., Blanco, J., García, S., Serrano, A. & Punzón, A. (2015) Aggregated clumps of lithistid aponges: a singular, reef-Like bathyal habitat with relevant paleontological connections. PLoS ONE, e0125378. 10.1371/journal.pone.0125378

Morrow, C.C., Picton, B.E., Erpenbeck, D., Boury-Esnault, N., Maggs, C.A. & Allcock, A.L. (2012) Congruence between nuclear and mitochondrial genes in Demospongiae: A new hypothesis for relationships within the G4 clade (Porifera: Demospongiae). Molecular Phylogenetics and Evolution, 62, 174–190. 10.1016/j.ympev.2011.09.016

Nguyen, L.-T., Schmidt, H., von Haeseler, A. & Minh, B. (2015) IQ-TREE: A fast and effective stochastic algorithm for estimating maximum likelihood phylogenies. Molecular Biology and Evolution, 32, 268–274. 10.1093/molbev/msu300

Ngwakum, B., Payne, R.P., Teske, P.R., Janson, L., Kerwath, S.E. & Samaai, T. (2021) Hundreds of new DNA barcodes for South African sponges. Systematics and Biodiversity, 19 (7), 747–769. 10.1080/14772000.2021.1915896

Nichols, S. (2005) An evaluation of support for order-level monophyly and interrelationships within the class Demospongiae using partial data from the large subunit rDNA and cytochrome oxidase subunit I. Molecular Phylogenetics and Evolution, 34, 81–96. 10.1016/j.ympev.2004.08.019

Nunley, R.M., Rutkowski, E.C., Toonen, R.J. & Vicente, J. (2025) Potential transoceanic dispersal of Geodia cf. papyracea and six new tetractinellid sponge species descriptions within the Hawaiian reef cryptofauna. PeerJ, 13 (e18903). 10.7717/peerj.18903

Plese, B., Kenny, N.J., Rossi, M.E., Cárdenas, P., Schuster, A., Taboada, S., Koutsouveli, V. & Riesgo, A. (2021) Mitochondrial evolution in the Demospongiae (Porifera): Phylogeny, divergence time, and genome biology. Molecular Phylogenetics and Evolution, Feb:155:107011. 10.1016/j.ympev.2020.107011

Pöppe, J., Sutcliffe, P., Hooper, J.N., Wörheide, G. & Erpenbeck, D. (2010) CO1 barcoding reveals new clades and radiation patterns of Indo-Pacific sponges of the family Irciniidae. PLoS ONE, 5 (4), e9950. 10.1371/journal.pone.0009950

Rannala, B. & Yang, Z. (2020) Chapter 5.5 Species Delimitation. In: Scornavacca, C., Delsuc, F., & Galtier, N. (Eds), Phylogenetics in the Genomic Era>. No commercial publisher, pp. 1–568.

Rho, B.-J. & Sim, C.-J. (1981) A taxonomic study on the sponges in Korea 3. Choristida. Journal of the Korean Research Institute of the Living Arts, 28.

Rossi, M.E., Keating, J.N., Kenny, N.J., Giacomelli, M., Álvarez-Carretero, S., Schuster, A., Cárdenas, P., Taboada, S., Koutsouveli, V., Donoghue, P.C.J., Riesgo, A. & Pisani, D. (2026) Independent origins of spicules reconcile paleontological and molecular evidence of sponge evolutionary history. Science Advances, 12, eadx1754. 10.1126/sciadv.adx1754

Rot, C., Goldfarb, I., Ilan, M. & Huchon, D. (2006) Putative cross-kingdom horizontal gene transfer in sponge (Porifera) mitochondria. BMC Evolutionary Biology, 6 (71), 1–11. 10.1186/1471-2148-6-71

Santos, G.G. & Pinheiro, U. (2016) A new species of Dercitus (Stoeba) from the Atlantic Ocean (Porifera: Demospongiae: Astrophorida). Journal of the Marine Biological Association of the United Kingdom, 96 (3), 681–686. 10.1017/S0025315414002100

Schmidt, O. (1862) Die Spongien des adriatischen Meeres. Wilhelm Engelmann, Leipzig, 88 pp.

Schmidt, O. (1864) Supplement der Spongien des adriatischen Meeres. In: Enthaltend die Histologie und systematische Ergänzungen., pp. 1–48.

Schmidt, O. (1868) Die Spongien der Küste von Algier. Mit Nachträgen zu den Spongien des Adriatischen Meeres (Drittes Supplement). Wilhelm Engelmann, Leipzig, 44 pp.

Schneider, C., Rasband, W. & Eliceiri, K. (2012) NIH Image to ImageJ: 25 years of image analysis. Nature Methods, 9 (7), 671–675. 10.1038/nmeth.2089

Schuster, A., Cárdenas, P., Pisera, A., Pomponi, S.A., Kelly, M., Wörheide, G. & Erpenbeck, D. (2018) Seven new deep-water Tetractinellida (Porifera: Demospongiae) from the Galápagos Islands – morphological descriptions and DNA barcodes. Zoological Journal of the Linnean Society, 184 (2), 273–303. 10.1093/zoolinnean/zlx110

Schuster, A., Erpenbeck, D., Pisera, A., Hooper, J., Bryce, M., Fromont, J. & Wörheide, G. (2015) Deceptive desmas: Molecular phylogenetics suggests a new classification and uncovers convergent evolution of lithistid Demosponges. Plos One, 10 (1), e116038.

Shilov, V., Kamenev, Y.O., Semenchenko, A.A., Kiyashkoa, S.I. & Mordukhovich, V.V. (2023) New sponge species of the family Vulcanellidae (Demospongiae: Tetractinellida) from the Piip submarine volcano and adjacent areas (Bering Sea, NW Pacific). Deep-Sea Research Part II, 105229, 1–23. 10.1016/j.dsr2.2022.105229

Sim-Smith, C., Cleveland Hickman, J. & Kelly, M. (2021) New shallow-water sponges (Porifera) from the Galápagos Islands. Zootaxa, 5012 (1), 1–71. 10.11646/zootaxa.5012.1.1

Sim-Smith, C. & Kelly, M. (2019) Review of the sponge genus Penares (Demospongiae, Tetractinellida, Astrophorina) in the New Zealand EEZ, with descriptions of new species. Zootaxa, 4638 (1), 001–056. 10.11646/zootaxa.4638.1.1

van Soest, R.W.M., Hooper, J.N.A. & Butler, P.J. (2020) Every sponge its own name: removing Porifera homonyms. Zootaxa, 4745 (1), 1–93. 10.11646/zootaxa.4745.1.1

Sollas, W.J. (1886) Preliminary account of the Tetractinellid sponges dredged by H.M.S. ‘Challenger’ 1872–76. Part I. The Choristida. Scientific Proceedings of the Royal Dublin Society, 5, 177–199.

Sollas, W.J. (1888) Report on the Tetractinellida collected by H.M.S. Challenger, during the years 1873–1876. Report on the Scientific Results of the Voyage of H.M.S. Challenger during the years 1873–76. Zoology, 25 (Part 63), 1–458.

Spalding, M.D., Fox, H.E., Allen, G.R., Davidson, N., Ferdaña, Z.A., Finlayson, M., Halpern, B.S., Jorge, M.A., Lombana, A., Lourie, S.A., Martin, K.D., McManus, E., Molnar, J., Recchia, C.A. & Robertson, J. (2007) Marine ecoregions of the world: A bioregionalization of coastal and shelf areas. BioScience, 57 (7), 573–583. 10.1641/B570707

Steffen, K., Proux-Wéra, E., Soler, L., Churcher, A., Sundh, J. & Cárdenas, P. (2023) Whole genome sequence of the deep-sea sponge Geodia barretti (Metazoa, Porifera, Demospongiae). G3 Genes|Genomes|Genetics, 13 (10), jkad192. 10.1093/g3journal/jkad192

Stone, R.P. (2014) The ecology of deep-sea coral and sponge habitats of the central Aleutian Islands of Alaska. National Marine Fisheries Service, NOAA. NOAA Professional Paper NMFS.

Stone, R.P., Lehnert, H. & Reiswig, H.M. (2011) A guide to the deepwater sponges of the Aleutian Island Archipelago. NOAA Professional Paper NMFS, 12, 1–187.

Tanita, S. (1965) Report on the sponges obtained from the adjacent waters of the Sado Island, Japan Sea. Bulletin Japan Sea Regional Fisheries Research Laboratory, 14, 43–66.

Tanita, S. & Hoshino, T. (1989) The Demospongiae of Sagami Bay. Biological Laboratory, Imperial Household: Japan, 197 pp.

Thacker, R., Hill, A., Hill, M., Redmond, N., Collins, A., Morrow, C., Spicer, L., Carmack, C., Zappe, M., Pohlmann, D., Hall, C., Diaz, M. & Bangalore, P. (2013) Nearly complete 28S rRNA gene sequences confirm new hypotheses of sponge evolution. Integrative and Comparative Biology, 53, 373–387. 10.1093/icb/ict071

Thiele, J. (1898) Studien über pazifische Spongien. I. Japanische Demospongien. Zoologica. Original-Abhandlungen aus dem Gesamtgebiete der Zoologie. Stuttgart., 24 (1), 1–72.

Topsent, E. (1893) Nouvelle série de diagnoses d’éponges de Roscoff et de Banyuls. Archives de Zoologie expérimentale et générale, 3 (1).

Trifinopoulos, J., Nguyen, L.-T., von Haeseler, A. & Minh, B. (2016) W-IQ-TREE: a fast online phylogenetic tool for maximum likelihood analysis. Nucleic Acids Research, 44, W232–W235. 10.1093/nar/gkw256

Turner, C.H., Ebert, E.E. & Given, R.R. (1965) Survey of the marine environment offshore of San Elijo Lagoon, San Diego County. Cali. Fish Game, 51, 81–112.

Turner, T.L. (2020) The order Tethyida (Porifera) in California: taxonomy, systematics, and the first member of the family Hemiasterellidae in the Eastern Pacific. Zootaxa, 4861 (2), 211–231. 10.11646/zootaxa.4861.2.3

Turner, T.L. (2026) Systematic revision of the family Tetillidae (Porifera: Demospongiae) in the temperate Northeast Pacific. Zoological Journal of the Linnean Society, in press. 10.1101/2025.09.30.679634

Turner, T.L., Morrow, C., Picton, B., Goodwin, C. & Thacker, R.W. (2025) A common garden of Halichondria sponges: taxonomic revision of Northeast Pacific Halichondriidae reveals many cryptic introduced species. Bulletin of the Society of Systematic Biologists, 1–56. 10.18061/bssb.5811

Turner, T.L. & Pankey, M.S. (2023) The order Axinellida (Porifera: Demospongiae) in California. Zootaxa, 5230 (5), 501–539. 10.11646/zootaxa.5230.5.1

Turner, T.L., Rouse, G.W., Weigel, B.L., Janusson, C., Lemay, M.A. & Thacker, R.W. (2024) Taxonomy and phylogeny of the family Suberitidae (Porifera: Demospongiae) in California. Zootaxa, 5447 (1), 1–28. 10.11646/zootaxa.5447.1.1

Uriz, M.J. (2002a) Family Ancorinidae Schmidt, 1870. *In*: Hooper, J. N. A. & van Soest, R. W. M. (Eds), Systema Porifera. A guide to the classification of sponges. Kluwer Academic/Plenum Publ., New York, pp. 108–126.

Uriz, M.J. (2002b) Family Geodiidae Gray, 1867. *In*: Hooper, J. N. A. & van Soest, R. W. M. (Eds), Systema Porifera. A guide to the classification of sponges. Kluwer Academic/Plenum Publ., New York, pp. 134–140.

Van Soest, R.W.M., Beglinger, E.J. & de Voogd, N.J. (2010) Skeletons in confusion: a review of astrophorid sponges with (dicho–)calthrops as structural megascleres (Porifera, Demospongiae,Astrophorida). Zookeys, 68, 1–88. 10.3897/zookeys.68.729

de Voogd, N.J., Alvarez, B., Boury-Esnault, N., Cárdenas, P., Díaz, C., Dohrmann, M., Downey, R., Goodwin, C., Hajdu, E., Hooper, J.N.A., Kelly, M., Klautau, M., Lim, S.C., Monconi, R., Morrow, C.C., Pinheiro, U., Pisera, A.B., Ríos, P., Rützler, K., Schönberg, C., Turner, T.L., Vacelet, J., van Soest, R.W.M. & Xavier, J. (2025) World Porifera Database. Available from: https://www.marinespecies.org/porifera on 2025-12-05. doi:10.14284/359 (accessed 5 September 2025)

Watson, J.R., Hays, C.G., Raimondi, P.T., Mitarai, S., Dong, C., McWilliams, J.C., Blanchette, C.A., Caselle, J.E. & Siegel, D.A. (2011) Currents connecting communities: nearshore community similarity and ocean circulation. Ecology, 92 (6), 1193–1200. 10.1890/10-1436.1

Williams, S.E., Varliero, G., Lurgi, M., Stach, J.E.M., Race, P.R. & Curnow, P. (2024) Diversity and structure of the deep-sea sponge microbiome in the equatorial Atlantic Ocean. Microbiology, 170 (7), 001478. 10.1099/mic.0.001478

Wilson, H.V. (1904) Reports on an exploration off the West Coasts of Mexico, Central and South America, and off the Galapagos Islands, in charge of Alexander Agassiz, by the U.S. Fish Commission Steamer ‘Albatross’ during 1891, Lieut. Commander Z.L. Tanner, U.S.S., commanding. XXX. The Sponges. Memoirs of the Museum of Comparative Zoology at Harvard College, 30 (1), 1–164.

Zeng, C., Thomas, L.J., Kelly, M. & Gardner, J.P.A. (2016) The complete mitochondrial genome of the deep-sea sponge Poecillastra laminaris (Astrophorida, Vulcanellidae). Mitochondrial DNA A DNA Mapp Seq Anal., 27 (3), 1658–1659. 10.3109/19401736.2014.958716

